# Deep topic modeling of spatial transcriptomics in the rheumatoid arthritis synovium identifies distinct classes of ectopic lymphoid structures

**DOI:** 10.1101/2025.01.08.631928

**Authors:** Preethi K. Periyakoil, Melanie H. Smith, Meghana Kshirsagar, Daniel Ramirez, Edward F. DiCarlo, Susan M. Goodman, Alexander Y. Rudensky, Laura T. Donlin, Christina S. Leslie

**Author notes:** These authors contributed equally to this work (Preethi K. Periyakoil, Melanie H. Smith).

## Abstract

Single-cell RNA sequencing studies have revealed the heterogeneity of cell states present in the rheumatoid arthritis (RA) synovium. However, it remains unclear how these cell types interact with one another *in situ* and how synovial microenvironments shape observed cell states. Here, we use spatial transcriptomics (ST) to define stable microenvironments across eight synovial tissue samples from six RA patients and characterize the cellular composition of ectopic lymphoid structures (ELS). To identify disease-relevant cellular communities, we developed DeepTopics, a scalable reference-free deconvolution method based on a Dirichlet variational autoencoder architecture. DeepTopics identified 22 topics across tissue samples that were defined by specific cell types, activation states, and/or biological processes. Some topics were defined by multiple colocalizing cell types, such as CD34^+^ fibroblasts and LYVE1^+^ macrophages, suggesting functional interactions. Within ELS, we discovered two divergent cellular patterns that were stable across ELS in each patient and typified by the presence or absence of a “germinal-center-like” topic. DeepTopics is a versatile and computationally efficient method for identifying disease-relevant microenvironments from ST data, and our results highlight divergent cellular architectures in histologically similar RA synovial samples that have implications for disease pathogenesis.

## Introduction

Rheumatoid arthritis (RA) is a systemic autoimmune disease primarily characterized by joint inflammation that, if left untreated, can lead to permanent joint damage. In RA, inflammation of the synovium (joint lining) is classically characterized by an influx of leukocytes and an expansion of resident stromal cells. Infiltrating lymphocytes can form ectopic lymphoid structures (ELS), also called lymphocyte aggregates or tertiary lymphoid follicles, that are present in both early and late stages of disease^1, 2, 3^. In some patients, ELS can form functional germinal centers resulting in local autoantibody production^4, 2, 3^. Patients with a lympho-myeloid synovial pathotype, who also have tissue ELS, have the highest levels of serum inflammation markers (erythrocyte sedimentation rate (ESR) and C-reactive protein), anti-cyclic citrullinated peptide (CCP) autoantibodies, and disease activity scores (swollen joint counts and disease activity score-28 (DAS28)-ESR)^4, 5^. It remains unclear how the synovial microenvironment supports ELS formation and what causes some, but not all, ELS to form germinal-center-like structures.

Recent advances in single-cell sequencing have facilitated the cataloguing of cell types and states observed in the RA synovium resulting in unprecedented insight into potential pathogenic mechanisms^6^. Now, the challenge is to better define how these diverse cell types are organized into interacting communities with distinct microenvironments that perpetuate ongoing pathogenic inflammation. With the development of spatial transcriptomics (ST) technologies, we can now study individual cell subtypes and states within both the overall tissue architecture and within individual microenvironments^7, 8^.

A technical limitation of most sequencing-based ST platforms is that transcripts are associated to spatial capture regions and not to individual cells. Though technologies such as 10x Visium HD technology^9^ or Slide-seq V2^10^ have 2-10 µm capture regions (as opposed to the 55 µm capture region in the more widely-used 10x Visium platform^11^), each capture region can still encompass multiple cells or partial cells; this uncertainty is due to the fact that transcripts are barcoded based on their physical position in a fixed array, not based on individual cells.

Analysis typically requires a computational deconvolution step to determine which cell types are represented in the observed gene expression data in a particular capture region. Many current algorithms that deconvolve ST data, such as MuSIC^12^, DestVI^13^, or Tangram^14^, rely on the availability of a single-cell RNA-sequencing (scRNA-seq) reference atlas. Reference-based deconvolution approaches have several inherent limitations. First, relevant high-quality scRNA-seq reference atlases are not always available for a given tissue or disease state. Second, the differing capture biases between scRNA-seq and ST technologies may lead to an incompatible representation of cell types and cell-type-specific gene markers, resulting in incorrect or incomplete annotations. Relatedly, the fine-grained cell types that can be resolved in scRNA-seq may not be identifiable in ST, potentially confounding atlas-based deconvolution approaches. Unsupervised machine learning methods offer a solution to these problems, and previous studies have shown that topic modeling approaches such as latent Dirichlet allocation (LDA)^15, 16^ or non-negative matrix factorization (NMF)^17, 18^ enable ST spot deconvolution without requiring a reference scRNA-seq atlas. However, existing topic model methods are typically computationally expensive and do not scale as the number of input examples (capture regions) increases. As ST datasets continue to grow and improve in resolution, there is a need for reference-free deconvolution methods that are scalable and computationally efficient.

To address this issue, we adapted the Dirichlet variational autoencoder (VAE)^19^ model we introduced in Kshirsagar et al.^20^ to develop DeepTopics for scalable reference-free ST deconvolution. Standard autoencoders are neural networks with an encoder-decoder architecture, where the encoder learns to map input examples to a lower dimensional latent space, and the decoder attempts to reconstruct the original data from this latent representation, thus encouraging the latent features to be informative. VAEs are unsupervised deep learning models that incorporate probabilistic modeling concepts into encoder-decoder based neural networks. As a Dirichlet VAE, DeepTopics is a deep learning implementation of topic modeling that embeds the input data into a lower-dimensional space indexed by “topics”, where each topic is a distribution over genes that may correspond to the expression signature of a single cell type or other coordinated expression patterns in ST capture regions. This embedding process is probabilistic: the encoder neural network transforms an input example 𝒙, representing the gene-level count data from an ST region, into the parameters of a Dirichlet distribution. The VAE then samples from this distribution to produce the topic weight vector 𝒛, encoding the relative strength of different topics in the input example. Topics are learned during training and stored in the decoder network of DeepTopics, which takes the latent representation 𝒛 and the topic parameters and reconstructs the transcript data as 𝒙’. The training of the model involves learning the encoder and decoder (including topic) parameters such that for each input 𝒙, the reconstruction loss between the output 𝒙’ and 𝒙 is minimized.

Probabilistic models that are trained using traditional inference methods such as Gibbs sampling^21^ or stochastic variational inference^13^ are computationally inefficient and do not scale well to the large and high-dimensional training sets inherent in ST data. In VAEs, the inference step is done by the encoder neural network, and training relies on backpropagation-based gradient computation, both of which are very efficient. Previous work in single-cell transcriptomic analysis has used VAEs, including scVI^22^, a neural network-based variational inference method that uses a Gaussian prior on the latent distribution of cell types. Our work differs from this as we use a Dirichlet prior for the distribution of the topics in the spots, which leads to an interpretable latent space. Notably, unlike in the case of a Gaussian VAE, the Dirichlet distribution enforces that the latent topic weight vector 𝒛 only has positive topic membership components.

Here, we used DeepTopics to study the spatial organization of synovial tissues from seropositive RA patients that contain ELS on histology to identify shared microenvironments across tissues. We defined 22 topics, many of which encompass multiple colocalizing cell types and a range of predicted biological functions, that inform on possible cellular interactions and pathologic pathways. We then trained DeepTopics on ELS microenvironments and showed that cellular ELS composition and architecture vary significantly between patients, and that these ELS topic enrichments may be reflected in patient serologic status. Thus, our data support complex cellular and molecular heterogeneity within histologically similar tissues.

## Results

### DeepTopics identifies spatially defined topics and demonstrates superior scalability compared to traditional topic models

To investigate the spatial organization of synovial niches in RA, we applied the 10x Visium CytAssist^23^ workflow to eight tissue sections from six seropositive patients undergoing arthroplasty who met 2010 American College of Rheumatology/European League Against Rheumatism (ACR/EULAR)^24^ classification criteria and had moderate or high disease activity (**Supplementary Table 1**). All eight synovial sections displayed lymphocytic inflammation with numerous well-defined lymphoid aggregates and a high degree of plasma cell infiltration, as observed on hematoxylin and eosin (H&E) histological sections.

We aggregated the eight spatial transcriptomics datasets corresponding to the tissue sections and trained a single DeepTopics model on the combined dataset (**Figure 1A**, **Methods**). The model encoder learns to map spot-level transcript count data to the parameters of a Dirichlet distribution of dimension *k*, a hyperparameter of the model. Sampling from the Dirichlet distribution yields a multinomial distribution over *k* topics, where each topic is in turn represented as a multinomial distribution over genes and stored in the decoder. We chose *k* = 25 based on model selection to ensure that each topic had sufficient discriminative marker genes (**Methods**). We then applied this model to each of the eight datasets to infer topic weights (also called topic posterior probabilities) for each spot, which we visualized over the spot array (**Figure 1B**). Three topics were driven by high mitochondrial transcripts or genes highly expressed by all cells and were not analyzed further. We annotated each remaining topic by identifying the most discriminative genes and by inferring topic posteriors across scRNA-seq clusters from the Accelerating Medicines Partnership (AMP) RA synovial single-cell atlas^25^ (**Figure 1C**). To compare our approach to cell type annotation, we also scored each spot using gene signatures of known cell types (**Supplementary Figure 1**, **Methods**).

**Figure 1:**
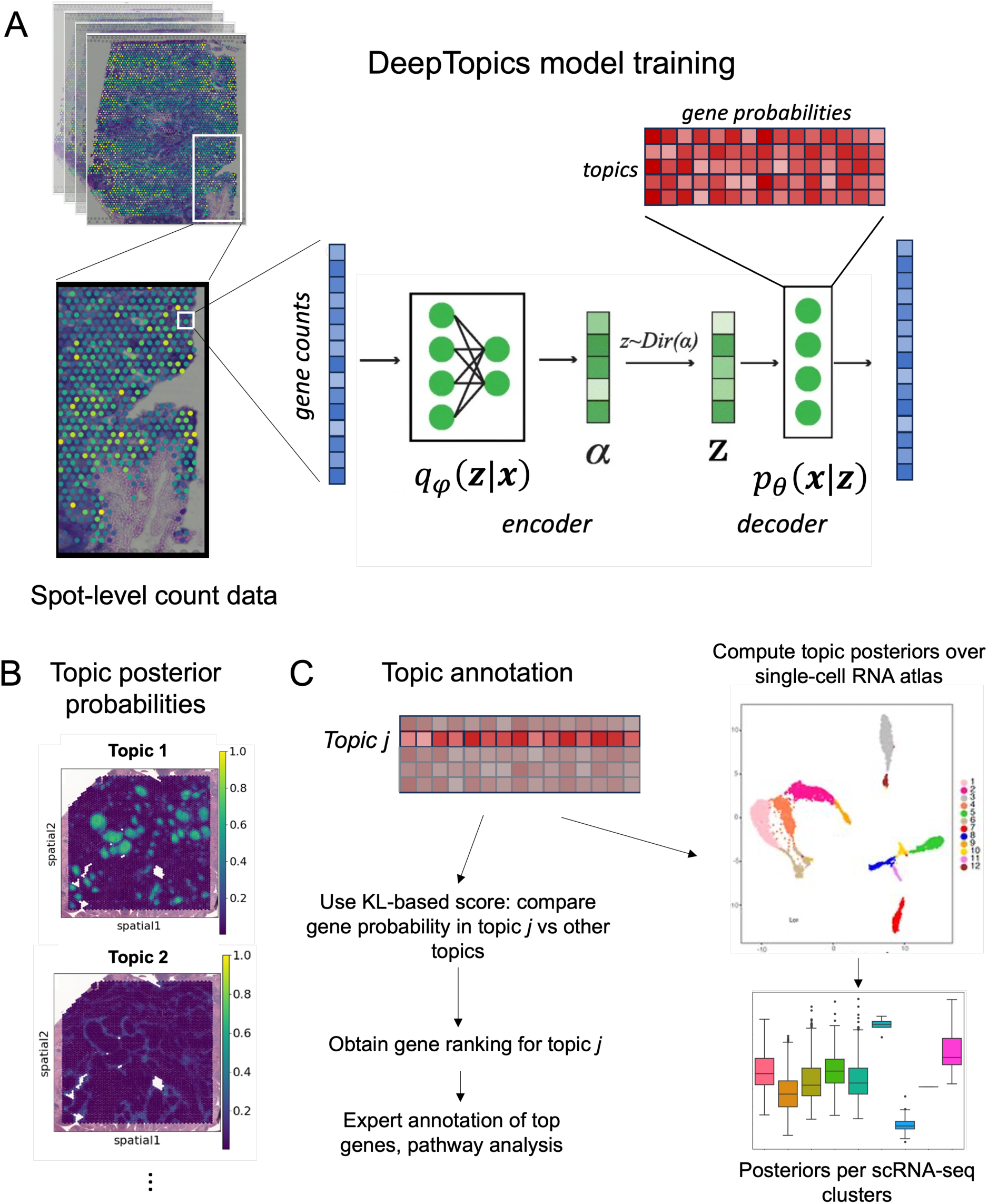
DeepTopics analysis enables unsupervised ST deconvolution and topic annotation for RA synovial tissue. **A.** Model training. ST datasets generated from synovial tissue sections (N = 8) were aggregated into a combined corpus of documents (spots), and one topic model was trained on raw spot-level count of data. The encoder maps the count data for a spot to the parameter vector of a Dirichlet distribution. Sampling from this distribution gives the posterior probability over topics. Each topic is a multinomial probability distribution over all the genes, stored in the decoder, and the model learns the topics as well as the encoder-decoder parameters that accurately reconstruct the count data. **B.** Visualization of the topic posterior probabilities in each spot (shown for one tissue section). **C.** Topic annotation. The top-ranked genes for each topic using a KL-based score were used for expert annotation and pathway analysis, and the trained model was applied to an annotated RA scRNA-seq atlas to compute topic posteriors over all cells and determined topic enrichments over cell types.

To demonstrate the scalability of our method compared to more traditional topic modeling methods for ST, such as LDA or NMF-based models, we ran DeepTopics on one, two, four, six, and all eight of the RA synovium ST datasets (resulting in datasets containing 4722, 9506, 18951, 28039, and 37592 spots, respectively). We then performed the same benchmarking using the R-based LDA topic modeling method STdeconvolve^16^, the R-based Poisson NMF topic modeling method fastTopics^18^, and a Python-based LDA topic model implemented in scikit-learn^26^ (**Supplementary Table 2**). We found that DeepTopics took 5.25 hours to train on all eight samples, compared to 24.51 hours for fastTopics and 72.42 hours for the scikit-learn LDA model (**Supplementary Table 2**). STdeconvolve could not be used to train on all eight datasets due to lack of computing resources. Taken together, these results confirmed the superior scalability of our model.

### DeepTopics identifies cellular gene expression programs and communities

When we trained one DeepTopics model on all eight ST datasets, we defined 22 topics based on distinct patterns of gene expression. For each topic, we applied a Kullback-Leibler (KL)-based^27^ score to identify genes that discriminate the topic from all other topics,(**Methods**, **Supplementary Table 3).** The resulting gene expression signatures enabled annotation of each topic as containing one or more known synovial cell type (**Figure 2A**, **Supplementary Figure 2**). For example, topic 1 had high KL scores for multiple lymphocyte genes (T cell: *CD3D*, *CCR7*^28^, *IL7R*, *FOXP3*^29^, *CXCL13*; B cell: *MS4A1*^30^, *IGHD*) and dendritic cell genes (*LAMP3*). Topics 2 and 3 had high KL scores for genes expressed by IgG (*IGHG1*) and IgM (*IGHM*) plasma cells, respectively. Several topics were enriched for known markers of synovial myeloid cells, including topics 11 (LYVE1^31^) and 14 (MERTK^32^).

**Figure 2:**
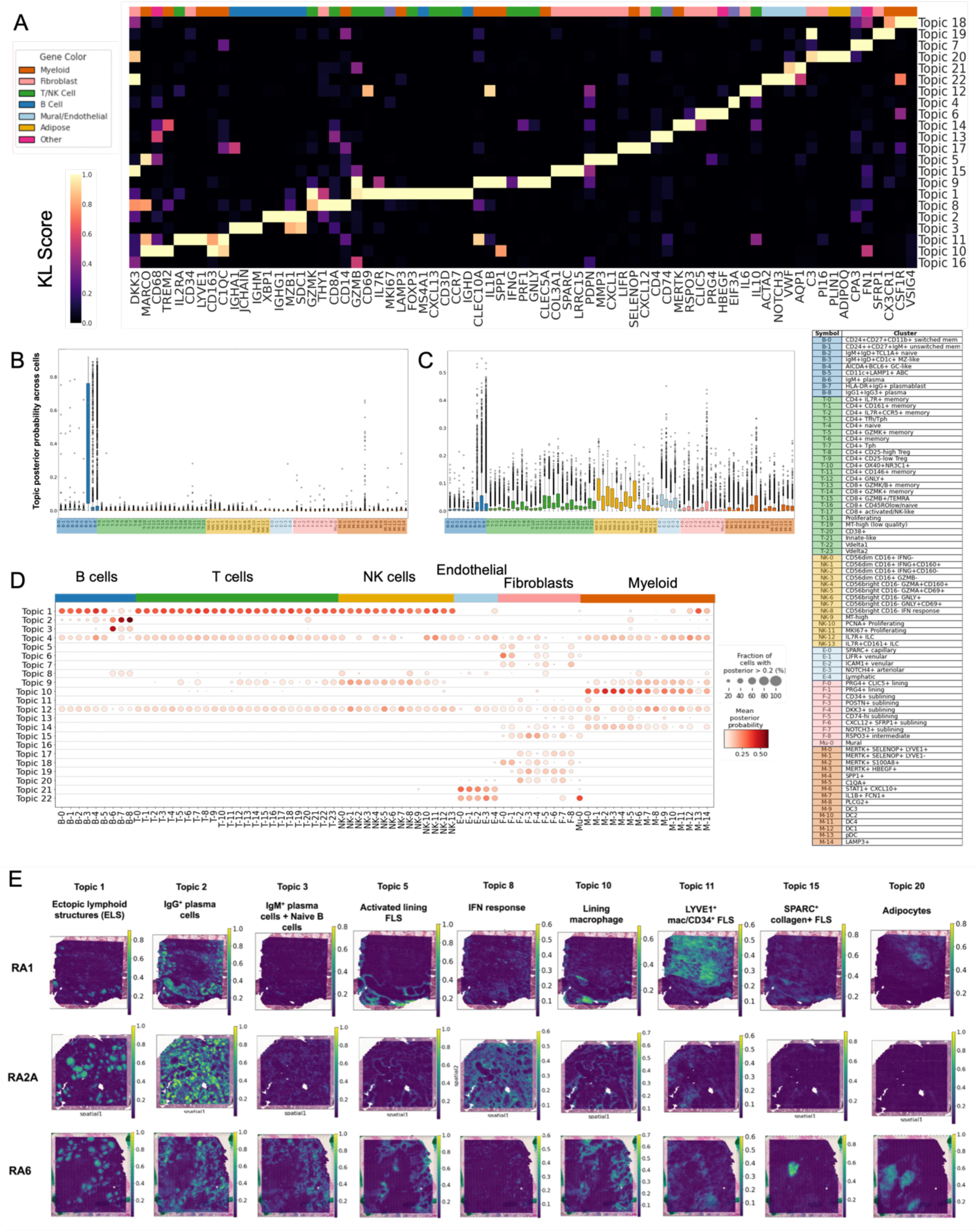
Topic modeling reveals collaborating cell types and spatial organization. **A.** Min-max normalized KL-based scores for known RA synovial cell type marker genes across topics. Column colors indicate the cell type in which the marker gene is predominantly found within the synovium^44^. **B,C.** Posterior probabilities of topic 3 and 8, respectively, across cells in the AMP RA scRNA-seq atlas^44^ grouped by cell type. Each box represents the interquartile range of posterior probabilities for the topic across cells belonging to the cluster. Posterior probabilities of each topic across the cells in the AMP scRNA-seq atlas in **Supplementary Figure 3. D.** Mean posterior probability of each topic in each cell cluster with a mean posterior probability >0.2, normalized using min-max scaling across cells. Labels for each cell type cluster found in the scRNA-seq atlas used for topic annotation. These labels and colors are used to refer to each cell type cluster in panels (**B**), (**C**), and (**D**). **E.** The spatial distribution of selected topics (8 of 22) in two of the eight tissue sections. Posterior probabilities of each topic were obtained for each spot and overlaid onto the corresponding H&E. Full list of topics and their spatial distribution are shown in **Supplementary Figure 4.**

As expected for resident stromal cells that are found throughout the synovium, fibroblast-like synoviocytes (FLS) were found in multiple topics. Lining FLS markers^33^ were found in both topics 5 and 6, with topic 5 displaying the more activated phenotype as evidenced by enriched expression of matrix metalloproteinase *MMP3* and the chemokine *CXCL1*. Topic 6 gene expression patterns were similar to that of the “resting lining” state that we defined previously^6^ and exhibited higher expression of the lining marker gene *CLIC5* as well as genes associated with production of synovial fluid components: *PRG4* (lubricin, a large proteoglycan), *XYLT1* (xylosyltransferase 1, which is required for the synthesis of glycosaminoglycan chains on proteoglycans) and *PCSK6* (subtilisin/kexin-like protease PACE4, a protease implicated in aggrecan degradation and proteoglycan release by human cartilage^34^). Topics 15, 17, and 19 have high KL scores for sublining FLS markers such as SPARC (a fibrotic fibroblast marker^35^), CXCL12, and SFRP1^36^, respectively.

Topic 20 was enriched for adipocyte markers such as ADIPOQ^37^ and PLIN1^38^. Detection of adipocytes is of interest as they have not been included in existing scRNA-seq efforts given the differences in cell isolation protocols needed to isolate adipocytes. Topic 20 additionally includes expression of complement protein *C3*, primarily expressed by FLS, and *PI16*, a marker of both progenitor-like fibroblasts^6, 39^ and adipocyte progenitors^40^. Topics 21 and 22 were enriched for endothelial and mural cell markers (*VWF*^41^, *AQP1*^42^, and *ACTA2*^43^). Of note, across all topics, while some were predicted to be dominated by a single cell type based on gene expression, such as topics 2 and 3 by plasma cells, others were predicted to be a mixture of cell types, such as topic 1, as mentioned above, with multiple lymphocyte subsets including CXCL13^+^ T cells, FOXP3^+^ regulatory T cells, IgD^+^ naïve B cells, and LAMP3^+^ DCs, or topic 11 with a mix of genes expressed by progenitor-like FLS (*CD34*, *MFAP5*, *DPT*) and myeloid cells (*CD163*, *LYVE1*, *C1QC*) (**Figure 2A**). The predicted co-occurrence of multiple cell types in a single topic introduces the prospect of identifying novel cellular interactions and microenvironments.

To further explore the range of cell types in each topic, we applied the trained DeepTopics model to the recently published AMP RA scRNA-seq atlas^25^ that contains 77 RA synovial cell types, including B, T, NK, myeloid, FLS, and endothelial cell types (**Figures 2B-D**, **Supplementary Figure 3**). Comparing the topic posterior probabilities in each scRNA cell cluster vs. all other clusters by a one-sided Mann Whitney U test confirmed that some topics were relatively cell-type-specific, while others represented expression programs of multiple cell types (**Figures 2C-D**, **Supplementary Figure 3)**. Topic 3 is most enriched in naïve B cells: B-2 (IgM+ plasma cells) (U = 3.43 x 10^8^, adjusted p < 10^-16^) and B-6 (IgM+ IgD+ TCLA1+ naive) (U = 1.17 x 10^8^, adjusted p < 10^-16^) (**Figure 2C**), whereas topic 11 is most enriched in specific subtypes of FLS and myeloid cells: F-2 (CD34^+^ sublining FLS) (U = 5.78 x 10^8^, adjusted p < 10^-16^) and M-0 (MERTK^+^SELEOP^+^LYVE1^+^ macrophages) (U = 7.34 x 10^8^, adjusted p < 10^-16^) in agreement with the KL-based gene signature for this topic (described above). Topic 20, which includes adipocytes as per KL scoring above, was also enriched in multiple FLS sublining subtypes, including CD34^+^ (U = 6.94 x 10^8^, adjusted p < 10^-16^), DKK3^+^ (U = 1.99 x 10^8^, adjusted p = 5.90 x 10^-3^), CXCL12^+^ SFRP1^+^ (U = 7.49 x 10^8^, adjusted p < 10^-16^), and NOTCH3^+^ (U = 2.00 x 10^8^, adjusted p = 5.72 x 10^-55^) (**Supplementary Figure 3**), supporting a possible role for FLS-adipocyte interactions in synovial homeostasis or pathology^44^. Topic 8 is notable for an enrichment in specific cell states across multiple cell types: including several NK cell clusters such as NK-2 (CD56^low^CD16^+^IFNG^+^CD160^-^) (U = 3.35 x 10^7^, adjusted p = 4.88 x10^-11^) and NK-8 (CD56^hi^ CD16^-^ IFN response (U = 1.68 x 10^6^, adjusted p = 3.19 x 10^-3^). Further, topic 8 contained cells in B-7 (HLA-DR^+^IgG^+^ plasmablasts), T-12/13/15 (IFNγ producing) CD4^+^ and CD8^+^ T cells, F-5 (CD74^hi^ sublining) FLS and M-6 (STAT1^+^CXCL10^+^) macrophages all have high relative mean posterior probabilities for topic 8 (**Figure 2D**). The presence of distinct cell types that produce and respond to IFNγ in topic 8 suggests that DeepTopics can identify not only the localization of cell types but also shared biological processes and functions involving multiple cell types. Identification of this microniche is particularly relevant to RA given that JAK inhibitors, which inhibit signaling downstream of IFNγ, are an approved treatment option for patients that have failed other therapies^45^.

To further analyze biological processes represented in topics, we performed pathway analysis on top ranked genes for each topic using g:Profiler^46^ and confirmed that certain topics were defined by shared function rather than by enrichment for specific cell types (**Supplementary Table 4**). For example, topic 12 was enriched for pathways such as TNF signaling, AP-1, and interleukins. While topic 12 was enriched in subsets of all major cell types (**Figure 2E, Supplementary Figure 3**), the pathway results suggest that this topic is likely dominated by cytokine signaling pathways shared across cell types. Pathway analysis also allowed annotation of topics that did not have dominant cell type enrichments, such as topic 16, which is enriched for genes associated with all sublining FLS as well as some myeloid subsets. Pathway analysis of topic 16 identified pathways of “developmental processes”, “anatomical structure development”, “regulation of cell migration”, and “extracellular matrix (ECM) organization” leading us to a topic annotation of “stromal homeostasis”. Topic 18 was also associated with genes expressed by both lining and sublining FLS and was enriched for “ECM organization” as well as “cell-substrate adhesion” pathways. However, it did not exhibit enrichment of developmental processes as topic 16 did. Thus, we termed topic 18 “ECM organization”.

Interestingly, every topic containing FLS except topic 15 (SPARC^+^ FLS) was enriched for pathways associated with the complement system, suggesting a widespread role for complement in FLS function in RA^47, 48^. As expected for topic 20, which is predicted to contain adipocytes, there was an enrichment for lipid metabolism and storage pathways.

We next visualized the posterior probabilities of each topic superimposed on the H&E of each tissue slice (**Figure 2F**, **Supplementary Figure 4**) to identify tissue niches. The location of several topics, such as lining FLS topics 5 and 6 and ELS topic 1, were as expected based on the gene signature lists and predicted cellular makeup. Plasma cell topics 2 and 3 localized in areas surrounding ELS as previously published^49^, but topic 2 (IgG^+^ plasma cells) also covered large swaths of a subset of the tissues (RA2A and RA4). We further observed topic localizations that offered new insights into cellular function, such as in the two myeloid-rich topics that can localize to the lining, topic 10 and topic 14. Topic 10 is enriched for multiple macrophage subsets (**Figure 2D**) as well as the gene *MMP3*, which is primarily expressed by the activated lining FLS. Topic 14 exhibits expression of macrophage marker *MERTK*, which is expressed in the lining in RA in remission but not active RA^50^ as well as genes primarily expressed by resting lining FLS (e.g., *CLIC5, XYLT1)*. Interestingly, Topic 14 also exhibits *RSPO3* expression, possibly implicating Wnt signaling in the maintenance of this topic’s expression program^51^. Our observation of macrophage populations associated with more (topic 10) versus less (topic 14) FLS activation suggests that these two states may be stable microenvironments, and additional work is needed to understand their relative functions.

### RA synovial tissues exhibit marked variation in topic abundance and distribution

While DeepTopics was trained on all 8 tissue sections and learned a unified set of topics applicable to all sections, we observed large variations in topic abundance between tissue sections and especially between tissues derived from different patients. To quantify this variation, we calculated the cumulative distribution functions (CDFs) of the posterior probabilities for each topic across the spots in each of the eight samples (selected topics shown in **Figure 3A**, full set of topics in **Supplementary Figure 5**). We assessed the significance of these differences by performing one-sided Mann Whitney U tests to identify the samples that were significantly enriched for each topic (**Methods**). The effect sizes associated with these comparisons were quantified using rank-biserial correlation^52^ (**Figure 3B**, **Methods**). Notably, RA4 is heavily enriched for interleukin signaling topic 12 (U = 9.02 x 10^7^, adjusted p < 10^-16^, effect size = 0.87), while both sections from RA2 (RA2A and RA2B) are enriched for interferon response topic 8 (RA2A: U = 7.03 x 10^7^, adjusted p < 10^-16^, effect size = 0.73; RA2B: U = 7.03 x 10^7^, adjusted p < 10^-16^, effect size = 0.43), compared to all other samples. We also saw significant differences in IgG and IgM plasma cell topic abundance (topics 2 and 3, respectively). Tissue sections from patients RA5 and RA6 were significantly enriched for the IgM plasma cell/naive B cell topic (RA5A: U = 1.27 x 10^8^, adjusted p < 10^-16^, effect size = 0.40; RA5B: U = 1.27 x 10^8^, adjusted p < 10^-16^, effect size = 0.44; and RA6: U = 1.27 x 10^8^, adjusted p < 10^-16^, effect size = 0.58), but not the IgG^+^ plasma cell topic (topic 2). Interestingly, these two patients also had high titer serum rheumatoid factor (**Supplementary Table 1**). Given that rheumatoid factor is most commonly an IgM isotype and that synovial plasma cells are known to locally produce immunoglobulin, including rheumatoid factor^53^, it is possible that this abundance of tissue IgM^+^ plasma cells (topic 3) is related to the high titer serum rheumatoid factor. Finally, RA3, one of the samples with less pronounced ectopic lymphoid structures on H&E (along with RA1), was significantly enriched for ECM organization topic 18 (U = 9.23 x 10^7^, adjusted p < 10^-16^, effect size = 0.78), SPARC^+^ FLS topic 15 (U = 7.59 x 10^7^, adjusted p < 10^-16^, effect size = 0.40), stromal homeostasis (topic 16) (U = 1.31 x 10^8^, adjusted p < 10^-16^, effect size = 0.43), and SFRP1^+^ FLS topic 19 (U = 7.38 x 10^7^, adjusted p < 10^-16^, effect size = 0.41), suggesting that this tissue, even with the presence of ELS, could be more similar to the previously defined fibroblast-rich, pauci-immune endotype^25, 5^.

**Figure 3:**
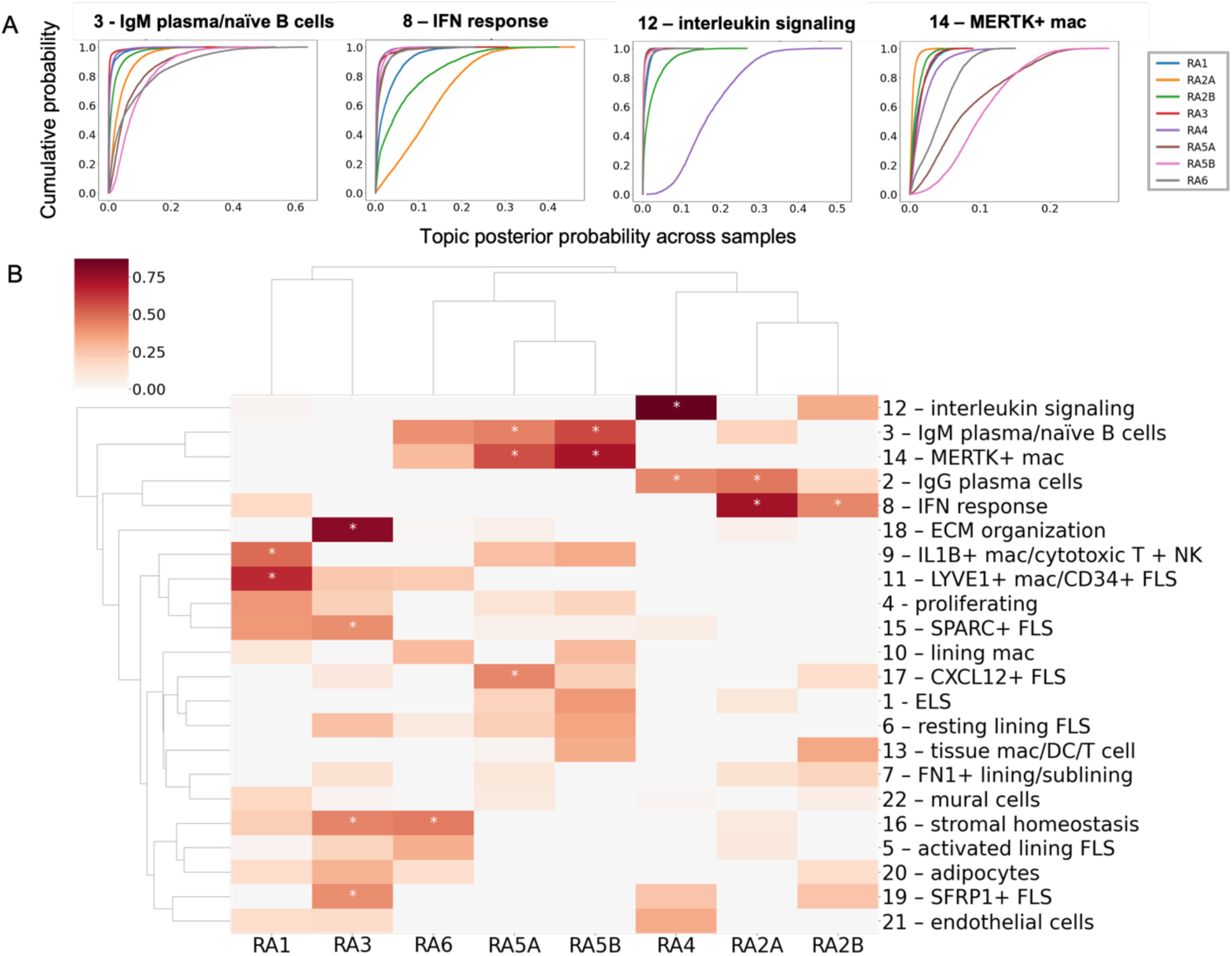
Topics generated by DeepTopics display differential enrichment across synovial samples of the same RA pathotype. **A.** Empirical cumulative distribution functions (CDFs) of the posterior probabilities of selected individual topics across all spots in each sample. **B.** Enrichment of each topic in every sample. Red shading indicates statistically significant topic enrichment in the given sample compared to the full set of samples (Mann Whitney U test, adjusted p < 0.05 considered significant). Shading color scale indicates the effect size (rank-biserial correlation) of the given topic in the given sample. Asterisks (*) indicates enrichment > 0.4.

To quantify topic co-localization across all spots in individual samples, we used the global Lee’s L spatial statistic^54^ which measures co-occurrence in spots and their neighbors (**Methods**, **Figure 4**, and **Supplementary Figure 6**). As expected, topics are generally segregated into sublining and lining localizations. For example, in samples RA2A (**Figure 4A**) and RA5A (**Figure 4B**), topics corresponding to resting lining FLS (topic 5), activated lining FLS (topic 6), and lining macrophages (topic 10) all significantly co-localize (adjusted p = 0.002 in RA2A and 0.0021 in RA5A). Similarly, in these two samples, several sublining topics (e.g., CXCL12^+^ FLS (topic 17) and SFRP1^+^ FLS (topic 19)) significantly colocalize (adjusted p = 0.002 and 0.0021, respectively). Notably, there were also differences in topic colocalizations between samples and patients. Topic 4 (proliferating cells) co-localizes with different topics, likely indicating variation in the types of dividing cells between tissue sections. While this topic significantly co-localizes with ELS (topic 1) in RA2A (adjusted p = 0.002), this colocalization is unique to this tissue section. In other sections, topic 4 co-localizes with LYVE1^+^ macrophage/CD34^+^ FLS (topic 11) (RA1, adjusted p = 0.0023; RA5A, adjusted p = 0.0021; RA5B, adjusted p = 0.0022), adipocytes (topic 20) (RA2B, adjusted p = 0.0022; RA3, adjusted p = 0.0021), SPARC^+^ FLS (RA6, adjusted p = 0.0023), and CXCL12^+^ FLS (RA3, adjusted p = 0.0021; RA4, adjusted p = 0.0024; RA5B, adjusted p = 0.0022). Some samples, such as RA2A, had less distinct groupings of sublining and lining topics, as observed by both lining and sublining colocalizations of the MERTK^+^ macrophage topic (topic 14) (**Figure 4A**). In addition, SPARC^+^ FLS (topic 15) and the FN1^+^ lining/sublining (topic 7) colocalize with sublining topics in some tissues but with lining topics in others (**Figure 4A**). These differences may indicate a fluidity between lining and sublining environments in the inflamed synovium, which is supported by a previous mouse model study in which the induction of arthritis resulted in an expansion of FLS with an intermediate lining/sublining phenotype^55^. It is also possible that due to the spatial resolution of the 10x Visium technology, the RNA capture spots associated with the lining also capture sublining cells directly adjacent to the lining.

**Figure 4:**
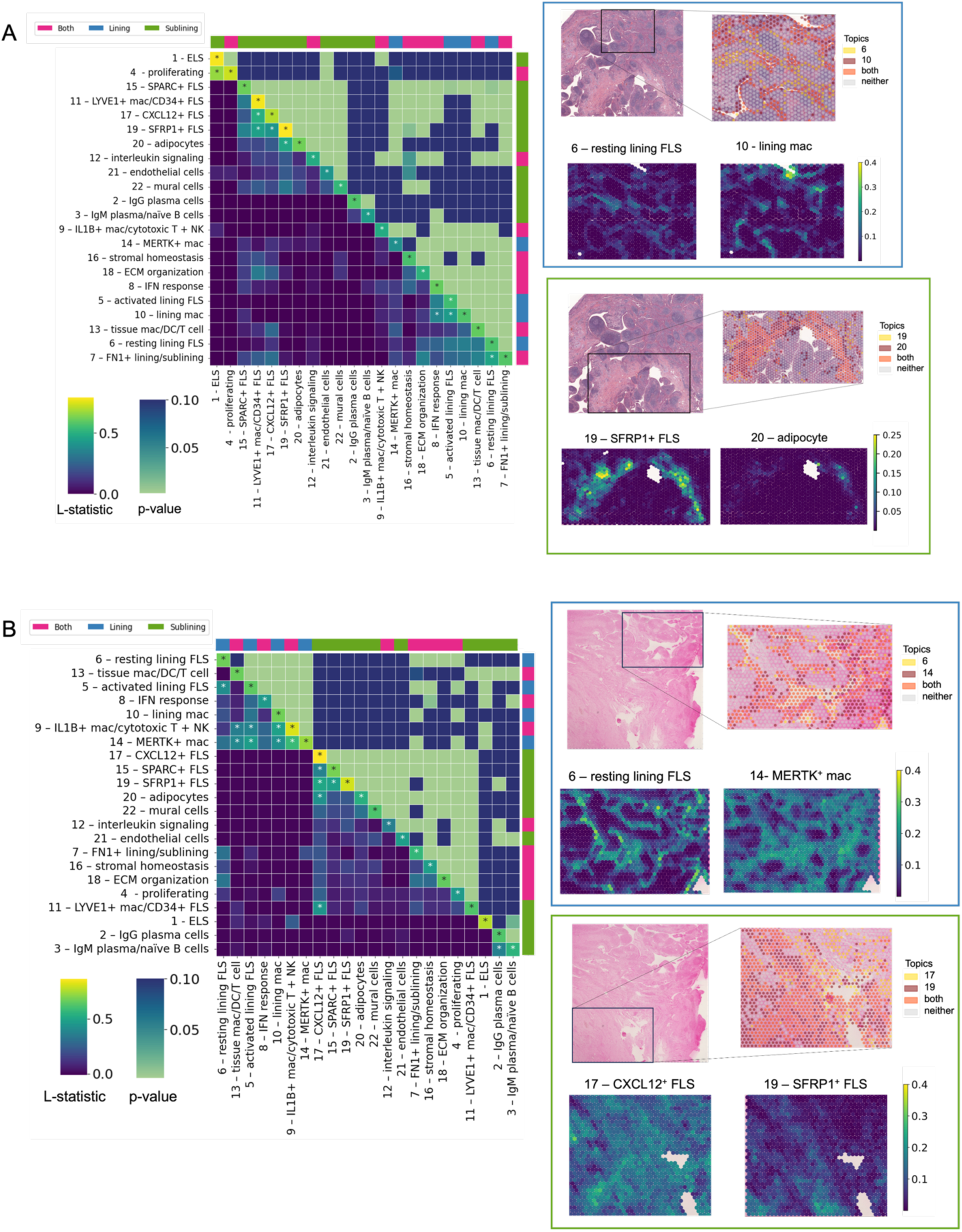
Spatial statistics reveal plasticity in synovial lining and sublining cell states. A, **B.** Statistically significant (Lee’s L statistic, adjusted p < 0.05) spatial correlations of topics in two samples: (**A**) RA2A and (**B**) RA5A. L statistic for each topic pair shown on the lower half of the heatmap and the diagonal. Lee’s L p-value shown on the upper half of the heatmap. Larger values of Lee’s L are represented by brighter colors on the lower half of the heatmap and the diagonal, indicating a larger correlation, and asterisks (*) indicate strong correlations (Lee’s L > 0.35). Brighter colors on the upper half of the heatmap indicate lower adjusted p-values (higher significance). Outer bar colors indicate synovial localization of the given topic (sublining, lining, both). Spatial distributions of selected topics from each category are also shown for both samples (right). Blue boxes represent lining FLS topics and green boxes represent sublining FLS topics. Region of interest from H&E shown with RNA capture spots colored based on topic posterior probabilities above the 75^th^ percentile for indicated topics.

There were also differences in tissue localization of topics implicated in cytokine signaling pathways that are targeted by approved RA therapies. Topic 8, which is enriched for IFNγ producing and responding cells, can strongly localize to either the synovial lining (RA2) or the sublining (RA3), as has been described previously^6^. This differential localization appears to be dependent on the patient from which the tissue section is derived, as sections from the same patient exhibit similar topic 8 localization (**Figure 2F** and **Supplementary Figure 4**). Both IFN response (topic 8) and IL1B^+^ macrophages/cytotoxic T/NK cells (topic 9) colocalized with lining topics in all samples except RA3, indicating a possible different tissue architecture in this sample as highlighted above by its abundance of FLS-associated topics. Interleukin signaling (topic 12) colocalizes with lining topics in RA1, RA4, RA5B, and RA6 (adjusted p < 0.05 in all cases) but with sublining topics in RA2A, RA2B, RA3 and RA5A (adjusted p < 0.05 in all cases). While the association of cytokine signaling with the synovial lining may be counterintuitive given the sublining localization of most lymphocytes, these results are in agreement with previous reports of an exaggerated cytokine signaling response observed in lining FLS^6^. The tissue localization of cytokine signaling pathways to the lining versus sublining in different tissues may have implications for targeting these cytokines in the treatment of RA. One hypothesis is that the synovial microenvironment in which cytokine production/response occurs may play a role in determining the efficacy, either positively or negatively, of cytokine inhibitors such as TNF or IL-6 receptor inhibitors.

This spatial analysis highlighted the discrete tissue niches formed by ELS and plasma cells. ELS (topic 1) did not significantly colocalize with any other topic except proliferating cells (topic 4) in RA2A (**Figure 4A**). Additionally, the IgG and IgM plasma cell topics do not significantly colocalize with any topics except one another (see RA5A in **Figure 4B**). These results suggest that ELS and plasma cells form distinct microenvironments, which may stem from the fact that these cell types and structures are not found in the healthy synovium. Given the discrete tissue microenvironment of ELS suggested by the spatial colocalization analysis, we next performed a focused analysis on these structures.

### Ectopic lymphoid structures (ELS) exhibit both cell type and architectural heterogeneity

In early RA, 40-50% of patient synovia have a lympho-myeloid pathotype associated with the presence of ELS and high plasma cell accumulation^4, 5^. ELS may also have implications for predicting treatment response as patients with synovial ELS are more likely to respond favorably to treatment with tumor necrosis factor inhibitors (TNFi)^56, 57, 58^. A possible pathogenic role for synovial ELS is suggested by their ability to support local B cell maturation and activation. Synovial ELS exhibit evidence of B cell somatic hypermutation^59, 2^, isotype class switching^59, 2^, clonal expansion^2, 3^, and *in situ* production of autoantibodies^59, 60^. In fact, a subset of ELS, possibly up to 20%^61^, are able to support the formation of structures that have features of a germinal center. It remains unclear how the synovial stromal architecture supports ELS formation and what drives the formation of functional germinal centers in some, but not all, ELS.

To address these questions, we used performed hotspot analysis using the Getis-ord GI* statistic^62^ to identify capture spots with highest abundance of topic 1 (ELS) whose neighbors also highly express this topic (**Figure 5A**, **Methods**). We then aggregated the set of all identified spots across every sample and trained a second DeepTopics model on these spots alone (**Figure 5B**, **Methods**). Performing model selection as before, we trained an eight-topic model and carried out topic annotation as described above (**Figure 1C**). KL scores for selected genes of interest are shown in **Figure 5C**, with the top 100 ranked genes in each topic provided in **Supplementary Table 5**, and the pathway analysis results in **Supplementary Table 6**. Finally, we applied the trained model to the raw RNA counts of the same ELS capture spots in each sample to determine topic posteriors across cells to visualize the spatial distribution of topics within each ELS (**Figure 5D, Supplementary Figure 7)**. We also applied the trained model to the AMP RA scRNA-seq atlas^25^ (**Supplementary Figure 8**) to aid in topic annotation.

**Figure 5:**
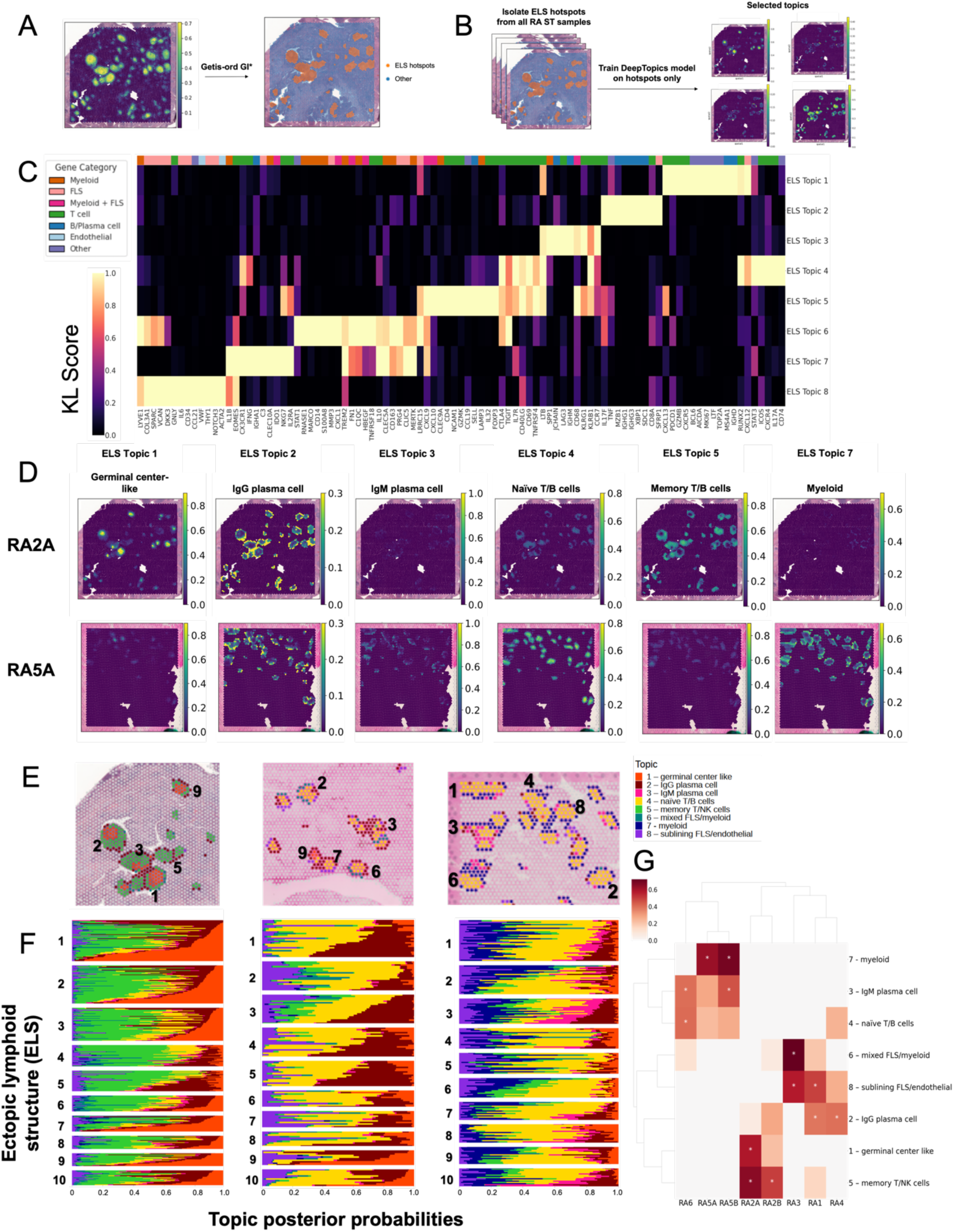
ELS deconvolution reveals patient-driven variation in cell composition. **A.** Identification of the ELS topic hotspots using the Getis-ord GI* statistic. **B.** Diagram of the training process on the ELS spots. The hotspots across each sample were aggregated into one combined set, and one model was trained on count data for these spots. **C.** KL scores of marker genes of known RA synovial cell types. **D.** Spatial distribution of the posterior probabilities of selected topics in selected samples. Results for all tissue sections in **Supplementary Figure 7**. **E.** Topic compositions of individual ELS plotted on the spatial H&E for selected samples: RA4 (left), RA5A (middle), and RA5B (right). Results for additional tissues shown in **Supplementary Figure 9**. Each spot in each ELS is colored according to the topic with the maximum posterior probability in that spot. Spots numbered according to size (1 = largest based on the number of RNA capture spots encompassed by the ELS). **F.** Stacked bar plot of topic posterior probability in each spot within each ELS; ELS numbers and topic colors as in panel (**E**). **G.** Enrichment of each ELS topic in every sample. Red shading indicates statistically significant topic enrichment in the given sample compared to the full set of samples (Mann Whitney U test, adjusted p < 0.05 considered significant). Shading color scale indicates effect size (rank-biserial correlation) of the given topic in the given sample. Asterisks (*) indicates enrichment > 0.4.

Within the capture spots identified as ELS, we identified multiple immune-cell rich topics as well as those with dominant stromal cell gene expression signatures (e.g., FLS, macrophage, and endothelial cells). We identified a germinal center-like topic (ELS topic 1) with expression of germinal center markers *AICDA* and *BCL6* as well as proliferation markers *MKI67* and *TOP2A*. ELS topic 1 also exhibits expression of chemokines implicated in ectopic germinal center formation, *CXCR5* and *CXCL13*^63^. Using the AMP RA scRNA-seq atlas^25^, we found enrichment of all non-plasma cell B cell subsets (B0-5 in **Figure 2B-D**). Of note, the areas with the highest abundance of ELS topic 1 expression overlay with areas on the H&E where there are central clearings within an ELS, which are characteristic of germinal centers. In line with the expected occurrence of germinal centers in the synovium, not all ELS exhibit expression of germinal center-like topic 1. Circumferentially around areas of ELS topic 1 expression, we observed ELS topic 5, which had predicted enrichment for memory T cell subtypes including CD8^+^ *GZMK/GZMB^+^*memory, CD4^+^ TPH and CD8^+^ *GZMB*^+^ TEMRA. Some ELS, especially in tissue sections from RA5 and RA6, are instead dominated by central expression abundance of ELS topic 4, which has predicted enrichment of naïve B and T cell subsets (IgM^+^ *IGHD^+^ TCL1A^+^* naïve B cells, CD4^+^ naïve T cells and CD56^hi^ CD16^-^ *GNLY^+^* CD69^+^ NK cells). Around these more central regions of the ELS are IgG plasma cells (ELS topic 2), IgM plasma cells (ELS topic 3), and three non-lymphocyte topics: mixed fibroblast/myeloid (ELS topic 6), myeloid (ELS topic 7) and sublining FLS/endothelial cells (ELS topic 8). Even within an ELS, there is evidence of multiple stromal and myeloid cell topics that likely modulate lymphocyte activation and function.

From the spatial distribution of ELS topic posterior probabilities on the tissue sections, we observed that topic abundance varies both by patient and by individual ELS. For example, both sections from RA2 were enriched for the germinal center-like topic (ELS topic 1), but individual ELS exhibited variations in the amount of topic 1 expression (**Figure 5D, Supplementary Figure 7**). To further investigate these differences, we isolated all ELS covering at least ten RNA capture spots and visualized the dominant topic (defined as the topic with the maximum posterior probability) in each spot on the tissue section (**Figure 5E**) as well as the posterior probabilities of each ELS topic in each spot in individual ELS (**Figure 5E-F**, **Supplementary Figure 9**). These results highlight different topic abundances between ELS in the same tissue as well as ELS cellular compositions across patients. While ELS in RA2A and RA2B had strong representation of either the germinal center-like topic (ELS topic 1) or the memory T and B cell topic (ELS topic 5), the ELS in RA4 and RA5 were dominated by the naive T and B cell topic (ELS topic 3). Interestingly, these naive T and B cells are surrounded by IgG plasma cells in RA4, while in RA5A, they are surrounded by the myeloid topic, as supported by global Lee’s L statistics (**Supplementary Figure 10**).

When we compared the overall topic abundance across all ELS from each sample (**Figure 5G, Supplementary Figure 11**), we observed clear differences in topic co-occurrence between samples. First, there appeared to be two overall patterns of ELS topic expression: (i) dominance of the naïve T and B cell topic along with IgM plasma cells (ELS topic 3), and (ii) abundance of the germinal center and/or memory T and B cell topics along with IgG plasma cells (ELS topic 2). Comparing these results with the visualization of ELS topic expression on the tissue sections, it appears that ELS centers comprise either germinal center-like cells or naïve T and B cells, depending on the patient. It is important to consider that the tissue sections analyzed are one snapshot in time and restricted to a single section, which could be missing the true “center” of an ELS. Thus, some of our observations could be due to differences in ELS maturation/age or due to spatial sampling effects. However, by including many ELS (140 FLS annotated as in **Figure 5E**) in our analysis and restricting to those ELS that encompassed at least 10 RNA capture spots, we believe that we have identified real variations in established ELS. Overall, we have found that even between samples with well-defined ELS on H&E, there is heterogeneity in cell composition in ELS pointing towards differences in factors driving their formation and maintenance.

## Discussion

In disease states, alterations in tissue function are likely not only due to the cell types and states present but also to their spatial organization within the tissue. This spatial information has multiple possible implications for treatment: (1) the formation of defined disease-associated microenvironments may propagate pathogenic signaling pathways, making them resistant to inhibition through existing therapies, and (2) pathways relevant to microenvironments may not be targeted by any existing therapies and instead may represent an opportunity for the development of novel therapies. These issues are of interest in RA, where up to 17% of patients have “difficult-to-treat” RA, defined as persistent disease activity after trials of multiple medications^64, 65^. Here, we apply ST to RA synovial tissue to understand microenvironments associated with tissue inflammation in patients with established RA. These patients had moderate disease activity (CDAI > 10) at the time of tissue excision, as well as a robust lymphocyte infiltration on synovial histology, both indicating ongoing active disease with a wide range of disease durations (0.6 – 28 years) (**Supplementary Figure 1**). By applying a novel deep topic modeling method that is not dependent on an existing scRNA-seq atlas and exhibits superior scalability to existing methods, we identified shared topics across patient tissues each containing one or more cell types with common cellular programs. These topics suggest both novel cellular interactions and disease-relevant microenvironments. Notably, in these patients with shared histologic features, such as lymphocyte infiltration and multiple ELS, we uncovered marked differences in topic abundance across patients as well as two divergent ELS structures with implications for disease pathogenesis.

While some topics were dominated by single cell types (e.g., topics 2 and 3 by plasma cells), some topics included multiple cell types and states suggesting novel interactions. In topic 11, LYVE1^+^ FOLR2^+^ macrophages and CD34^+^ FLS co-occur with *IL10* expression (**Figure 2A, D**), which may represent an anti-inflammatory niche. LYVE1^+^ macrophages are enriched in RA patients in sustained disease remission but can also be found in the sublining of patients with active disease^50^. These macrophages express more resolvin, an inflammation resolving lipid mediator, and IL-10, an anti-inflammatory cytokine, as compared to pro-inflammatory cytokines^50^. CD34^+^ FLS demonstrate a progenitor-like gene expression program and exhibit dampened responses to exogenous cytokine stimulation^39, 6^. These findings suggest that LYVE1^+^ macrophages and CD34^+^ FLS may form an anti-inflammatory tissue niche that contributes to restoring synovial tissue homeostasis. Work in limb ischemia suggests that this interaction could be mediated by oncostatin M (OSM) signaling^66^. Additional work is needed to understand this interaction in the synovium.

Our analysis of relative topic enrichment across tissue sections (**Figure 3**) suggested three patterns: (1) RA1 and RA3 were enriched for multiple fibroblast topics (SPARC^+^ FLS, SFRP1^+^ FLS) as well as stromal homeostasis, ECM organization, and LYVE1^+^ macrophage/CD34^+^ FLS topics; (2) RA2 and RA4 were enriched in IgG plasma cells as well as interleukin signaling (RA4) or IFN response (RA2); and (3) RA5 and RA6 were enriched in IgM plasma cells/naïve B cells. Although RA5 and RA6 expressed the IgG plasma cell topic (topic 2), it was the relative excess of IgM plasma cells that was notable. This dichotomization of IgM plasma cell topic abundance was even clearer in our ELS analysis (**Figure 5**) in which ELS in some patients were encircled by plasma cells of only IgG isotypes (RA1, RA2, RA3 and RA4) and in others by a relative abundance of IgM isotypes (RA5 and RA6). Notably, RA5 and RA6 had high titer serum rheumatoid factors (RF) (>20 times the upper limit of normal), which is an IgM isotype. Thus, it is possible that the excess RF detected in the serum is being locally produced by IgM^+^ synovial plasma cells and reflected in the relative enrichment of IgM plasma cell topics in RA5 and RA6. The presence of IgM plasma cells indicates that although some synovial B cells have undergone class switching from naïve IgM/IgD subsets to memory IgG isotypes, class switching is not universally observed even in patients with long-standing disease (23 and 28 years in RA5 and RA6, respectively). Further work will be needed to determine if high titer serum RF, which is associated with higher disease activity^67^ and more severe systemic manifestations such as rheumatoid vasculitis or the formation of rheumatoid nodules^68^, is associated with synovial IgM enrichment.

Recent studies using scRNA-seq have shown the heterogeneity of cell states present in RA synovial tissue during active disease, and the co-occurrence of specific cell subtypes within individual synovial samples suggests that there are multiple tissue endotypes observed in disease^6, 69^. In this work, we observed stable topic abundance between sections from a given tissue, supporting the existence of tissue endotypes such as the recently proposed Cell Type Abundance Phenotypes (CTAPs)^25^ and adding a layer of complexity to previously defined synovial pathotypes^5^. Importantly, our data also suggest that multiple microenvironments, as represented by the topics, exist in each tissue, possibly complicating this picture. What drives the formation of tissue endotypes and what role tissue microenvironments may play in this process has yet to be determined. The stark spatial differences defined in this work across histologically similar synovia highlight the need to further investigate how specific tissue microenvironments may relate to clinical features and treatment response. ST will undoubtedly play an important role in our understanding of tissue level pathophysiology for multiple diseases, and DeepTopics provides a novel, highly scalable, reference-free method for ST analysis to support future studies.

## Methods

### Human synovial tissue

RA synovial tissue was obtained from patients enrolled in the Hospital for Special Surgery FLARE study of RA patients undergoing arthroplasty or synovectomy (approved by HSS IRB #2014-233). Enrollment complied with all relevant ethical regulations and informed consent was obtained from all participants. At the time of tissue collection, all patients were being treated with methotrexate. One patient was also on low dose prednisone and other oral disease modifying antirheumatic drugs (DMARDs) (triple therapy). Another patient was on targeted biologic therapy (Janus kinase (JAK) inhibitor) that was held prior to surgery per established clinical recommendations^70^. Synovial tissue was formalin fixed and paraffin embedded (FFPE). RA synovial tissue was scored for lymphocytic inflammation, plasma cell infiltrate, and degree of lining hyperplasia through histologic analysis (H&E) by a musculoskeletal pathologist.

### Spatial transcriptomics by 10X Genomics Visium with CytAssist

We selected 8 tissue sections from 6 patients (**Supplementary Table 1**) for analysis using Visium CytAssist Spatial Gene Expression platform (10x Genomics) in conjunction with the Integrated Genomics Operation and Molecular Cytology core facilities at the Memorial Sloan-Kettering Cancer Center (MSK). Each tissue section was placed on a CytAssist slide (6.5 mm x 6.5 mm area) with 4993 capture spots per area. Slides were prepared, including deparaffinization and staining with hematoxylin & eosin (H&E), by the Molecular Cytology Core at MSK. After imaging, slides were de-stained by incubation 15 minutes at 42°C with 0.1N HCl and de-crosslinked by incubation at 95°C for 1 hour with Decrosslinking Buffer (10X Genomics)^23^. Probe pairs from the Human Transcriptome Probe Kit v2 (10X Genomics PN 1000466)^23^ targeting 18,536 genes (54,018 probes) covering the whole transcriptome were added to the slides and allowed to hybridize overnight at 50°C. Bound pairs were ligated to one another, and slides were washed according to instructions provided by the manufacturer. Slides were loaded on the Visium CytAssist Instrument for probe release from the tissue onto the Visium CytAssist Spatial Gene Expression Slide. Probe extension, elution, and library preparation proceeded with the Visium FFPE Reagent Kit v2 (10X Genomics PN 1000436)^23^ according to the manufacturer’s protocol with 8-12 cycles of PCR as determined by real-time PCR. Indexed libraries were pooled and sequenced on a NovaSeq X in a PE28/88 run using the NovaSeq X 10B or 25B Reagent Kit (Illumina)^71^. An average of 163 million paired reads was generated per sample, corresponding to ∼41K reads per spot. An average of 4699 (SD = 191) spots were analyzed per tissue section.

For each tissue section, all spots present were analyzed. DeepTopics finds topics with genes that correspond to low quality spots, such as mitochondrial genes that are excessively expressed in some spots; we exclude these topics from downstream analyses. Also, uninformative or background genes that are too rare (i.e., that are in fewer than 10 spots) or too ubiquitous (i.e., that are in more than 95% of the spots) are also excluded from further analysis.

### Gene signature scoring

Prior to deconvolution, we curated gene signatures for the following cell types: endothelial, tissue macrophage, CD14+ monocyte, CD16+ monocyte, dendritic cells, pDCs, lining FLS, sublining FLS, progenitor FLS, naive CD20+ B cells, B cells, plasma cells and plasmablasts, CD4 Tph/Tfh cells, CD4 T cells, CD8 T cells, naive T cells, NK cells, neutrophils, and mast cells, and the spots in each ST dataset were scored for these gene signatures using the score_genes function in Scanpy^72^.

### Dirichlet VAE model design and implementation

Similar to Latent Dirichlet Allocation (LDA) topic model-based ST deconvolution, the Dirichlet VAE architecture we use in DeepTopics was originally developed in natural language processing (NLP), where the goal is to learn multiple topics present in a corpus of documents from word count (bag-of-words) data. In the NLP setting, every LDA topic is a multinomial probability distribution over words, and the word count data in every document is generated via a multinomial distribution over topics, also called posterior topic probabilities. In applying LDA to ST data, each spot (capture region) in the input ST data is treated as a document and the genes as words; the learned topics can be viewed as cellular expression programs^15^. Specifically, input spot 𝑖 is a represented as a vector of transcript counts indexed by genes, 𝒙_𝑖_ = ’𝑔_𝑖,1_, 𝑔_𝑖,2_, … , 𝑔_𝑖,𝑇_*, where 𝑇 is the total number of genes in the ‘vocabulary’ of the model. In either LDA or a DirVAE topic model, 𝒙_𝑖_ is viewed as a ‘bag of genes’ generated by multinomial 𝒛_𝑖_ over topics, where each topic 𝑗 is defined by a multinomial 𝜽_j_ over genes, and the number of topics, 𝑘, is a hyperparameter. In LDA, parameters of the multinomials 𝒛_𝑖_ and 𝜽_j_are governed by a pair of Dirichlet distributions governing the ‘peakiness’ of topic posteriors for each document and of genes in each topic, respectively.

The Dirichlet VAE model implemented as DeepTopics uses analogous concepts but encapsulates the topic decomposition of spot-level count data in a probabilistic encoder-decoder architecture (**Figure 1A**). The encoder uses a neural network to transform the bag of genes 𝒙_𝑖_ into 𝜶_𝑖_, a *k*-dimensional vector representing the parameters of a Dirichlet distribution. The model then samples the topic weight vector 𝒛_𝑖_ (of size 1 x 𝑘), from this distribution Dir(𝜶_𝑖_), specifying the parameters of a multinomial distribution over topics. Next, the decoder uses the topic parameters 𝜽, a matrix of size 𝑘 x 𝑇 that specifies the multinomial distribution of gene probabilities for each topic (in row 𝜽_j_), together with the topic proportions 𝒛_𝑖_ to reconstruct the input count data as 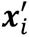. The reconstruction error between 𝒙_𝑖_ and 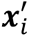 contributes to the overall loss of the model, as described below.

### Encoder-decoder network architecture

The encoder takes a *T*-dimensional gene count vector as input and passes it through 3 hidden layers with 300 units in each layer, which are fully connected, followed by the so-called "bottleneck" layer with *k* units. The encoder thereby reduces the T-dimensional input spot to a *k-* dimensional Dirichlet parameter vector 𝜶_𝑖_. The topic vector 𝒛_𝑖_ that is sampled from Dir(𝜶_𝑖_) is passed to the decoder which maps it via the output reconstruction layer into a *T*-dimensional output vector. The total number of parameters in the model is (T * 300) + (300 * 300) + (300 * 300) + (300 * k) + (k * T).

### Model training process

DeepTopics optimizes the following objective function, called the evidence lower bound (ELBO):

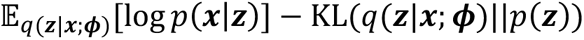

The first term, 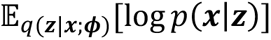, can be interpreted as the reconstruction error of the decoder (parameterized by 𝜽) in reproducing 𝒙 given the latent topic vector 𝒛, where 𝒛 is sampled from the Dirichlet prior produced by the encoder network, which approximates the posterior distribution 𝑞(𝒛|𝒙; 𝝓). The second term measures the Kullback-Leibler (KL) divergence between the approximate posterior 𝑞(𝒛|𝒙; 𝝓) and the prior 𝑝(𝒛).

Akin to the reparameterization trick^73^ used in Gaussian VAEs that allows the propagation of gradients in a neural network through a probabilistic layer, DeepTopics uses the implicit reparameterization gradients-based approach from Figurnov et al.^74^, which offers unbiased estimators for continuous distributions such as a Dirichlet distribution. These gradients are used in a stochastic gradient descent (SGD) optimization that trains the model in two steps: (1) a burn-in stage where the encoder-generated Dirichlet parameters 𝜶 are fixed to the input hyperparameters and only the decoder parameters 𝜽 are updated; and (2) second stage where the posterior distribution parameters 𝝓 are also optimized along with 𝜽.

For this study, the genes sequenced by 10x Visium CytAssist^23^ were used as the vocabulary for the DirVAE topic model. In each of the eight ST datasets, genes that were detected in fewer than ten or more than 95% of the spots were removed. The raw count matrices of all eight datasets were combined into one combined corpus of 37,592 spots, and the DirVAE model was trained on these raw transcript counts. Several combinations of hyperparameters were tried (see **Supplementary Table 5** for the values tested for each hyperparameter), and models were assessed based on the requirement to produce at least 30 discriminative marker genes for each topic (see below). Based on this criterion, the following combination yielded optimal results: 𝛼_3_ (the Dirichlet hyperparameter) = 10, *k* (the number of latent features, or topics) = 25, batch size = 64, prior burn-in steps (i.e., the number of steps run by the model before updating the Dirichlet parameter) = 80000, and maximum steps = 200000.

### Model inference and visualization

The trained DeepTopics model was applied to each of the eight ST datasets separately, and each row of the resulting spot-topic matrices contained the “membership” (posterior probability) of each topic in that spot. For the inference results for each sample, the matrix containing the topic assignments for each spot (i.e., the spot-topic matrix) was row-normalized, such that each spot was represented as a multinomial distribution over topics. The topic posterior probabilities across all spots in each ST sample were visualized using the sc.pl.spatial function in the Scanpy analysis suite^72^.

### Topic annotation

To annotate each topic, we used the approach from Dey et al.^27^, which ranks genes using a KL-based score that measures the similarity between the probability of the gene in a given topic versus all other topics. The equation for this KL-based score 𝐾𝐿^4^[𝑘] is as follows:

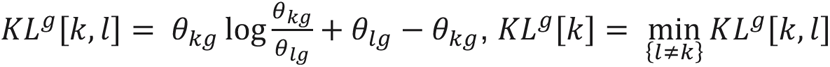

For each topic, genes in the vocabulary were ranked in descending order by their KL scores. The top ranked genes were further analyzed by domain experts to provide initial labels for each topic as a particular cell type or cell type mixture. Pathway analysis was conducted by applying the R package g:Profiler v0.2.113^46^ to the top 100 ranked genes in each topic. FDR correction was performed using the Benjamini-Hochberg method, and adjusted p-values below 0.05 were considered significant.

To further validate the topic labels, we utilized the AMP RA scRNA-seq atlas from Zhang et al.^25^ to detect enrichment of topics in cell types. After removing genes that were not present in the vocabulary used for model training, we applied the trained DirVAE topic model to the raw cell-level counts from the atlas, and the distribution of posterior probabilities of each topic across the cell clusters were plotted using Scanpy^72^. The clusters that were most highly enriched for the expression of each topic, comparing the posterior probability distributions in each cluster versus all other clusters, were determined using a one-sided Mann Whitney U test. Final topic annotations were determined using a combination of the results of the model application on the scRNA-seq atlas, the pathway analysis results, and manual analysis of the top ranked genes in each topic.

### Analysis of spatial variation of topics across samples

After inference, we found that certain topics were enriched only in a subset of the eight samples on which the model was trained. To compare the posterior distributions of each topic across all eight samples, the cumulative distribution functions of the posterior probabilities of each topic were computed for each sample. Topic enrichment was quantified using a one-sided Mann Whitney U test, and the effect size was calculated using rank biserial correlation^52^. False discovery rate (FDR) correction was applied using the Benjamini-Hochberg method. We filtered for adjusted p-values below 0.05 and effect sizes greater than 0.4.

### Spatial colocalization of topics

To measure topic colocalization across spots, a global Lee’s L statistic^54^ was calculated for every pair of topics in each sample. The diagonal of the weights matrix for Lee’s L was set to 1, imposing a high spatial weight for a spot with itself, while immediately adjacent spots were given a weight of 0.5. FDR correction was applied using the Benjamini-Hochberg method, and we filtered for adjusted p-values below 0.05 and values of the Lee’s L statistic above 0.35.

### Further deconvolution of ELS

Topic 1 was identified as the ELS topic based on its ranked gene list, the application of the model to the AMP scRNA-seq atlas, and comparison of the topic’s spatial distribution to the corresponding H&E section of each sample. Thus, for each of the eight ST samples, spots that were significantly enriched for topic 1, known as “hotspots”, were identified using the Getis-ord GI* statistic^62^ with a threshold of 1.96 standard deviations above the mean (97.5th percentile).

The hotspots for topic 1 were aggregated across all eight samples, and a second DeepTopics model was trained on the counts data for these spots. Several hyperparameter combinations were attempted, and the following combination yielded optimal results based on annotation by domain experts: alpha = 3, K = 8, batch size = 16, prior burn-in steps = 120000, and maximum steps = 200000. Each of the eight topics were annotated manually using the top 100 ranked genes, and the annotations were validated using inference on the AMP RA scRNA-seq atlas^25^ and pathway analysis with g:Profiler^46^ as before. Finally, the trained model was applied on the ELS hotspots from each individual sample, and the topic posteriors were visualized using Scanpy^72^.

For each sample, to further examine the topic distribution in each ELS, we used Loupe Browser v8.0.0 to identify the spatial barcodes associated with each individual ELS^75^. Aggregates that contained fewer than ten spots in each sample were removed, and the DeepTopics and the weight of each topic in every spot in the top ten largest aggregates was visualized using the plotting functions in the R package CountClust^27^. To visualize the cell composition in each ELS, we colored each spot in the ELS hotspots according to the topic with the highest posterior probability in that spot. Finally, to examine the relative enrichment of each ELS topic across an entire sample, we computed the cumulative distribution functions of the posterior probabilities of each topic for the ELS spots in each sample. Enrichment was quantified using a one-sided Mann Whitney U test, and the effect size was calculated with rank biserial correlation^52^ (with false discovery rate (FDR) correction applied using the Benjamini-Hochberg method). We filtered for adjusted p-values below 0.05 and effect sizes greater than 0.4.

### Comparison to existing topic modeling methods for ST deconvolution

To evaluate the scalability of our method compared to LDA and NMF-based methods for ST deconvolution, we benchmarked a DeepTopics model across varying dataset sizes (one, two, four, six, and all eight RA synovium ST datasets). The DeepTopics model was trained using 2 GPUs, each with 7 cores and 20 GB of memory per core. For comparison, we also tested the R-based LDA method STdeconvolve^16^, the R-based Poisson NMF method fastTopics^18^, and a Python-based LDA implementation in scikit-learn^26^ (**Supplementary Table 2**), using a single CPU with 24 cores and 8 GB of memory per core (as these models do not support GPU usage).

We carefully adjusted the hyperparameters of each model to match them as closely as possible, within the constraints of each model’s architecture and implementation. Despite inherent differences between the models, key parameters such as the number of topics (k), learning rates, and iterations were aligned to ensure a fair comparison. The training times for each model were recorded and compared across all datasets.

## Acknowledgements

We thank HSS orthopedic surgeons, clinical research coordinators (particularly Solana Cushing), and the HSS patients who contributed to this study.

We acknowledge the Accelerating Medicines Partnership® (AMP®) in Rheumatoid Arthritis and Lupus Network for the stimulating discussions and the large-scale sequencing of arthritis patient synovial tissues that formed the basis for this study. AMP is a public-private partnership (AbbVie Inc., Arthritis Foundation, Bristol-Myers Squibb Company, Foundation for the National Institutes of Health, GlaxoSmithKline, Janssen Research and Development, LLC, Lupus Foundation of America, Lupus Research Alliance, Merck Sharp & Dohme Corp., National Institute of Allergy and Infectious Diseases, National Institute of Arthritis and Musculoskeletal and Skin Diseases, Pfizer Inc., Rheumatology Research Foundation, Sanofi and Takeda Pharmaceuticals International, Inc.) created to develop new ways of identifying and validating promising biological targets for diagnostics and drug development.

We thank Dr. Frank Curriero from the Department of Epidemiology and the Department of Biostatistics at Johns Hopkins Bloomberg School of Public Health, as well as Dr. Doron Betel from the Weill Cornell Graduate School of Medical Sciences, for their advice on spatial statistics.

We acknowledge the use of the Integrated Genomics Operation Core at the Sloan Kettering Institute, funded by the NCI Cancer Center Support Grant (CCSG, P30 CA08748), Cycle for Survival, and the Marie-Josée and Henry R. Kravis Center for Molecular Oncology.

