## Supplementary figures for "Deep topic modeling of spatial transcriptomics in the rheumatoid arthritis synovium identifies distinct classes of ectopic lymphoid structures"

**Supplementary Table 1:** Patient clinical characteristics tissue histology findings.

| | RA1 | RA2 | RA3 | RA4 | RA5 | RA6 |
|---|---|---|---|---|---|---|
| **Age range (yrs)/ sex** | 50-60/M | 60-70/F | 60-70/F | 40-50/F | 40-50/F | 70-80/F |
| **RA criteria met** | 1987 + 2010 | 2010 | 1987 + 2010 | 2010 | 1987 + 2010 | 2010 |
| **Disease duration (yrs)** | 11 | 7.3 | 10.3 | 0.6 | 23 | 28 |
| **CDAI (on day of tissue collection)** | 20 | 11 | 43 | 22 | 27 | MD |
| **CCP (units)** | >250 | 21 | 26 | 180 | >250 | >250 |
| **RF (IU/mL)** | 37 | 33 | Neg | 34 | 522 | 1000 |
| **Active treatment** | MTX 20mg | Pred 5mg, MTX 17.5mg, SSZ, HCQ | MTX 25mg | MTX 15mg | MTX 7.5mg, upadacitinib | MTX 15mg |
| **Prior treatments** | TNFi | none | TNFi | none | TNFi, abatacept | HCQ |
| **Joint** | Knee | Knee | Knee | Hip | Knee | Knee |
| **Lymphocytic infiltrate** | Dense band-like or numerous large follicle-like aggregates | Dense band-like or numerous large follicle-like aggregates | Numerous lymphocytes or plasma cells, sometimes forming follicle-like aggregates | Numerous lymphocytes or plasma cells, sometimes forming follicle-like aggregates | Dense band-like or numerous large follicle-like aggregates | Dense band-like or numerous large follicle-like aggregates |
| **Lining hyperplasia (cells thick)** | 3-4 | 3-4 | >4 | 2-3 | 2-3 | 3-4 |
| **Plasma cell infiltrate** | >50% | >50% | <50% | >50% | >50% | <50% |

CDAI: clinical disease activity index (≤ 2.8 remission, >2.8-10 low, >10-22 moderate, >22 high), MD: missing data, RF: rheumatoid factor, CCP: anti-cyclic citrullinated peptide antibody, MTX: methotrexate, Pred: prednisone, TNFi: tumor necrosis factor inhibitor, SSZ: sulfasalazine, HCQ: hydroxychloroquine.

Note: Upadacitinib held 4 dosing cycles (4 days) prior to arthroplasty for RA5 in accordance with established recommendations from the American College of Rheumatology.

**Supplementary Table 2:** Scalability of DeepTopics compared to that of other traditional topic modeling ST deconvolution methods. Comparison of the time needed to train datasets of various sizes. Asterisks indicate the use of default parameters where model did not allow for customization of the indicated parameter.

| Method | Programming language | Number of spots | Total memory requested (GB) | Alpha | K | Max steps | Minimum change in loss | Processing time, training (hours) |
|---|---|---|---|---|---|---|---|---|
| **STdeconvolve LDA** | R | 4722 | 192 | 0.1 | 25 | 1000* | -* | 22.642 |
| **STdeconvolve LDA** | R | 9506 | 192 | 0.1 | 25 | 1000* | -* | 40.654 |
| **STdeconvolve LDA** | R | 18951 | 192 | 0.1 | 25 | 1000* | -* | 77.262 |
| **STdeconvolve LDA** | R | 28039 | 192 | 0.1 | 25 | 1000* | -* | 138.558 |
| **STdeconvolve LDA** | R | 37592 | 192 | 0.1 | 25 | 1000* | -* | failed due to lack of computing resources |
| **sklearn LDA** | Python | 4722 | 192 | 0.1 | 25 | 200000 | 0.0005 | 10.016 |
| **sklearn LDA** | Python | 9506 | 192 | 0.1 | 25 | 200000 | 0.0005 | 17.735 |
| **sklearn LDA** | Python | 18951 | 192 | 0.1 | 25 | 200000 | 0.0005 | 44.36 |
| **sklearn LDA** | Python | 28039 | 192 | 0.1 | 25 | 200000 | 0.0005 | 47.692 |
| **sklearn LDA** | Python | 37592 | 192 | 0.1 | 25 | 200000 | 0.0005 | 72.424 |
| **fastTopics NMF** | R | 4722 | 192 | - | 25 | 100* | 0.0005 | 2.939 |
| **fastTopics NMF** | R | 9506 | 192 | - | 25 | 100* | 0.0005 | 4.596 |
| **fastTopics NMF** | R | 18951 | 192 | - | 25 | 100* | 0.0005 | 7.908 |
| **fastTopics NMF** | R | 28039 | 192 | - | 25 | 100* | 0.0005 | 13.339 |
| **fastTopics NMF** | R | 37592 | 192 | - | 25 | 100* | 0.0005 | 24.514 |
| **DeepTopics** | Python | 4722 | 140 | 10 | 25 | 200000 | 0.0005 | 1.043 |
| **DeepTopics** | Python | 9506 | 140 | 10 | 25 | 200000 | 0.0005 | 1.271 |
| **DeepTopics** | Python | 18951 | 140 | 10 | 25 | 200000 | 0.0005 | 2.621 |
| **DeepTopics** | Python | 28039 | 140 | 10 | 25 | 200000 | 0.0005 | 3.167 |
| **DeepTopics** | Python | 37592 | 140 | 10 | 25 | 200000 | 0.0005 | 5.253 |

A

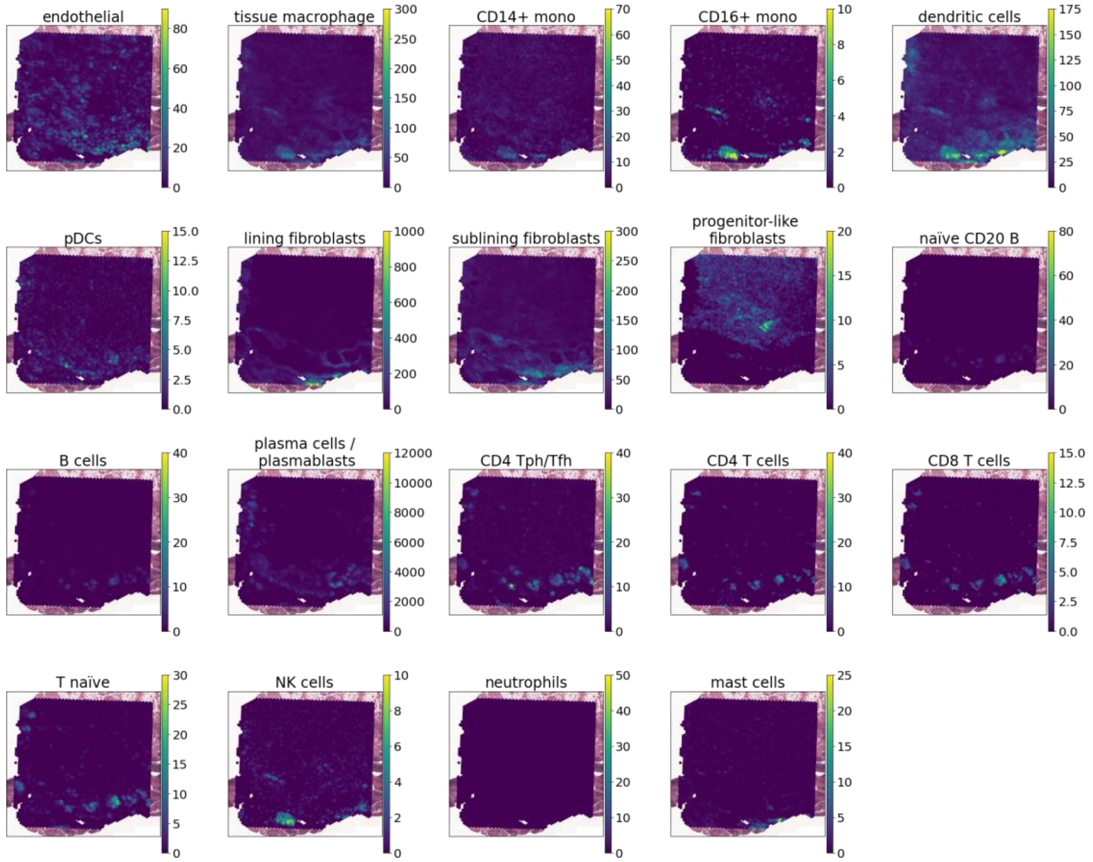
endothelial
tissue macrophage
CD14+ mono
CD16+ mono
dendritic cells
pDCs
lining fibroblasts
sublining fibroblasts
progenitor-like fibroblasts
naïve CD20 B
B cells
plasma cells / plasmablasts
CD4 Tph/Tfh
CD4 T cells
CD8 T cells
T naïve
NK cells
neutrophils
mast cells

B

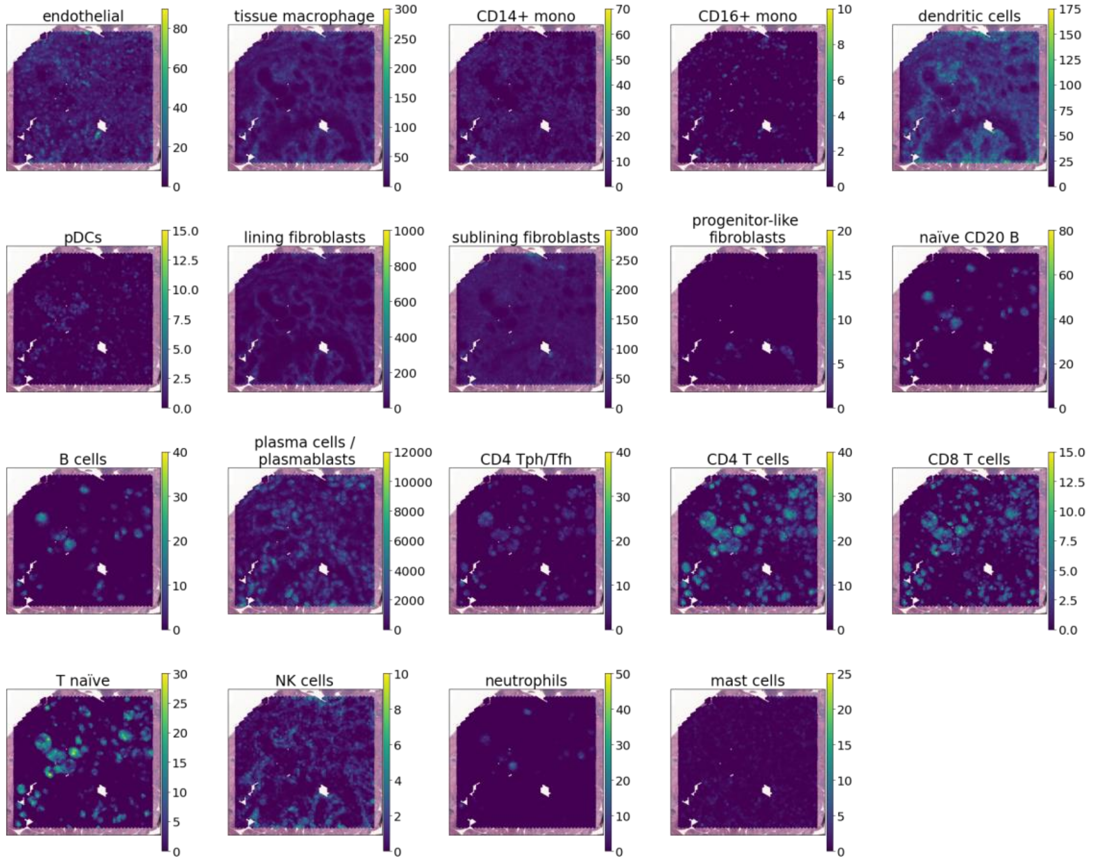
endothelial
tissue macrophage
CD14+ mono
CD16+ mono
dendritic cells
pDCs
lining fibroblasts
sublining fibroblasts
progenitor-like fibroblasts
naïve CD20 B
B cells
plasma cells / plasmablasts
CD4 Tph/Tfh
CD4 T cells
CD8 T cells
T naïve
NK cells
neutrophils
mast cells

C

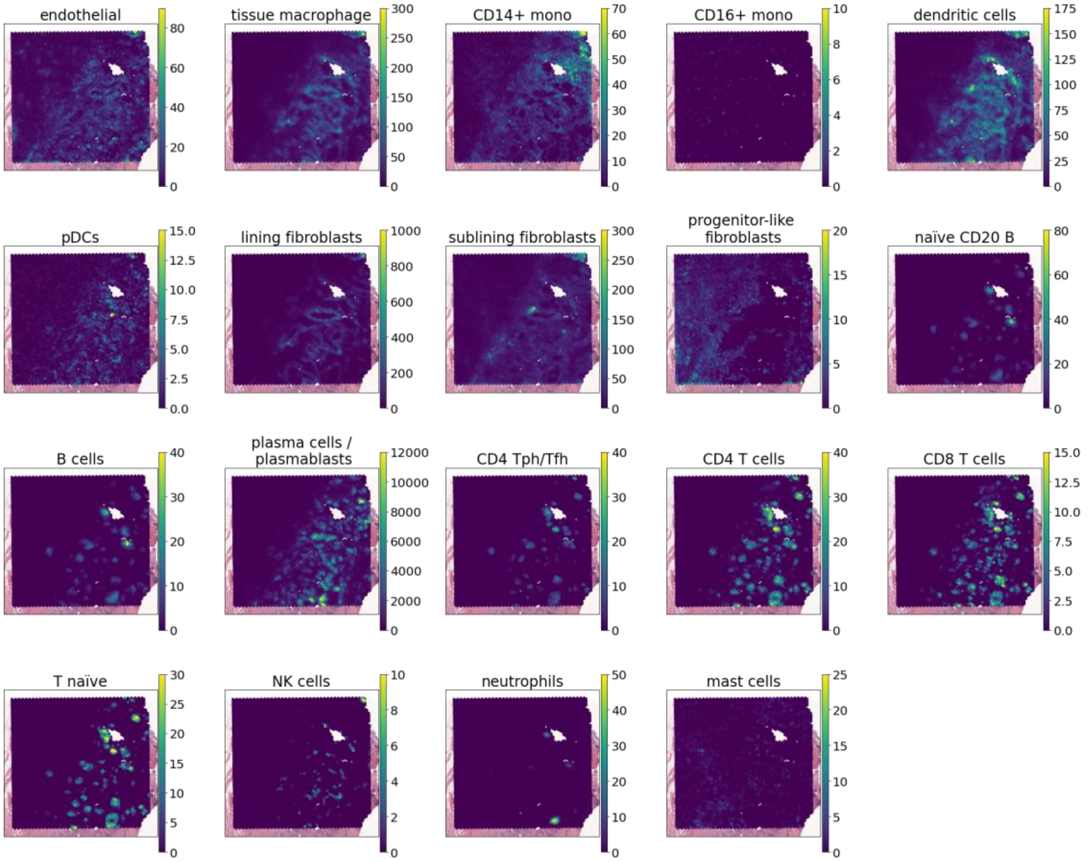
endothelial
tissue macrophage
CD14+ mono
CD16+ mono
dendritic cells
pDCs
lining fibroblasts
sublining fibroblasts
progenitor-like fibroblasts
naïve CD20 B
B cells
plasma cells / plasmablasts
CD4 Tph/Tfh
CD4 T cells
CD8 T cells
T naïve
NK cells
neutrophils
mast cells

D

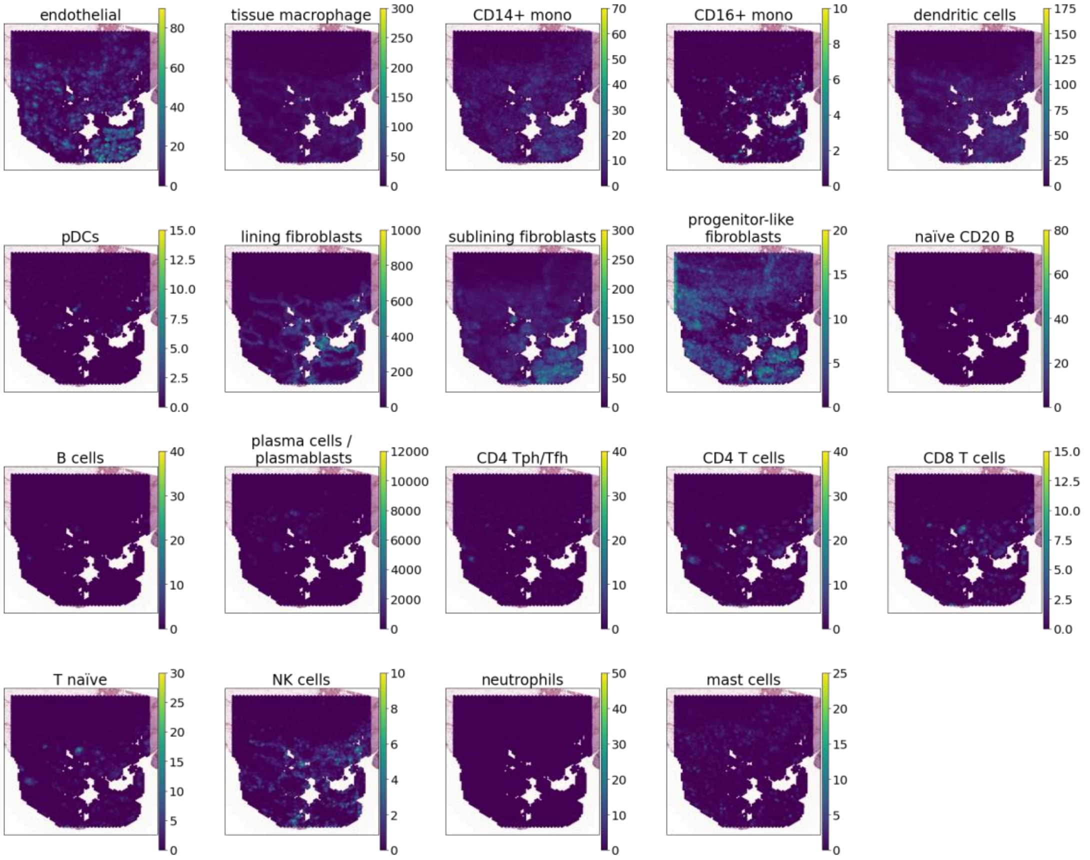
endothelial
tissue macrophage
CD14+ mono
CD16+ mono
dendritic cells
pDCs
lining fibroblasts
sublining fibroblasts
progenitor-like fibroblasts
naïve CD20 B
B cells
plasma cells / plasmablasts
CD4 Tph/Tfh
CD4 T cells
CD8 T cells
T naïve
NK cells
neutrophils
mast cells

E

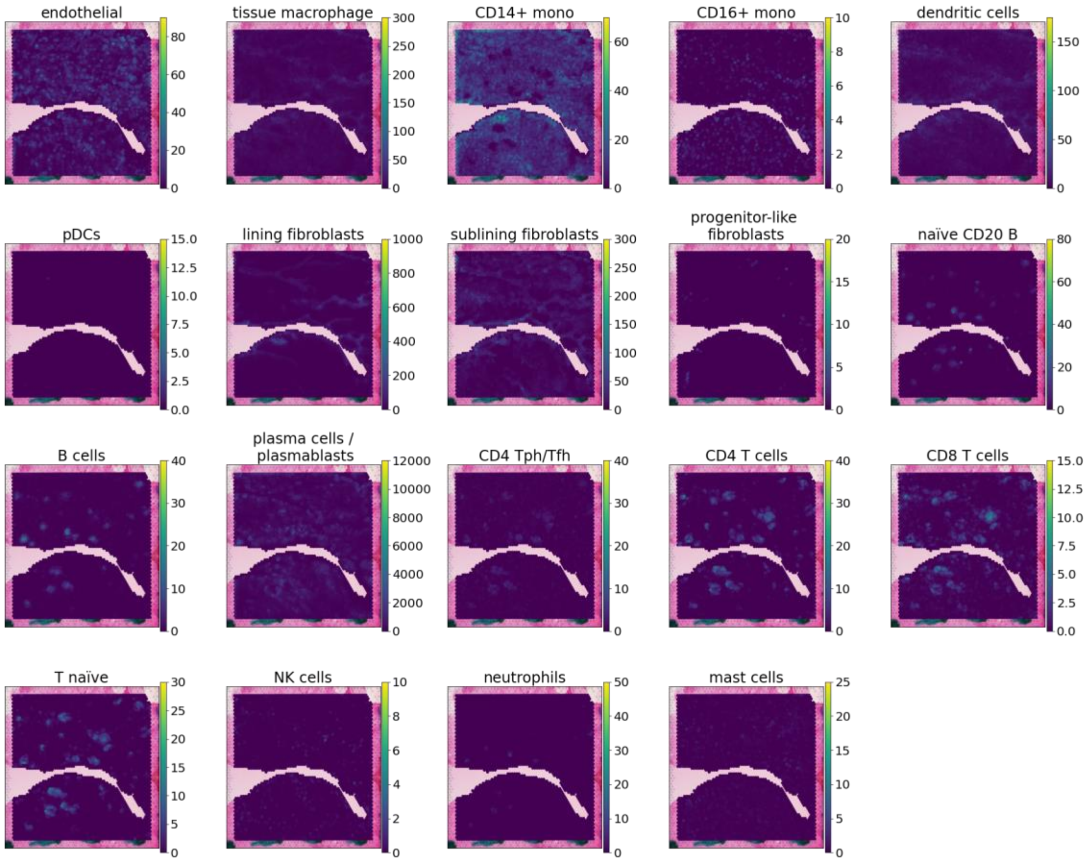

endothelial
0 20 40 60 80
tissue macrophage
0 50 100 150 200 250 300
CD14+ mono
0 20 40 60
CD16+ mono
0 2 4 6 8 10
dendritic cells
0 50 100 150
pDCs
0.0 2.5 5.0 7.5 10.0 12.5 15.0
lining fibroblasts
0 200 400 600 800 1000
sublining fibroblasts
0 50 100 150 200 250 300
progenitor-like fibroblasts
0 5 10 15 20
naïve CD20 B
0 20 40 60 80
B cells
0 10 20 30 40
plasma cells / plasmablasts
0 2000 4000 6000 8000 10000 12000
CD4 Tph/Tfh
0 10 20 30 40
CD4 T cells
0 10 20 30 40
CD8 T cells
0.0 2.5 5.0 7.5 10.0 12.5 15.0
T naïve
0 5 10 15 20 25 30
NK cells
0 2 4 6 8 10
neutrophils
0 10 20 30 40 50
mast cells
0 5 10 15 20 25


F

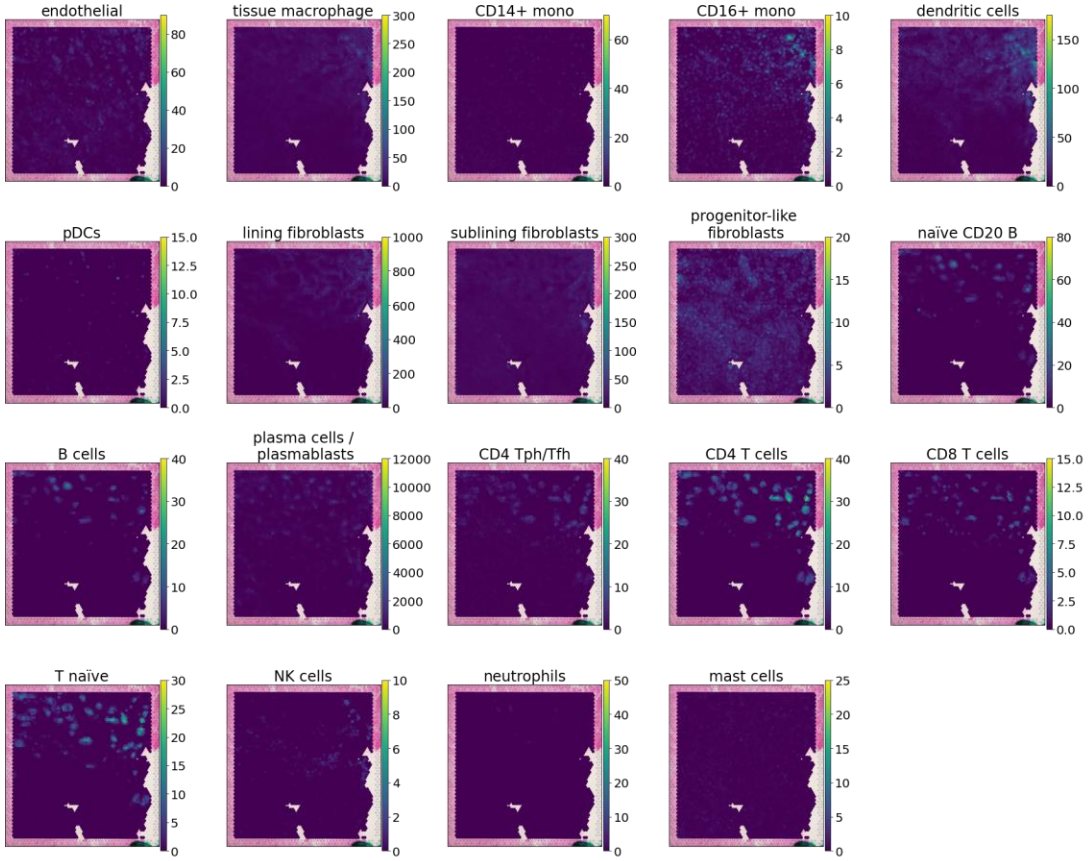

endothelial
0 20 40 60 80
tissue macrophage
0 50 100 150 200 250 300
CD14+ mono
0 20 40 60
CD16+ mono
0 2 4 6 8 10
dendritic cells
0 50 100 150
pDCs
0.0 2.5 5.0 7.5 10.0 12.5 15.0
lining fibroblasts
0 200 400 600 800 1000
sublining fibroblasts
0 50 100 150 200 250 300
progenitor-like fibroblasts
0 5 10 15 20
naïve CD20 B
0 20 40 60 80
B cells
0 10 20 30 40
plasma cells / plasmablasts
0 2000 4000 6000 8000 10000 12000
CD4 Tph/Tfh
0 10 20 30 40
CD4 T cells
0 10 20 30 40
CD8 T cells
0.0 2.5 5.0 7.5 10.0 12.5 15.0
T naïve
0 5 10 15 20 25 30
NK cells
0 2 4 6 8 10
neutrophils
0 10 20 30 40 50
mast cells
0 5 10 15 20 25

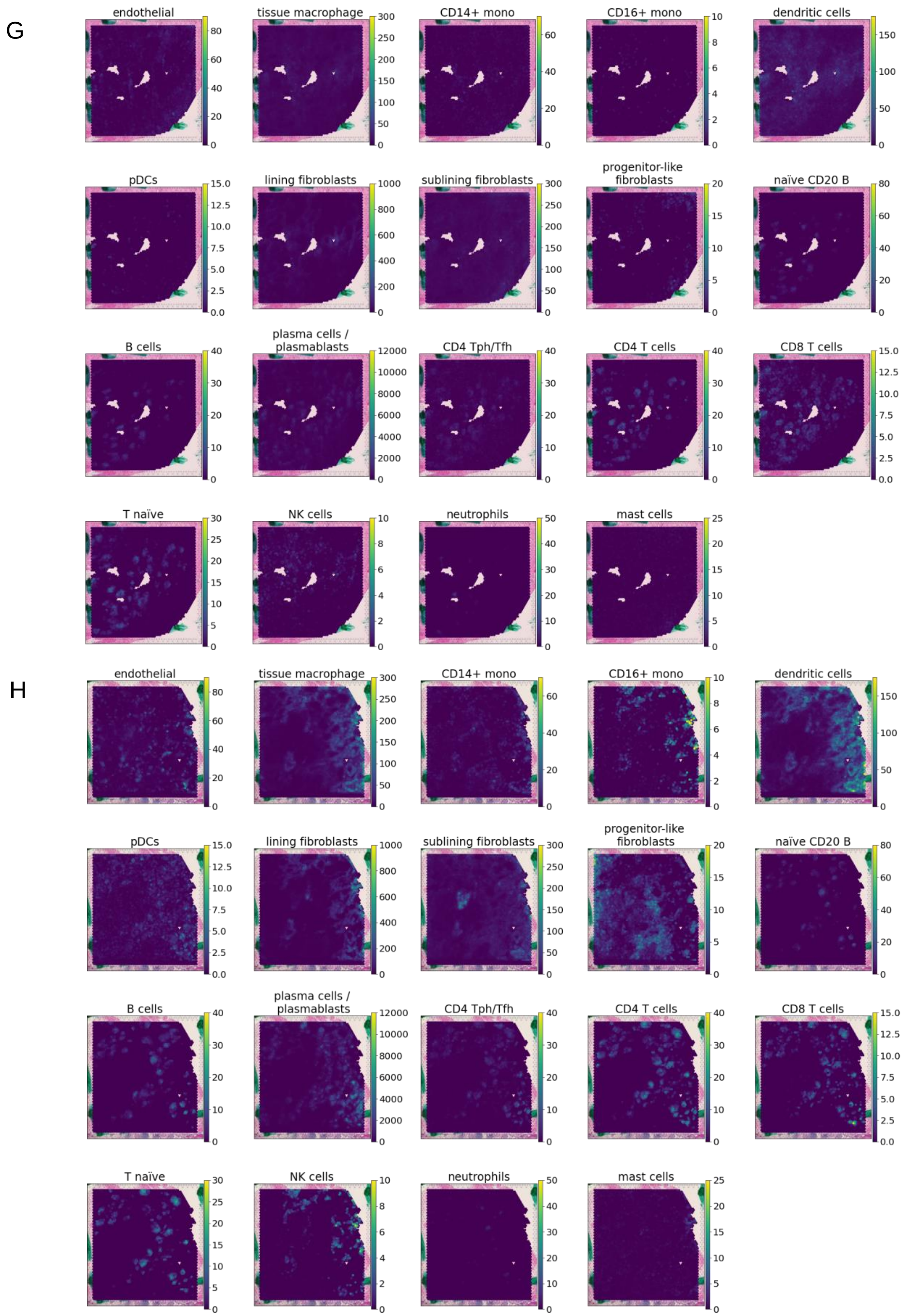


**Supplementary Figure 1:** Gene signature scoring of ST capture spots. Results of scoring of spots based on curated gene signatures of known cell types in RA, for each sample **(A)** RA1, **(B)** RA2A, **(C)** RA2B, **(D)** RA3, **(E)** RA4, **(F)** RA5A, **(G)** RA5B, **(H)** RA6.

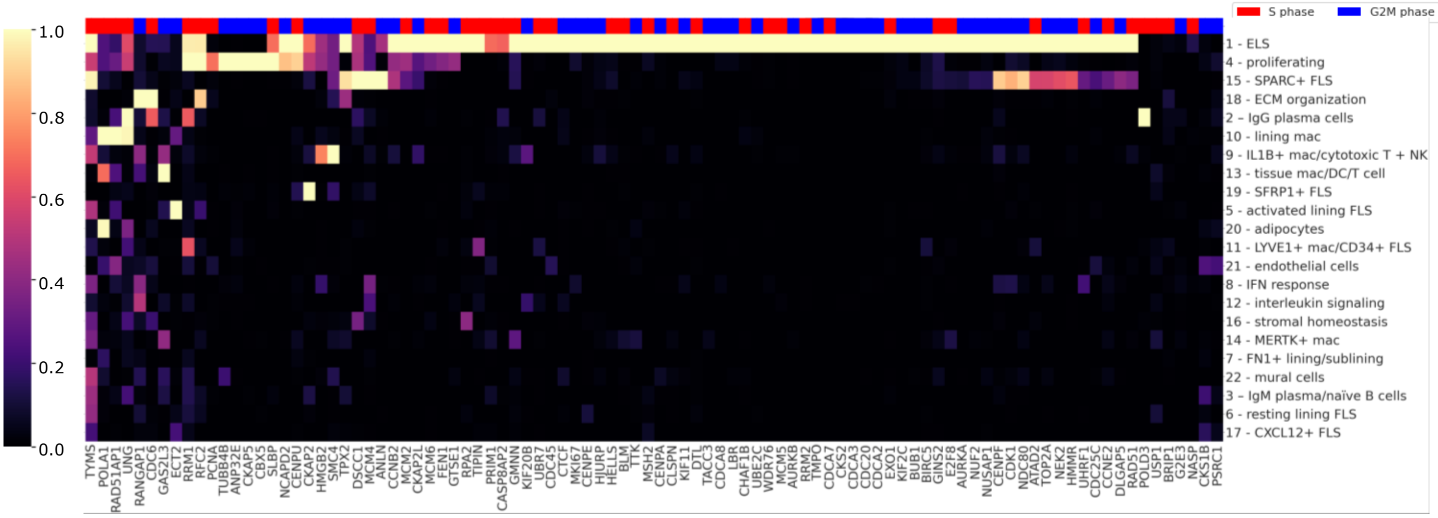


**Supplementary Figure 2:** Rankings of known proliferation markers in each topic. Min-max normalized KL-based scores for G2M (blue) and S phase (red) marker genes (obtained from the Scanpy analysis suite) across topics.

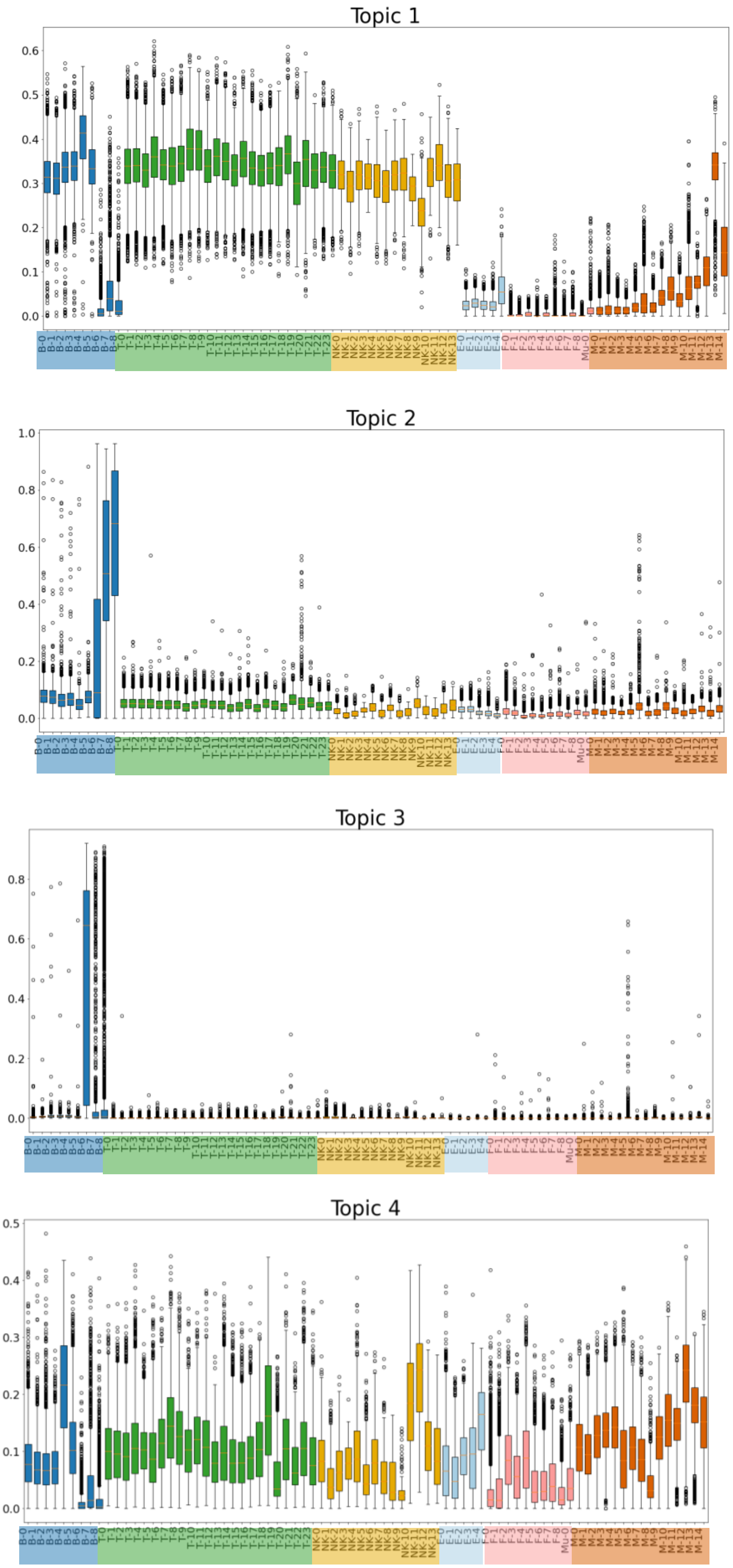


| Symbol | Cluster |
|---|---|
| B-0 | CD24+CD27+CD11b+ switched mem |
| B-1 | CD24++CD27+IgM+ unswitched mem |
| B-2 | IgM+IgD+TCL1A+ naive |
| B-3 | IgM+IgD+CD1c+ MZ-like |
| B-4 | AICDA+BCL6+ GC-like |
| B-5 | CD11c+LAMP1+ ABC |
| B-6 | IgM+ plasma |
| B-7 | HLA-DR+IgG+ plasmablast |
| B-8 | IgG1+IgG3+ plasma |
| T-0 | CD4+ IL7R+ memory |
| T-1 | CD4+ CD161+ memory |
| T-2 | CD4+ IL7R+CCR5+ memory |
| T-3 | CD4+ Tfh/Tph |
| T-4 | CD4+ naive |
| T-5 | CD4+ GZMK+ memory |
| T-6 | CD4+ memory |
| T-7 | CD4+ Tph |
| T-8 | CD4+ CD25-high Treg |
| T-9 | CD4+ CD25-low Treg |
| T-10 | CD4+ OX40+NR3C1+ |
| T-11 | CD4+ CD146+ memory |
| T-12 | CD4+ GNLY+ |
| T-13 | CD8+ GZMK/B+ memory |
| T-14 | CD8+ GZMK+ memory |
| T-15 | CD8+ GZMB+/TEMRA |
| T-16 | CD8+ CD45ROlow/naive |
| T-17 | CD8+ activated/NK-like |
| T-18 | Proliferating |
| T-19 | MT-high (low quality) |
| T-20 | CD38+ |
| T-21 | Innate-like |
| T-22 | Vdelta1 |
| T-23 | Vdelta2 |
| NK-0 | CD56dim CD16+ IFNG- |
| NK-1 | CD56dim CD16+ IFNG+CD160+ |
| NK-2 | CD56dim CD16+ IFNG+CD160- |
| NK-3 | CD56dim CD16+ GZMB- |
| NK-4 | CD56bright CD16- GZMA+CD160+ |
| NK-5 | CD56bright CD16- GZMA+CD69+ |
| NK-6 | CD56bright CD16- GNLY+ |
| NK-7 | CD56bright CD16- GNLY+CD69+ |
| NK-8 | CD56bright CD16- IFN response |
| NK-9 | MT-high |
| NK-10 | PCNA+ Proliferating |
| NK-11 | MKI67+ Proliferating |
| NK-12 | IL7R+ ILC |
| NK-13 | IL7R+CD161+ ILC |
| E-0 | SPARC+ capillary |
| E-1 | LIFR+ venular |
| E-2 | ICAM1+ venular |
| E-3 | NOTCH4+ arteriolar |
| E-4 | Lymphatic |
| F-0 | PRG4+ CLIC5+ lining |
| F-1 | PRG4+ lining |
| F-2 | CD34+ sublining |
| F-3 | POSTN+ sublining |
| F-4 | DKK3+ sublining |
| F-5 | CD74-hi sublining |
| F-6 | CXCL12+ SFRP1+ sublining |
| F-7 | NOTCH3+ sublining |
| F-8 | RSPO3+ intermediate |
| Mu-0 | Mural |
| M-0 | MERTK+ SELENOP+ LYVE1+ |
| M-1 | MERTK+ SELENOP+ LYVE1- |
| M-2 | MERTK+ S100A8+ |
| M-3 | MERTK+ HBEGF+ |
| M-4 | SPP1+ |
| M-5 | C1QA+ |
| M-6 | STAT1+ CXCL10+ |
| M-7 | IL1B+ FCN1+ |
| M-8 | PLCG2+ |
| M-9 | DC3 |
| M-10 | DC2 |
| M-11 | DC4 |
| M-12 | DC1 |
| M-13 | pDC |
| M-14 | LAMP3+ |

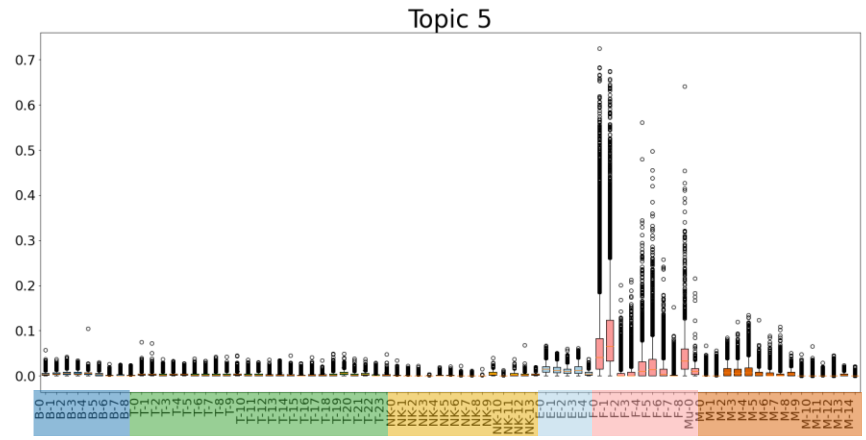


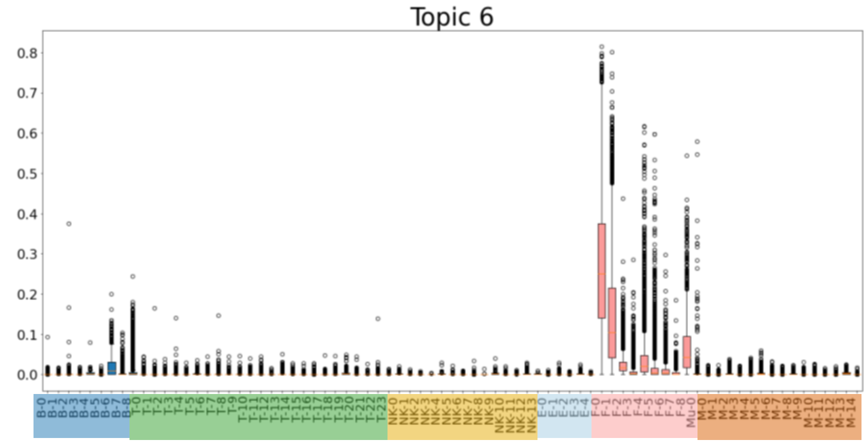


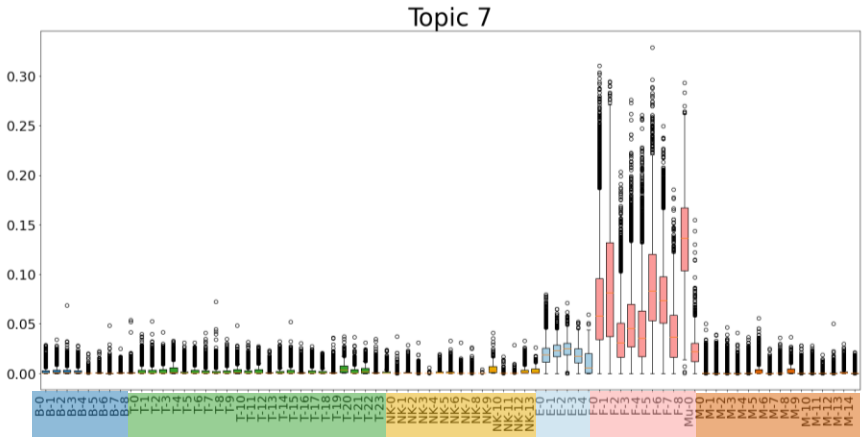


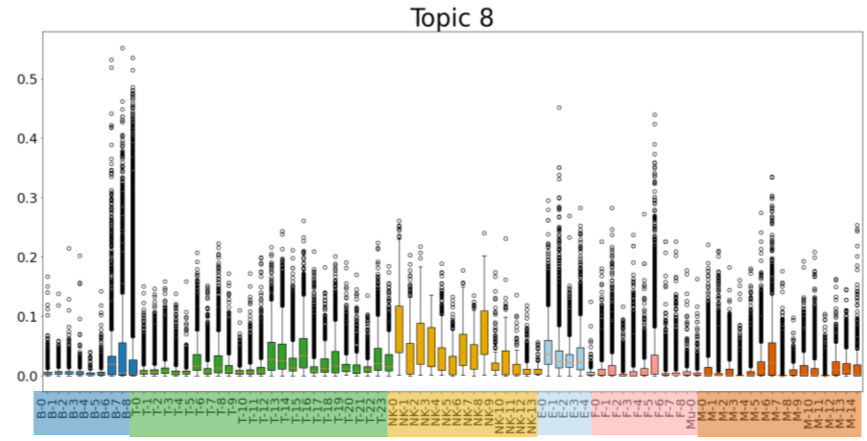


| Symbol | Cluster |
|---|---|
| B-0 | CD24+CD27+CD11b+ switched mem |
| B-1 | CD24++CD27+IgM+ unswitched mem |
| B-2 | IgM+IgD+TCL1A+ naive |
| B-3 | IgM+IgD+CD1c+ MZ-like |
| B-4 | AICDA+BCL6+ GC-like |
| B-5 | CD11c+LAMP1+ ABC |
| B-6 | IgM+ plasma |
| B-7 | HLA-DR+IgG+ plasmablast |
| B-8 | IgG1+IgG3+ plasma |
| T-0 | CD4+ IL7R+ memory |
| T-1 | CD4+ CD161+ memory |
| T-2 | CD4+ IL7R+CCR5+ memory |
| T-3 | CD4+ Tfh/Tph |
| T-4 | CD4+ naive |
| T-5 | CD4+ GZMK+ memory |
| T-6 | CD4+ memory |
| T-7 | CD4+ Tph |
| T-8 | CD4+ CD25-high Treg |
| T-9 | CD4+ CD25-low Treg |
| T-10 | CD4+ OX40+NR3C1+ |
| T-11 | CD4+ CD146+ memory |
| T-12 | CD4+ GNLY+ |
| T-13 | CD8+ GZMK/B+ memory |
| T-14 | CD8+ GZMK+ memory |
| T-15 | CD8+ GZMB+/TEMRA |
| T-16 | CD8+ CD45ROlow/naive |
| T-17 | CD8+ activated/NK-like |
| T-18 | Proliferating |
| T-19 | MT-high (low quality) |
| T-20 | CD38+ |
| T-21 | Innate-like |
| T-22 | Vdelta1 |
| T-23 | Vdelta2 |
| NK-0 | CD56dim CD16+ IFNG- |
| NK-1 | CD56dim CD16+ IFNG+CD160+ |
| NK-2 | CD56dim CD16+ IFNG+CD160- |
| NK-3 | CD56dim CD16+ GZMB- |
| NK-4 | CD56bright CD16- GZMA+CD160+ |
| NK-5 | CD56bright CD16- GZMA+CD69+ |
| NK-6 | CD56bright CD16- GNLY+ |
| NK-7 | CD56bright CD16- GNLY+CD69+ |
| NK-8 | CD56bright CD16- IFN response |
| NK-9 | MT-high |
| NK-10 | PCNA+ Proliferating |
| NK-11 | MKI67+ Proliferating |
| NK-12 | IL7R+ ILC |
| NK-13 | IL7R+CD161+ ILC |
| E-0 | SPARC+ capillary |
| E-1 | LIFR+ venular |
| E-2 | ICAM1+ venular |
| E-3 | NOTCH4+ arteriolar |
| E-4 | Lymphatic |
| F-0 | PRG4+ CLIC5+ lining |
| F-1 | PRG4+ lining |
| F-2 | CD34+ sublining |
| F-3 | POSTN+ sublining |
| F-4 | DKK3+ sublining |
| F-5 | CD74-hi sublining |
| F-6 | CXCL12+ SFRP1+ sublining |
| F-7 | NOTCH3+ sublining |
| F-8 | RSPO3+ intermediate |
| Mu-0 | Mural |
| M-0 | MERTK+ SELENOP+ LYVE1+ |
| M-1 | MERTK+ SELENOP+ LYVE1- |
| M-2 | MERTK+ S100A8+ |
| M-3 | MERTK+ HBEGF+ |
| M-4 | SPP1+ |
| M-5 | C1QA+ |
| M-6 | STAT1+ CXCL10+ |
| M-7 | IL1B+ FCN1+ |
| M-8 | PLCG2+ |
| M-9 | DC3 |
| M-10 | DC2 |
| M-11 | DC4 |
| M-12 | DC1 |
| M-13 | pDC |
| M-14 | LAMP3+ |

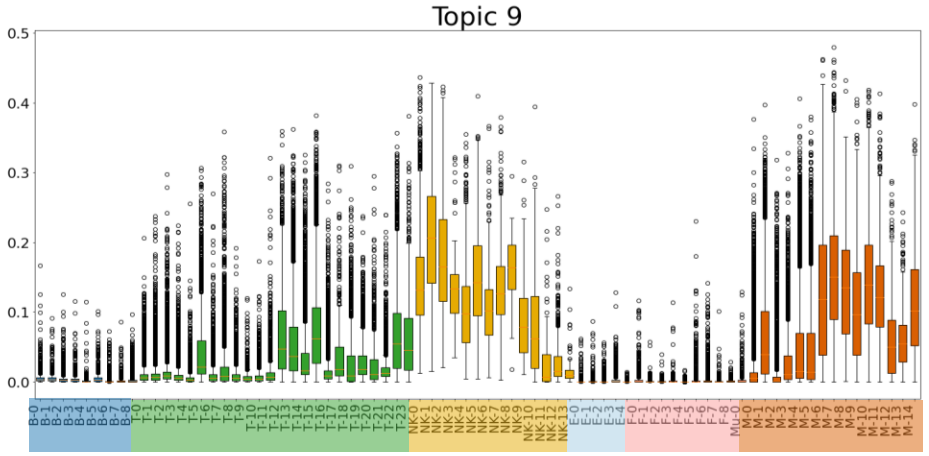


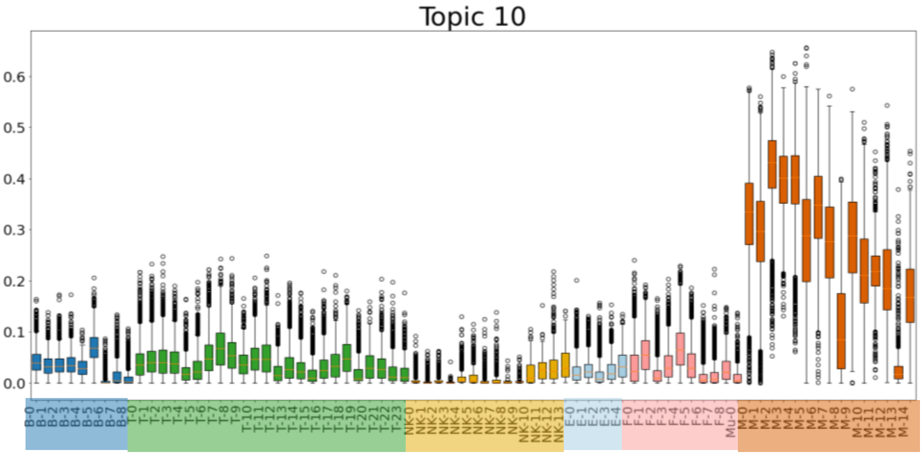


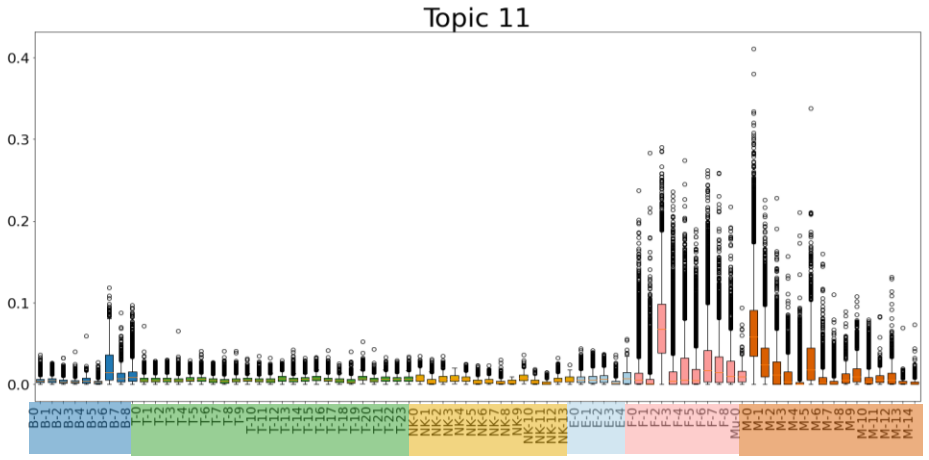


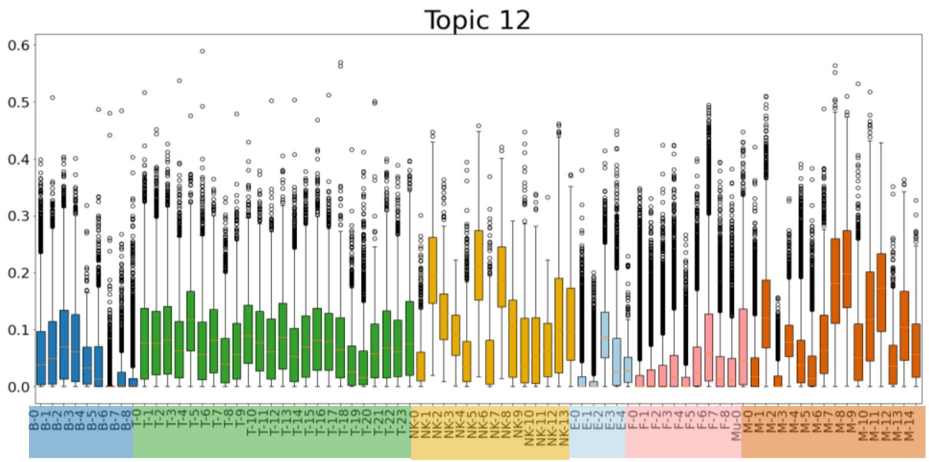


| Symbol | Cluster |
| --- | --- |
| B-0 | CD24+CD27+CD11b+ switched mem |
| B-1 | CD24++CD27+IgM+ unswitched mem |
| B-2 | IgM+IgD+TCL1A+ naive |
| B-3 | IgM+IgD+CD1c+ MZ-like |
| B-4 | AICDA+BCL6+ GC-like |
| B-5 | CD11c+LAMP1+ ABC |
| B-6 | IgM+ plasma |
| B-7 | HLA-DR+IgG+ plasmablast |
| B-8 | IgG1+IgG3+ plasma |
| T-0 | CD4+ IL7R+ memory |
| T-1 | CD4+ CD161+ memory |
| T-2 | CD4+ IL7R+CCR5+ memory |
| T-3 | CD4+ Tfh/Tph |
| T-4 | CD4+ naive |
| T-5 | CD4+ GZMK+ memory |
| T-6 | CD4+ memory |
| T-7 | CD4+ Tph |
| T-8 | CD4+ CD25-high Treg |
| T-9 | CD4+ CD25-low Treg |
| T-10 | CD4+ OX40+NR3C1+ |
| T-11 | CD4+ CD146+ memory |
| T-12 | CD4+ GNLY+ |
| T-13 | CD8+ GZMK/B+ memory |
| T-14 | CD8+ GZMK+ memory |
| T-15 | CD8+ GZMB+/TEMRA |
| T-16 | CD8+ CD45ROlow/naive |
| T-17 | CD8+ activated/NK-like |
| T-18 | Proliferating |
| T-19 | MT-high (low quality) |
| T-20 | CD38+ |
| T-21 | Innate-like |
| T-22 | Vdelta1 |
| T-23 | Vdelta2 |
| NK-0 | CD56dim CD16+ IFNG- |
| NK-1 | CD56dim CD16+ IFNG+CD160+ |
| NK-2 | CD56dim CD16+ IFNG+CD160- |
| NK-3 | CD56dim CD16+ GZMB- |
| NK-4 | CD56bright CD16- GZMA+CD160+ |
| NK-5 | CD56bright CD16- GZMA+CD69+ |
| NK-6 | CD56bright CD16- GNLY+ |
| NK-7 | CD56bright CD16- GNLY+CD69+ |
| NK-8 | CD56bright CD16- IFN response |
| NK-9 | MT-high |
| NK-10 | PCNA+ Proliferating |
| NK-11 | MKI67+ Proliferating |
| NK-12 | IL7R+ ILC |
| NK-13 | IL7R+CD161+ ILC |
| E-0 | SPARC+ capillary |
| E-1 | LIFR+ venular |
| E-2 | ICAM1+ venular |
| E-3 | NOTCH4+ arteriolar |
| E-4 | Lymphatic |
| F-0 | PRG4+ CLIC5+ lining |
| F-1 | PRG4+ lining |
| F-2 | CD34+ sublining |
| F-3 | POSTN+ sublining |
| F-4 | DKK3+ sublining |
| F-5 | CD74-hi sublining |
| F-6 | CXCL12+ SFRP1+ sublining |
| F-7 | NOTCH3+ sublining |
| F-8 | RSPO3+ intermediate |
| Mu-0 | Mural |
| M-0 | MERTK+ SELENOP+ LYVE1+ |
| M-1 | MERTK+ SELENOP+ LYVE1- |
| M-2 | MERTK+ S100A8+ |
| M-3 | MERTK+ HBEGF+ |
| M-4 | SPP1+ |
| M-5 | C1QA+ |
| M-6 | STAT1+ CXCL10+ |
| M-7 | IL1B+ FCN1+ |
| M-8 | PLCG2+ |
| M-9 | DC3 |
| M-10 | DC2 |
| M-11 | DC4 |
| M-12 | DC1 |
| M-13 | pDC |
| M-14 | LAMP3+ |

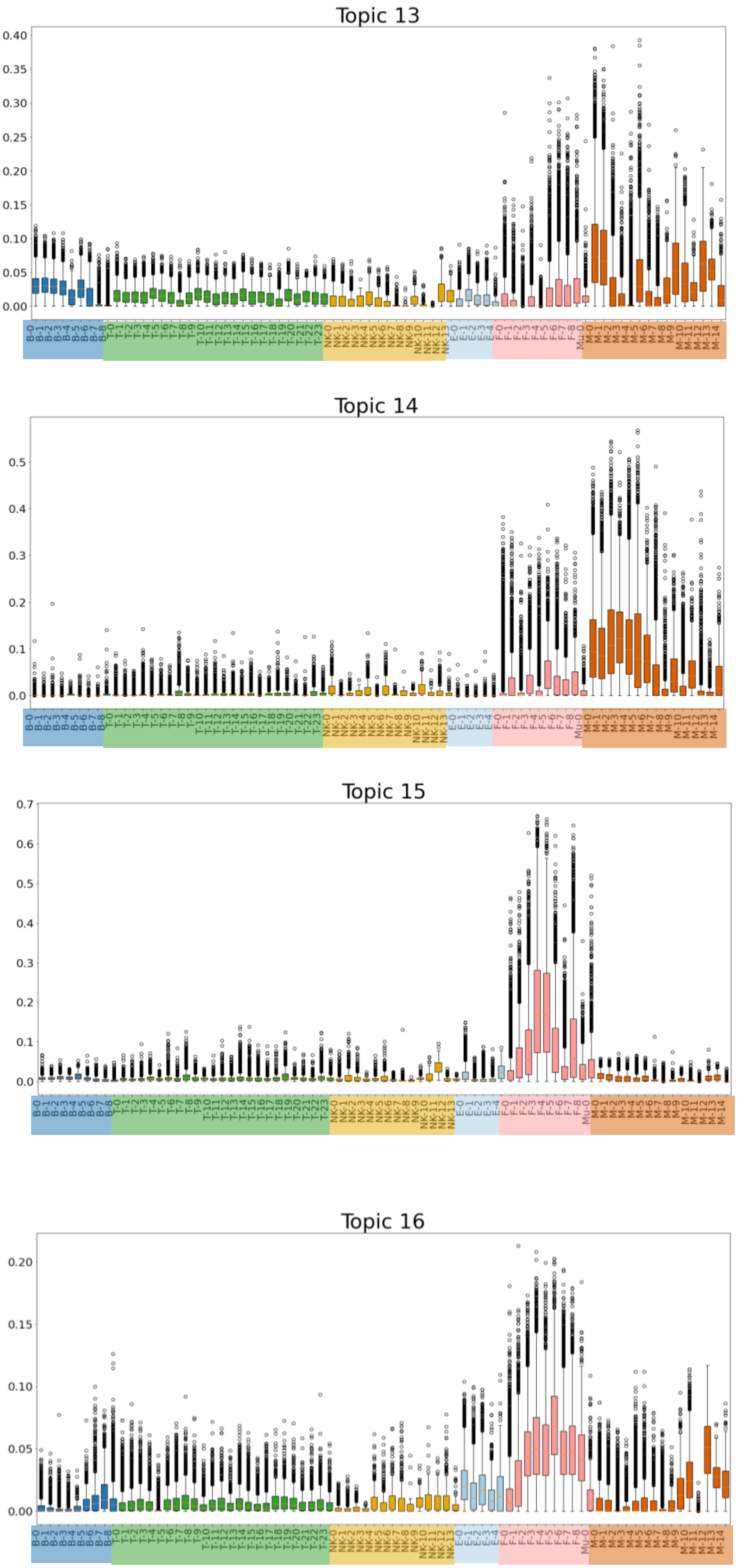


| Symbol | Cluster |
|---|---|
| B-0 | CD24+CD27+CD11b+ switched mem |
| B-1 | CD24++CD27+IgM+ unswitched mem |
| B-2 | IgM+IgD+TCL1A+ naive |
| B-3 | IgM+IgD+CD1c+ MZ-like |
| B-4 | AICDA+BCL6+ GC-like |
| B-5 | CD11c+LAMP1+ ABC |
| B-6 | IgM+ plasma |
| B-7 | HLA-DR+IgG+ plasmablast |
| B-8 | IgG1+IgG3+ plasma |
| T-0 | CD4+ IL7R+ memory |
| T-1 | CD4+ CD161+ memory |
| T-2 | CD4+ IL7R+CCR5+ memory |
| T-3 | CD4+ Tfh/Tph |
| T-4 | CD4+ naive |
| T-5 | CD4+ GZMK+ memory |
| T-6 | CD4+ memory |
| T-7 | CD4+ Tph |
| T-8 | CD4+ CD25-high Treg |
| T-9 | CD4+ CD25-low Treg |
| T-10 | CD4+ OX40+NR3C1+ |
| T-11 | CD4+ CD146+ memory |
| T-12 | CD4+ GNLY+ |
| T-13 | CD8+ GZMK/B+ memory |
| T-14 | CD8+ GZMK+ memory |
| T-15 | CD8+ GZMB+/TEMRA |
| T-16 | CD8+ CD45ROlow/naive |
| T-17 | CD8+ activated/NK-like |
| T-18 | Proliferating |
| T-19 | MT-high (low quality) |
| T-20 | CD38+ |
| T-21 | Innate-like |
| T-22 | Vdelta1 |
| T-23 | Vdelta2 |
| NK-0 | CD56dim CD16+ IFNG- |
| NK-1 | CD56dim CD16+ IFNG+CD160+ |
| NK-2 | CD56dim CD16+ IFNG+CD160- |
| NK-3 | CD56dim CD16+ GZMB- |
| NK-4 | CD56bright CD16- GZMA+CD160+ |
| NK-5 | CD56bright CD16- GZMA+CD69+ |
| NK-6 | CD56bright CD16- GNLY+ |
| NK-7 | CD56bright CD16- GNLY+CD69+ |
| NK-8 | CD56bright CD16- IFN response |
| NK-9 | MT-high |
| NK-10 | PCNA+ Proliferating |
| NK-11 | MKI67+ Proliferating |
| NK-12 | IL7R+ ILC |
| NK-13 | IL7R+CD161+ ILC |
| E-0 | SPARC+ capillary |
| E-1 | LIFR+ venular |
| E-2 | ICAM1+ venular |
| E-3 | NOTCH4+ arteriolar |
| E-4 | Lymphatic |
| F-0 | PRG4+ CLIC5+ lining |
| F-1 | PRG4+ lining |
| F-2 | CD34+ sublining |
| F-3 | POSTN+ sublining |
| F-4 | DKK3+ sublining |
| F-5 | CD74-hi sublining |
| F-6 | CXCL12+ SFRP1+ sublining |
| F-7 | NOTCH3+ sublining |
| F-8 | RSPO3+ intermediate |
| Mu-0 | Mural |
| M-0 | MERTK+ SELENOP+ LYVE1+ |
| M-1 | MERTK+ SELENOP+ LYVE1- |
| M-2 | MERTK+ S100A8+ |
| M-3 | MERTK+ HBEGF+ |
| M-4 | SPP1+ |
| M-5 | C1QA+ |
| M-6 | STAT1+ CXCL10+ |
| M-7 | IL1B+ FCN1+ |
| M-8 | PLCG2+ |
| M-9 | DC3 |
| M-10 | DC2 |
| M-11 | DC4 |
| M-12 | DC1 |
| M-13 | pDC |
| M-14 | LAMP3+ |

### Topic 17

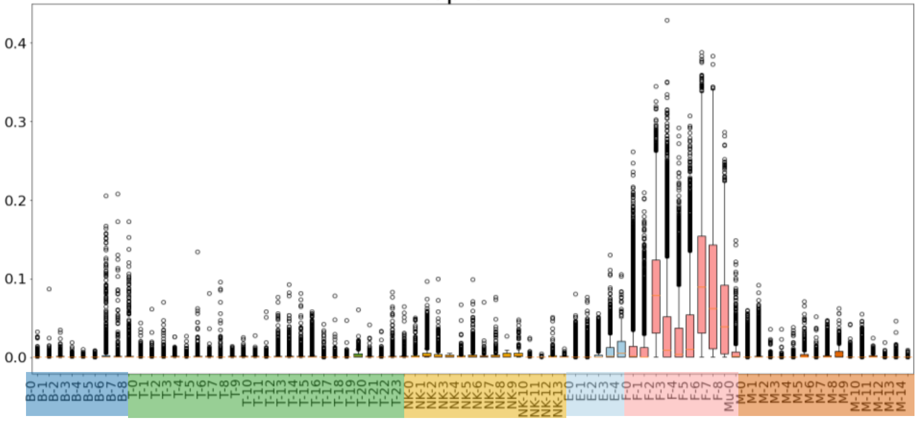


### Topic 18

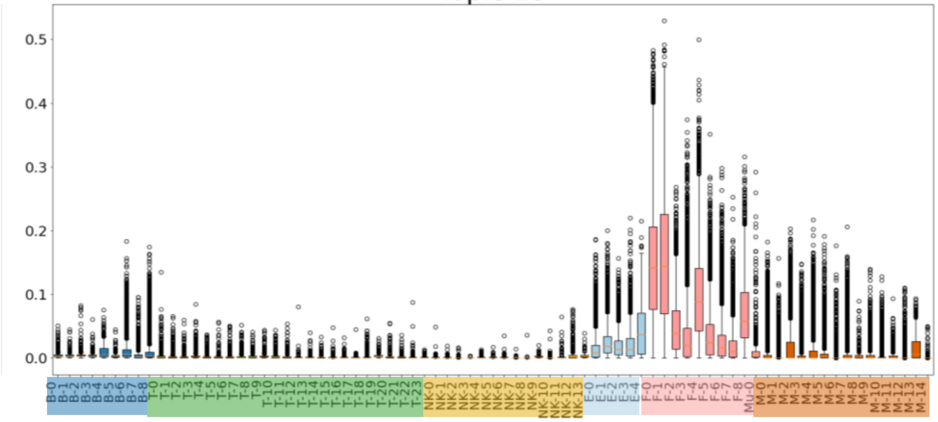


### Topic 19

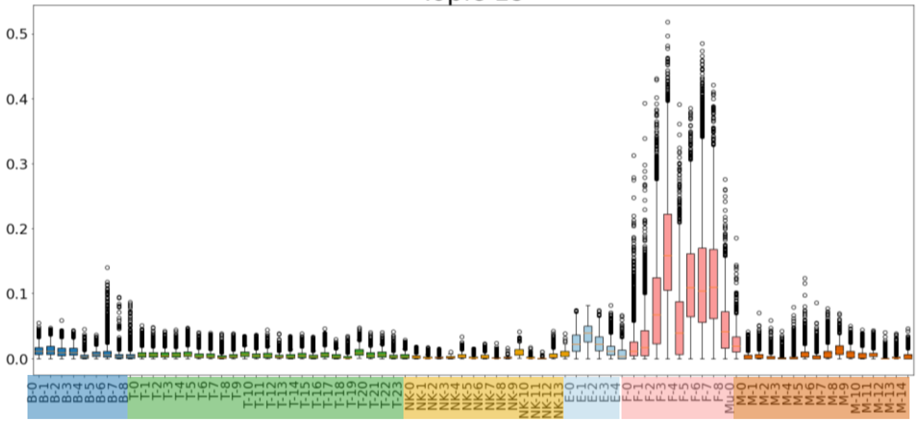


### Topic 20

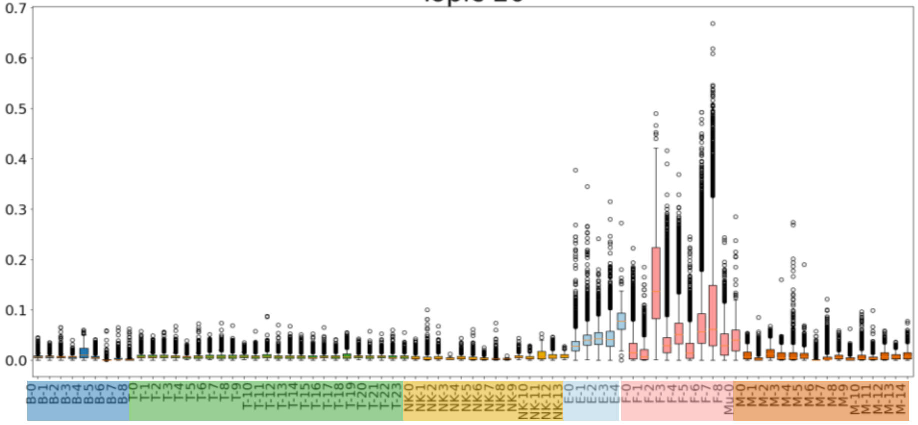


| Symbol | Cluster |
|---|---|
| B-0 | CD24+CD27+CD11b+ switched mem |
| B-1 | CD24++CD27+IgM+ unswitched mem |
| B-2 | IgM+IgD+TCL1A+ naive |
| B-3 | IgM+IgD+CD1c+ MZ-like |
| B-4 | AICDA+BCL6+ GC-like |
| B-5 | CD11c+LAMP1+ ABC |
| B-6 | IgM+ plasma |
| B-7 | HLA-DR+IgG+ plasmablast |
| B-8 | IgG1+IgG3+ plasma |
| T-0 | CD4+ IL7R+ memory |
| T-1 | CD4+ CD161+ memory |
| T-2 | CD4+ IL7R+CCR5+ memory |
| T-3 | CD4+ Tfh/Tph |
| T-4 | CD4+ naive |
| T-5 | CD4+ GZMK+ memory |
| T-6 | CD4+ memory |
| T-7 | CD4+ Tph |
| T-8 | CD4+ CD25-high Treg |
| T-9 | CD4+ CD25-low Treg |
| T-10 | CD4+ OX40+NR3C1+ |
| T-11 | CD4+ CD146+ memory |
| T-12 | CD4+ GNLY+ |
| T-13 | CD8+ GZMK/B+ memory |
| T-14 | CD8+ GZMK+ memory |
| T-15 | CD8+ GZMB+/TEMRA |
| T-16 | CD8+ CD45ROlow/naive |
| T-17 | CD8+ activated/NK-like |
| T-18 | Proliferating |
| T-19 | MT-high (low quality) |
| T-20 | CD38+ |
| T-21 | Innate-like |
| T-22 | Vdelta1 |
| T-23 | Vdelta2 |
| NK-0 | CD56dim CD16+ IFNG- |
| NK-1 | CD56dim CD16+ IFNG+CD160+ |
| NK-2 | CD56dim CD16+ IFNG+CD160- |
| NK-3 | CD56dim CD16+ GZMB- |
| NK-4 | CD56bright CD16- GZMA+CD160+ |
| NK-5 | CD56bright CD16- GZMA+CD69+ |
| NK-6 | CD56bright CD16- GNLY+ |
| NK-7 | CD56bright CD16- GNLY+CD69+ |
| NK-8 | CD56bright CD16- IFN response |
| NK-9 | MT-high |
| NK-10 | PCNA+ Proliferating |
| NK-11 | MKI67+ Proliferating |
| NK-12 | IL7R+ ILC |
| NK-13 | IL7R+CD161+ ILC |
| E-0 | SPARC+ capillary |
| E-1 | LIFR+ venular |
| E-2 | ICAM1+ venular |
| E-3 | NOTCH4+ arteriolar |
| E-4 | Lymphatic |
| F-0 | PRG4+ CLIC5+ lining |
| F-1 | PRG4+ lining |
| F-2 | CD34+ sublining |
| F-3 | POSTN+ sublining |
| F-4 | DKK3+ sublining |
| F-5 | CD74-hi sublining |
| F-6 | CXCL12+ SFRP1+ sublining |
| F-7 | NOTCH3+ sublining |
| F-8 | RSPO3+ intermediate |
| Mu-0 | Mural |
| M-0 | MERTK+ SELENOP+ LYVE1+ |
| M-1 | MERTK+ SELENOP+ LYVE1- |
| M-2 | MERTK+ S100A8+ |
| M-3 | MERTK+ HBEGF+ |
| M-4 | SPP1+ |
| M-5 | C1QA+ |
| M-6 | STAT1+ CXCL10+ |
| M-7 | IL1B+ FCN1+ |
| M-8 | PLCG2+ |
| M-9 | DC3 |
| M-10 | DC2 |
| M-11 | DC4 |
| M-12 | DC1 |
| M-13 | pDC |
| M-14 | LAMP3+ |

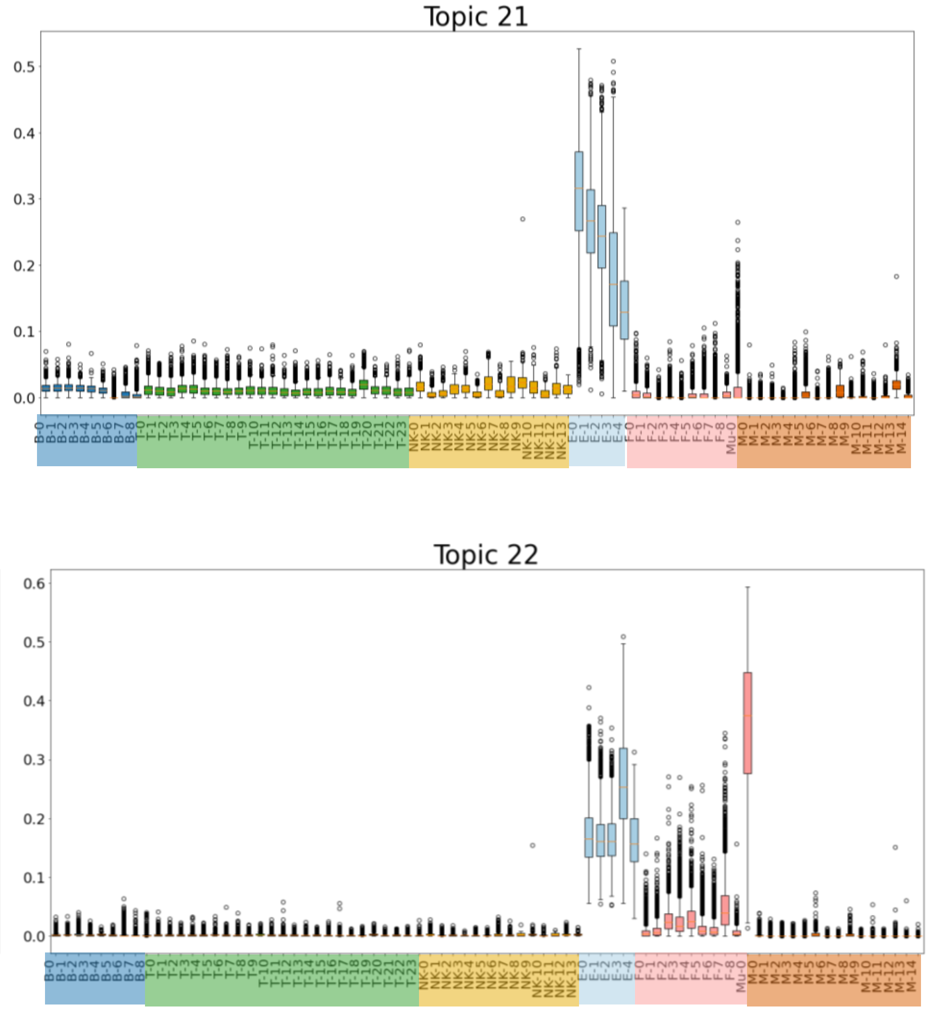


| Symbol | Cluster |
|---|---|
| B-0 | CD24+CD27+CD11b+ switched mem |
| B-1 | CD24++CD27+IgM+ unswitched mem |
| B-2 | IgM+IgD+TCL1A+ naive |
| B-3 | IgM+IgD+CD1c+ MZ-like |
| B-4 | AICDA+BCL6+ GC-like |
| B-5 | CD11c+LAMP1+ ABC |
| B-6 | IgM+ plasma |
| B-7 | HLA-DR+IgG+ plasmablast |
| B-8 | IgG1+IgG3+ plasma |
| T-0 | CD4+ IL7R+ memory |
| T-1 | CD4+ CD161+ memory |
| T-2 | CD4+ IL7R+CCR5+ memory |
| T-3 | CD4+ Tfh/Tph |
| T-4 | CD4+ naive |
| T-5 | CD4+ GZMK+ memory |
| T-6 | CD4+ memory |
| T-7 | CD4+ Tph |
| T-8 | CD4+ CD25-high Treg |
| T-9 | CD4+ CD25-low Treg |
| T-10 | CD4+ OX40+NR3C1+ |
| T-11 | CD4+ CD146+ memory |
| T-12 | CD4+ GNLY+ |
| T-13 | CD8+ GZMK/B+ memory |
| T-14 | CD8+ GZMK+ memory |
| T-15 | CD8+ GZMB+/TEMRA |
| T-16 | CD8+ CD45ROlow/naive |
| T-17 | CD8+ activated/NK-like |
| T-18 | Proliferating |
| T-19 | MT-high (low quality) |
| T-20 | CD38+ |
| T-21 | Innate-like |
| T-22 | Vdelta1 |
| T-23 | Vdelta2 |
| NK-0 | CD56dim CD16+ IFNG- |
| NK-1 | CD56dim CD16+ IFNG+CD160+ |
| NK-2 | CD56dim CD16+ IFNG+CD160- |
| NK-3 | CD56dim CD16+ GZMB- |
| NK-4 | CD56bright CD16- GZMA+CD160+ |
| NK-5 | CD56bright CD16- GZMA+CD69+ |
| NK-6 | CD56bright CD16- GNLY+ |
| NK-7 | CD56bright CD16- GNLY+CD69+ |
| NK-8 | CD56bright CD16- IFN response |
| NK-9 | MT-high |
| NK-10 | PCNA+ Proliferating |
| NK-11 | MKI67+ Proliferating |
| NK-12 | IL7R+ ILC |
| NK-13 | IL7R+CD161+ ILC |
| E-0 | SPARC+ capillary |
| E-1 | LIFR+ venular |
| E-2 | ICAM1+ venular |
| E-3 | NOTCH4+ arteriolar |
| E-4 | Lymphatic |
| F-0 | PRG4+ CLIC5+ lining |
| F-1 | PRG4+ lining |
| F-2 | CD34+ sublining |
| F-3 | POSTN+ sublining |
| F-4 | DKK3+ sublining |
| F-5 | CD74-hi sublining |
| F-6 | CXCL12+ SFRP1+ sublining |
| F-7 | NOTCH3+ sublining |
| F-8 | RSPO3+ intermediate |
| Mu-0 | Mural |
| M-0 | MERTK+ SELENOP+ LYVE1+ |
| M-1 | MERTK+ SELENOP+ LYVE1- |
| M-2 | MERTK+ S100A8+ |
| M-3 | MERTK+ HBEGF+ |
| M-4 | SPP1+ |
| M-5 | C1QA+ |
| M-6 | STAT1+ CXCL10+ |
| M-7 | IL1B+ FCN1+ |
| M-8 | PLCG2+ |
| M-9 | DC3 |
| M-10 | DC2 |
| M-11 | DC4 |
| M-12 | DC1 |
| M-13 | pDC |
| M-14 | LAMP3+ |

**Supplementary Figure 3:** Application of the trained DeepTopics model to a published scRNA-seq atlas[25] for RA. Each box represents the interquartile range of posterior probabilities for the topic across cells belonging to the cluster.

A

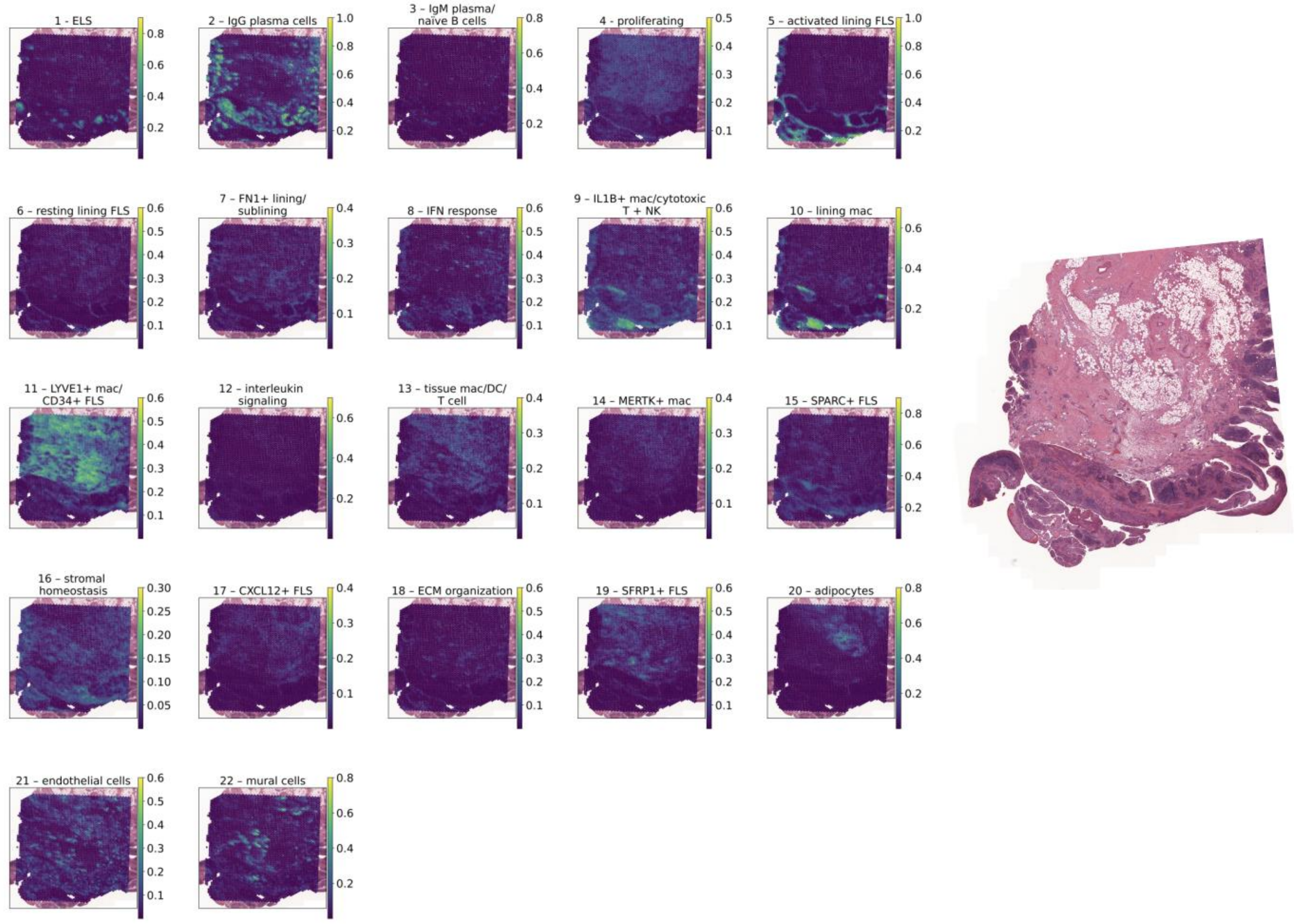
1 - ELS
2 – IgG plasma cells
3 – IgM plasma/ naive B cells
4 - proliferating
5 – activated lining FLS
6 – resting lining FLS
7 – FN1+ lining/ sublining
8 – IFN response
9 – IL1B+ mac/cytotoxic T + NK
10 – lining mac
11 – LYVE1+ mac/ CD34+ FLS
12 – interleukin signaling
13 – tissue mac/DC/ T cell
14 – MERTK+ mac
15 – SPARC+ FLS
16 – stromal homeostasis
17 – CXCL12+ FLS
18 – ECM organization
19 – SFRP1+ FLS
20 – adipocytes
21 – endothelial cells
22 – mural cells

B

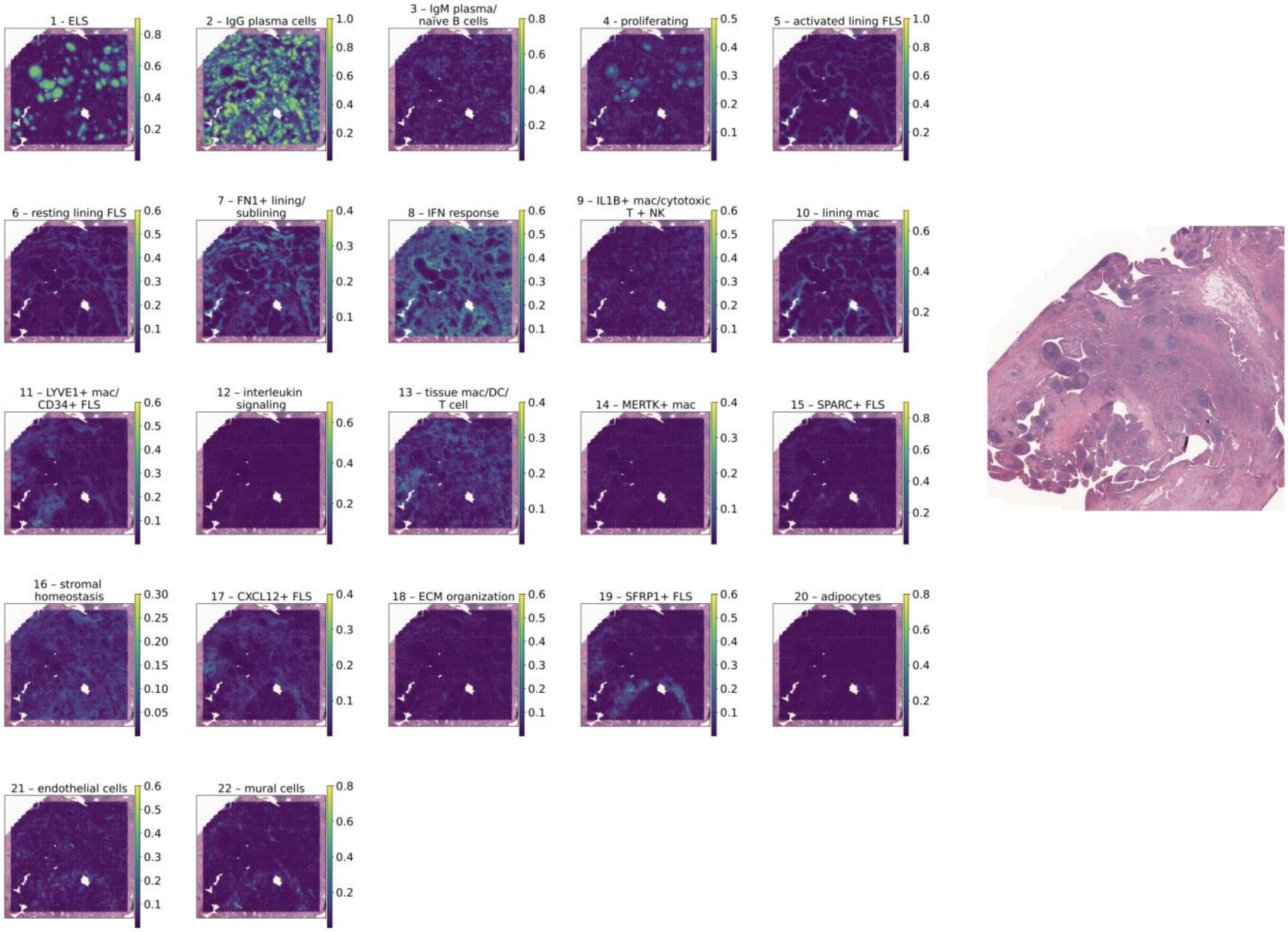
1 - ELS
2 – IgG plasma cells
3 – IgM plasma/ naive B cells
4 - proliferating
5 – activated lining FLS
6 – resting lining FLS
7 – FN1+ lining/ sublining
8 – IFN response
9 – IL1B+ mac/cytotoxic T + NK
10 – lining mac
11 – LYVE1+ mac/ CD34+ FLS
12 – interleukin signaling
13 – tissue mac/DC/ T cell
14 – MERTK+ mac
15 – SPARC+ FLS
16 – stromal homeostasis
17 – CXCL12+ FLS
18 – ECM organization
19 – SFRP1+ FLS
20 – adipocytes
21 – endothelial cells
22 – mural cells

C

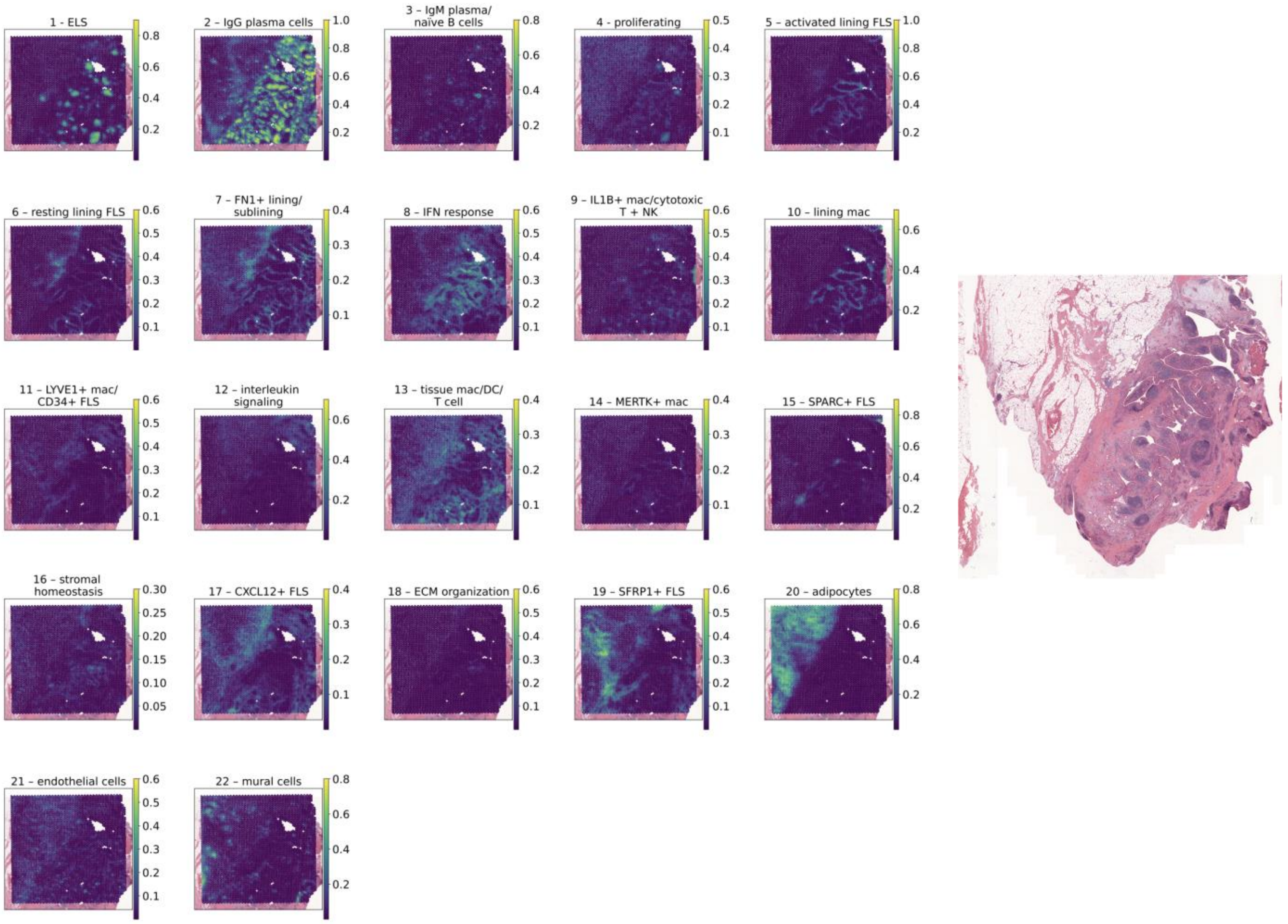
1 - ELS
2 – IgG plasma cells
3 – IgM plasma/ naïve B cells
4 - proliferating
5 – activated lining FLS
6 – resting lining FLS
7 – FN1+ lining/ sublining
8 – IFN response
9 – IL1B+ mac/cytotoxic T + NK
10 – lining mac
11 – LYVE1+ mac/ CD34+ FLS
12 – interleukin signaling
13 – tissue mac/DC/ T cell
14 – MERTK+ mac
15 – SPARC+ FLS
16 – stromal homeostasis
17 – CXCL12+ FLS
18 – ECM organization
19 – SFRP1+ FLS
20 – adipocytes
21 – endothelial cells
22 – mural cells

D

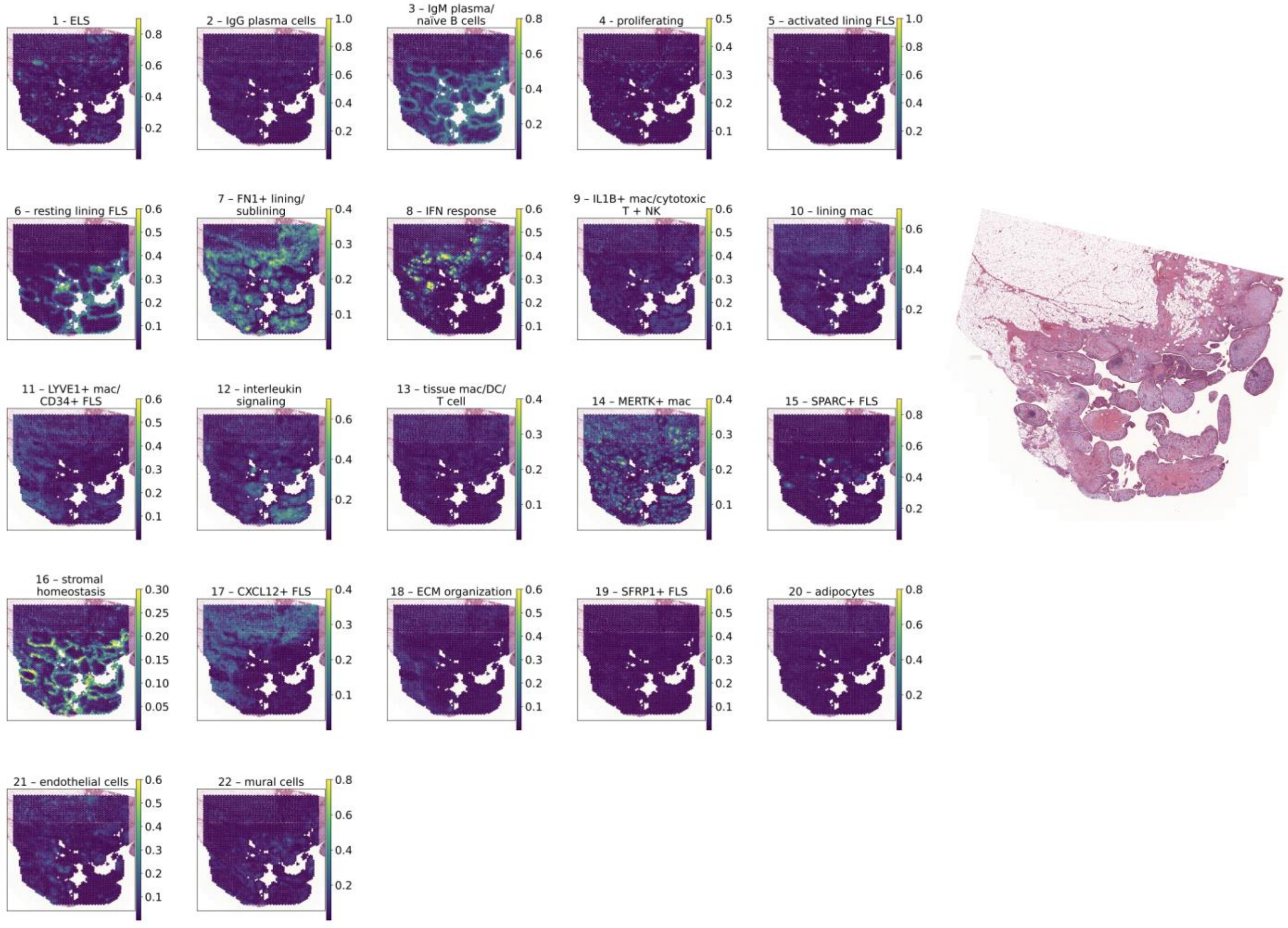
1 - ELS
2 – IgG plasma cells
3 – IgM plasma/ naïve B cells
4 - proliferating
5 – activated lining FLS
6 – resting lining FLS
7 – FN1+ lining/ sublining
8 – IFN response
9 – IL1B+ mac/cytotoxic T + NK
10 – lining mac
11 – LYVE1+ mac/ CD34+ FLS
12 – interleukin signaling
13 – tissue mac/DC/ T cell
14 – MERTK+ mac
15 – SPARC+ FLS
16 – stromal homeostasis
17 – CXCL12+ FLS
18 – ECM organization
19 – SFRP1+ FLS
20 – adipocytes
21 – endothelial cells
22 – mural cells

E

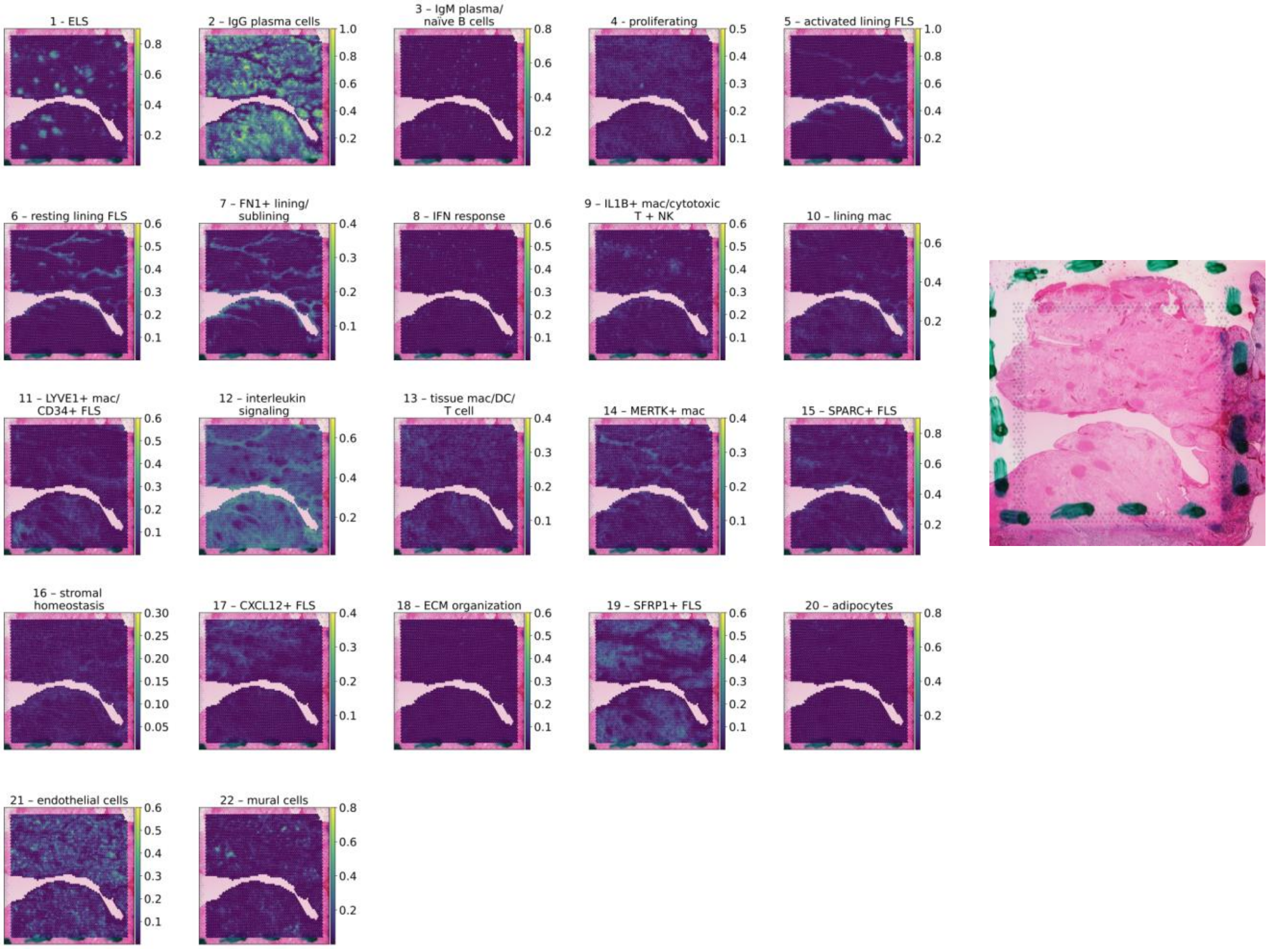
1 - ELS
2 - IgG plasma cells
3 – IgM plasma/ naive B cells
4 - proliferating
5 - activated lining FLS
6 – resting lining FLS
7 – FN1+ lining/ sublining
8 – IFN response
9 – IL1B+ mac/cytotoxic T + NK
10 – lining mac
11 – LYVE1+ mac/ CD34+ FLS
12 – interleukin signaling
13 – tissue mac/DC/ T cell
14 – MERTK+ mac
15 – SPARC+ FLS
16 – stromal homeostasis
17 – CXCL12+ FLS
18 – ECM organization
19 – SFRP1+ FLS
20 – adipocytes
21 – endothelial cells
22 – mural cells

F

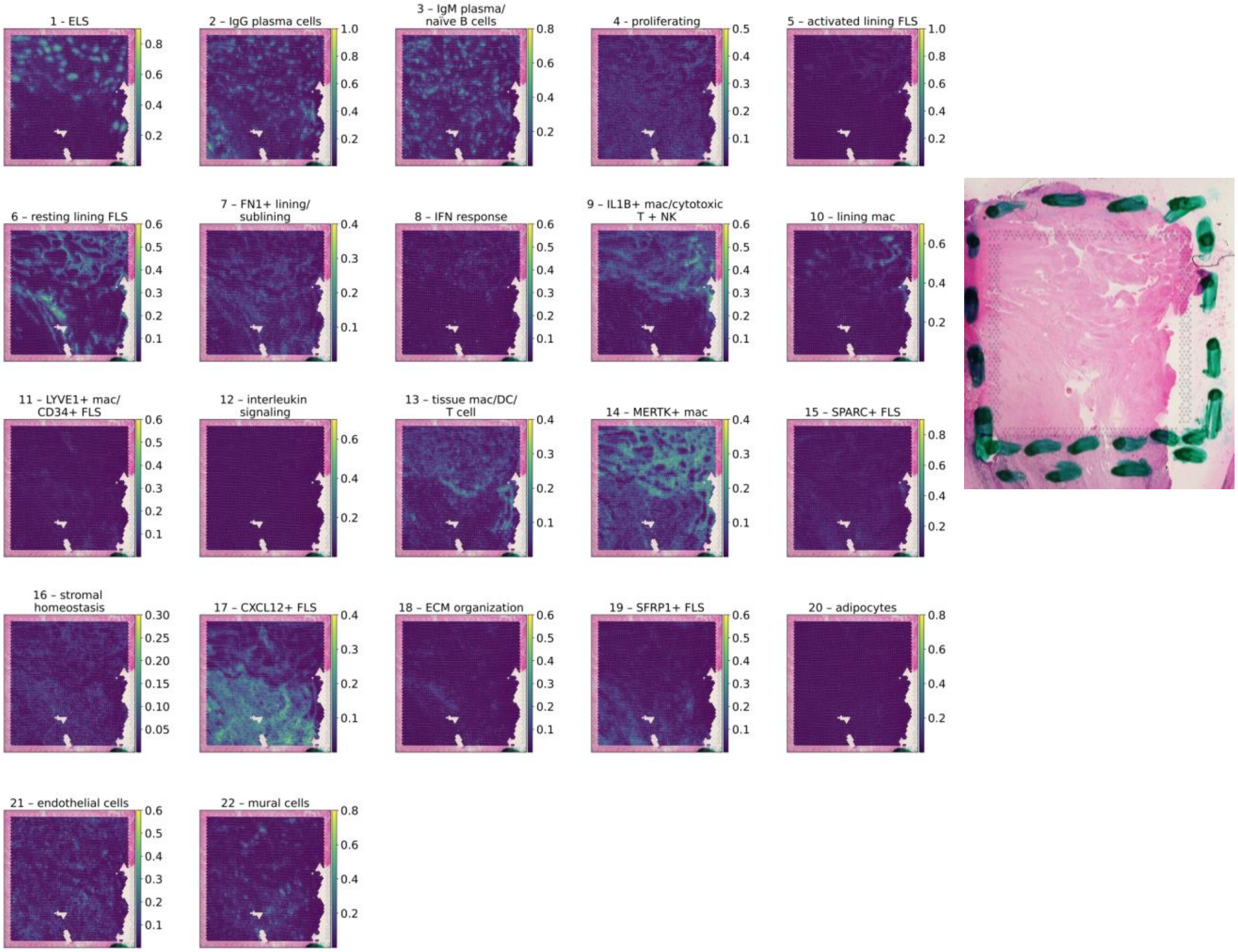
1 - ELS
2 – IgG plasma cells
3 – IgM plasma/ naive B cells
4 – proliferating
5 – activated lining FLS
6 – resting lining FLS
7 – FN1+ lining/ sublining
8 – IFN response
9 – IL1B+ mac/cytotoxic T + NK
10 – lining mac
11 – LYVE1+ mac/ CD34+ FLS
12 – interleukin signaling
13 – tissue mac/DC/ T cell
14 – MERTK+ mac
15 – SPARC+ FLS
16 – stromal homeostasis
17 – CXCL12+ FLS
18 – ECM organization
19 – SFRP1+ FLS
20 – adipocytes
21 – endothelial cells
22 – mural cells

G

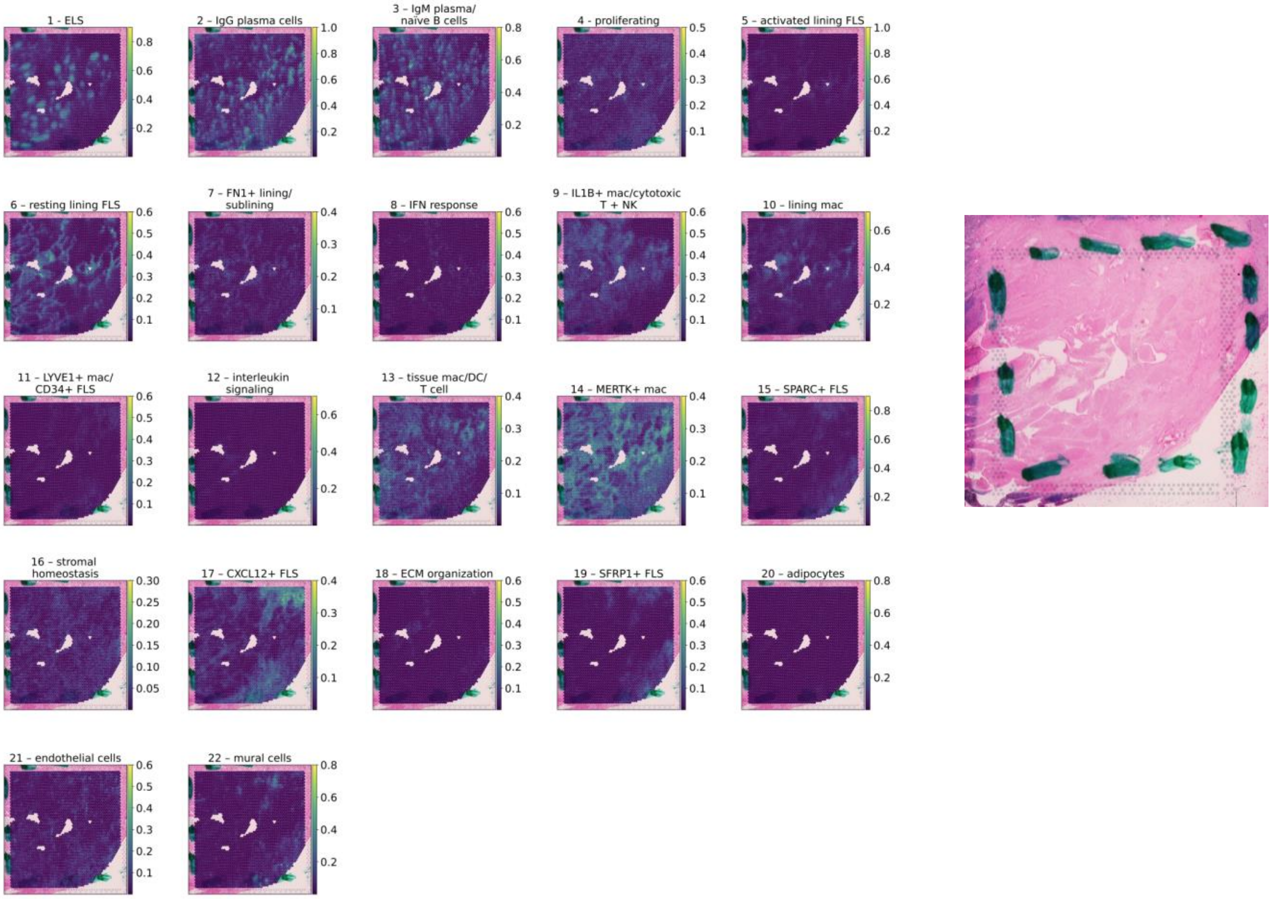


H

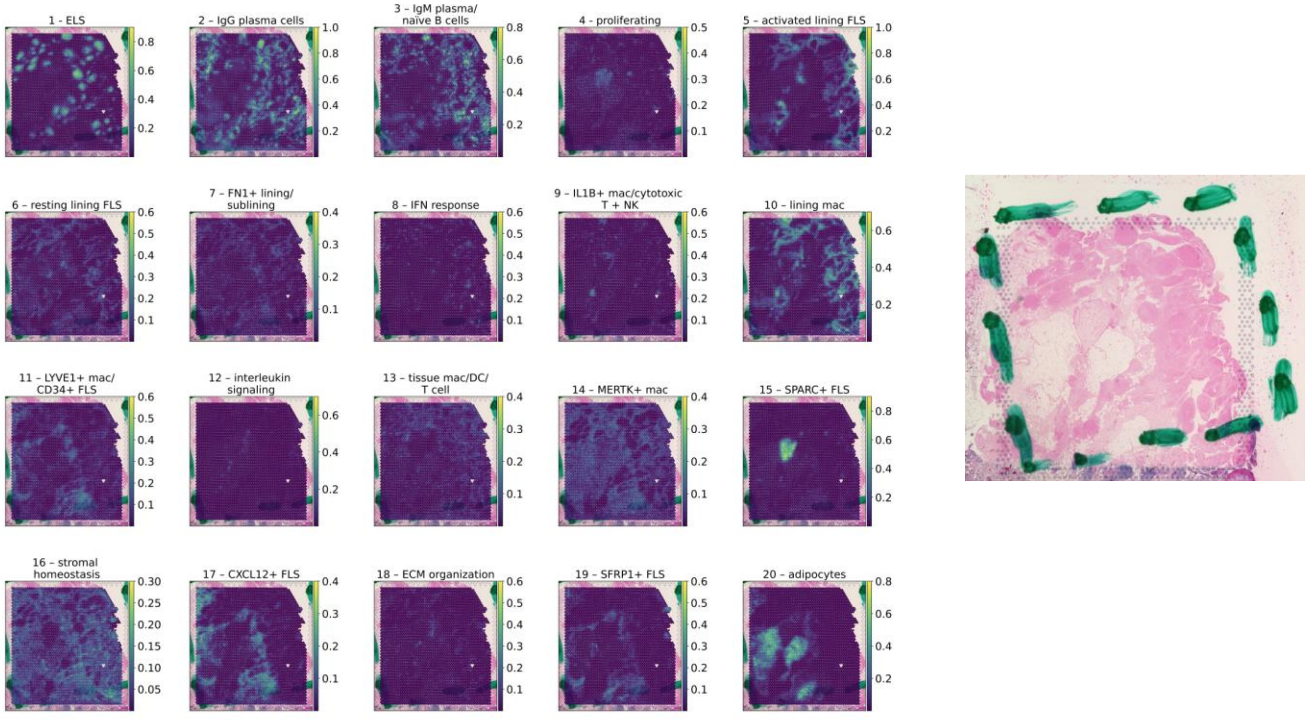


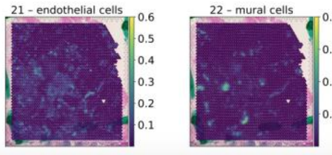


**Supplementary Figure 4:** Spatial distribution of each topic in each tissue section. Posterior probabilities of each topic in each RNA capture spot overlaid onto the corresponding H&E (shown on right for each tissue section): **(A)** RA1, **(B)** RA2A, **(C)** RA2B, **(D)** RA3, **(E)** RA4, **(F)** RA5A, **(G)** RA5B, **(H)** RA6.

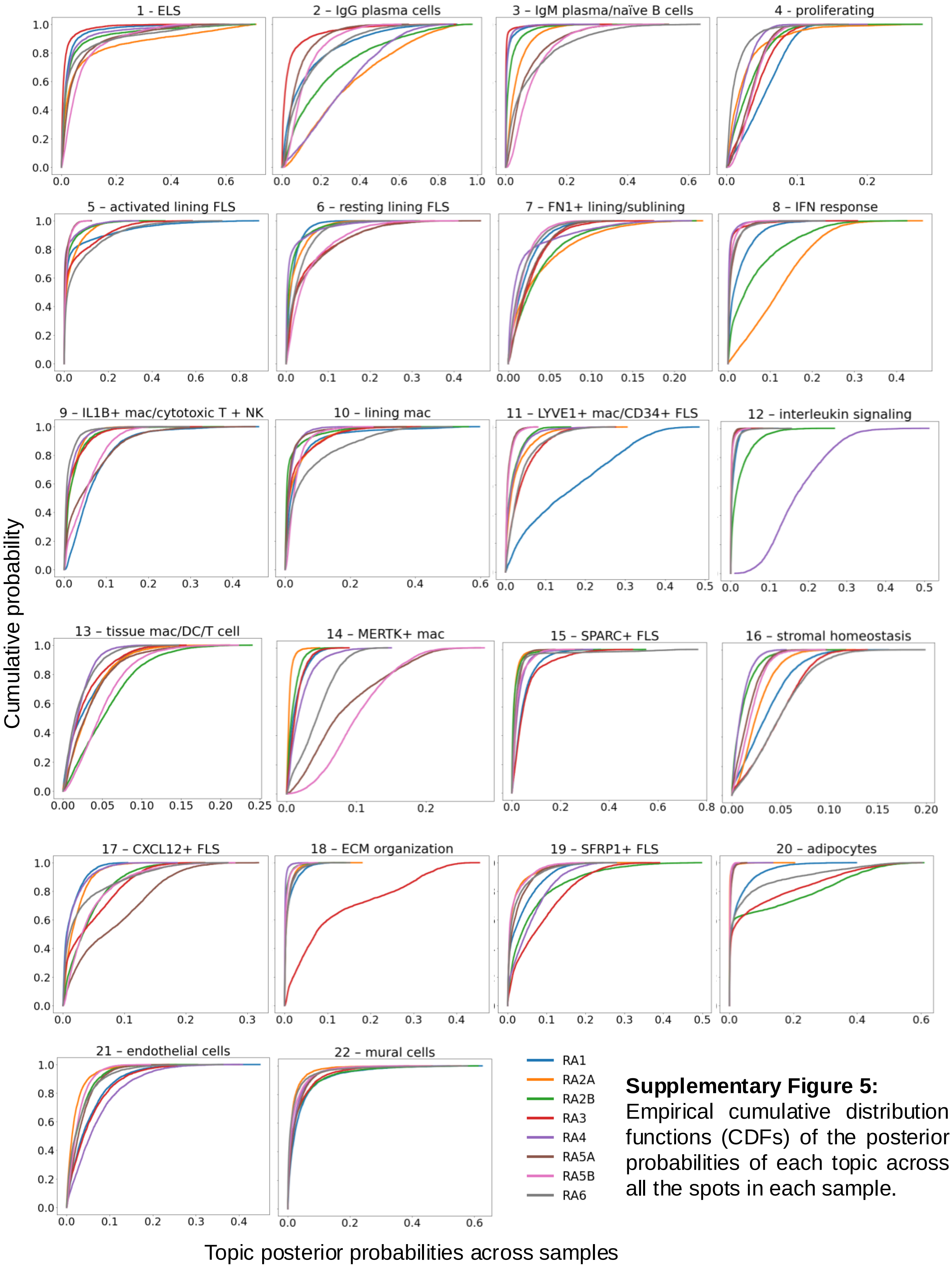


**Supplementary Figure 5:** Empirical cumulative distribution functions (CDFs) of the posterior probabilities of each topic across all the spots in each sample.

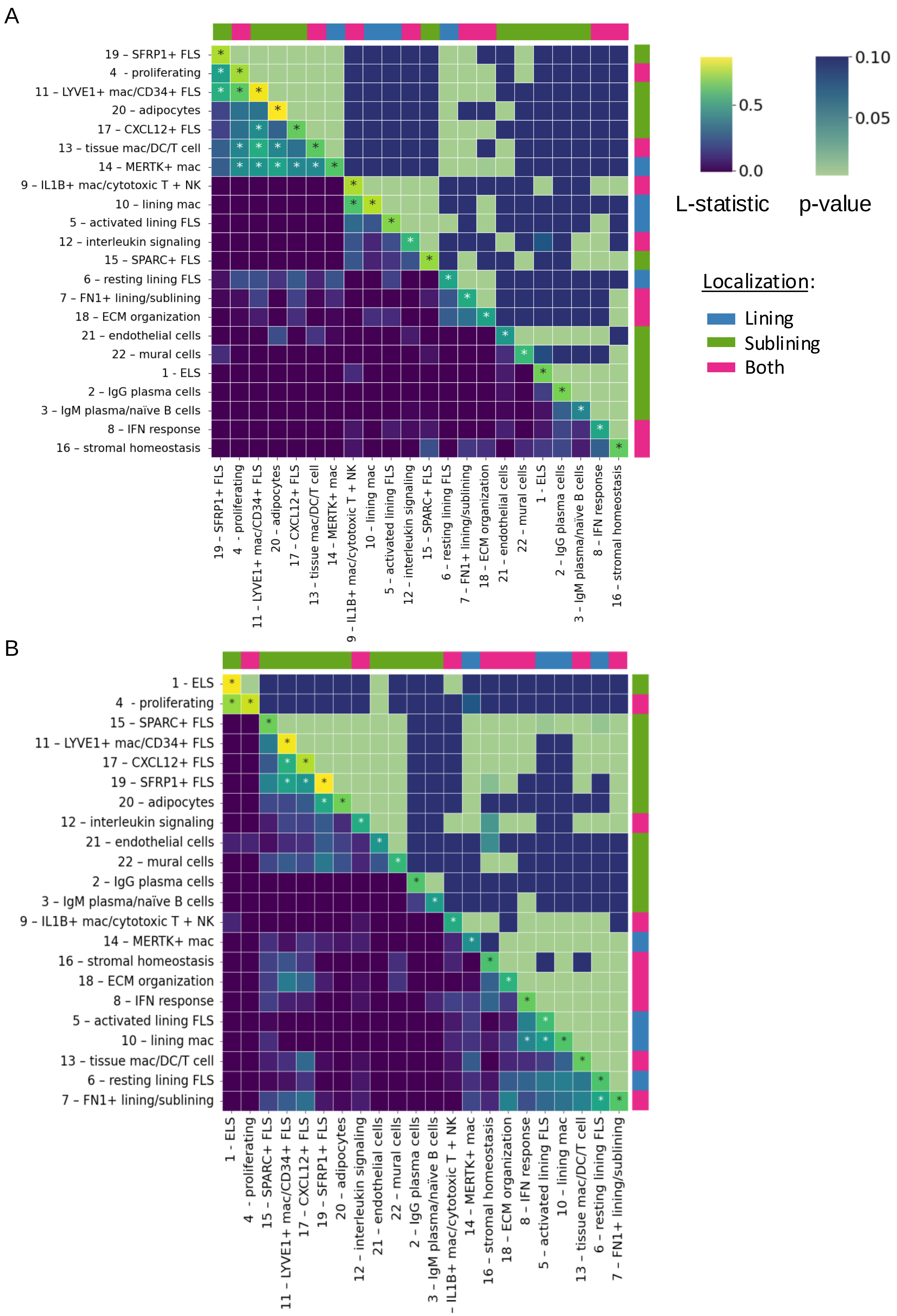

A
19 – SFRP1+ FLS
4 - proliferating
11 – LYVE1+ mac/CD34+ FLS
20 – adipocytes
17 – CXCL12+ FLS
13 – tissue mac/DC/T cell
14 – MERTK+ mac
9 – IL1B+ mac/cytotoxic T + NK
10 – lining mac
5 – activated lining FLS
12 – interleukin signaling
15 – SPARC+ FLS
6 – resting lining FLS
7 – FN1+ lining/sublining
18 – ECM organization
21 – endothelial cells
22 – mural cells
1 - ELS
2 – IgG plasma cells
3 – IgM plasma/naïve B cells
8 – IFN response
16 – stromal homeostasis
0.0
0.5
0.10
0.05
L-statistic
p-value
Localization:
Lining
Sublining
Both
B
1 - ELS
4 - proliferating
15 – SPARC+ FLS
11 – LYVE1+ mac/CD34+ FLS
17 – CXCL12+ FLS
19 – SFRP1+ FLS
20 – adipocytes
12 – interleukin signaling
21 – endothelial cells
22 – mural cells
2 – IgG plasma cells
3 – IgM plasma/naïve B cells
9 – IL1B+ mac/cytotoxic T + NK
14 – MERTK+ mac
16 – stromal homeostasis
18 – ECM organization
8 – IFN response
5 – activated lining FLS
10 – lining mac
13 – tissue mac/DC/T cell
6 – resting lining FLS
7 – FN1+ lining/sublining

C

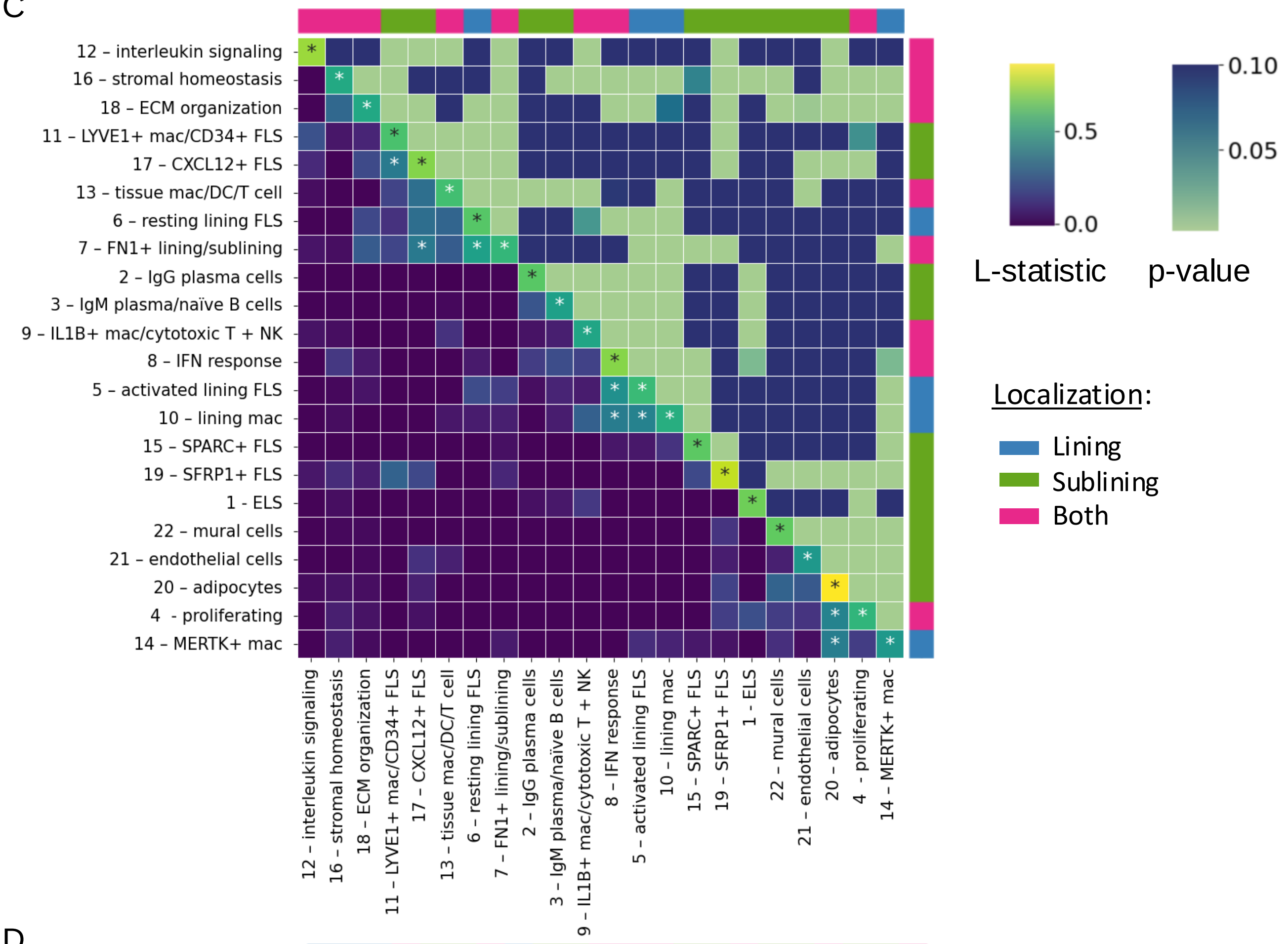

12 – interleukin signaling
16 – stromal homeostasis
18 – ECM organization
11 – LYVE1+ mac/CD34+ FLS
17 – CXCL12+ FLS
13 – tissue mac/DC/T cell
6 – resting lining FLS
7 – FN1+ lining/sublining
2 – IgG plasma cells
3 – IgM plasma/naïve B cells
9 – IL1B+ mac/cytotoxic T + NK
8 – IFN response
5 – activated lining FLS
10 – lining mac
15 – SPARC+ FLS
19 – SFRP1+ FLS
1 - ELS
22 – mural cells
21 – endothelial cells
20 – adipocytes
4 - proliferating
14 – MERTK+ mac
0.0
0.5
0.05
0.10
L-statistic
p-value
Localization:
Lining
Sublining
Both


D

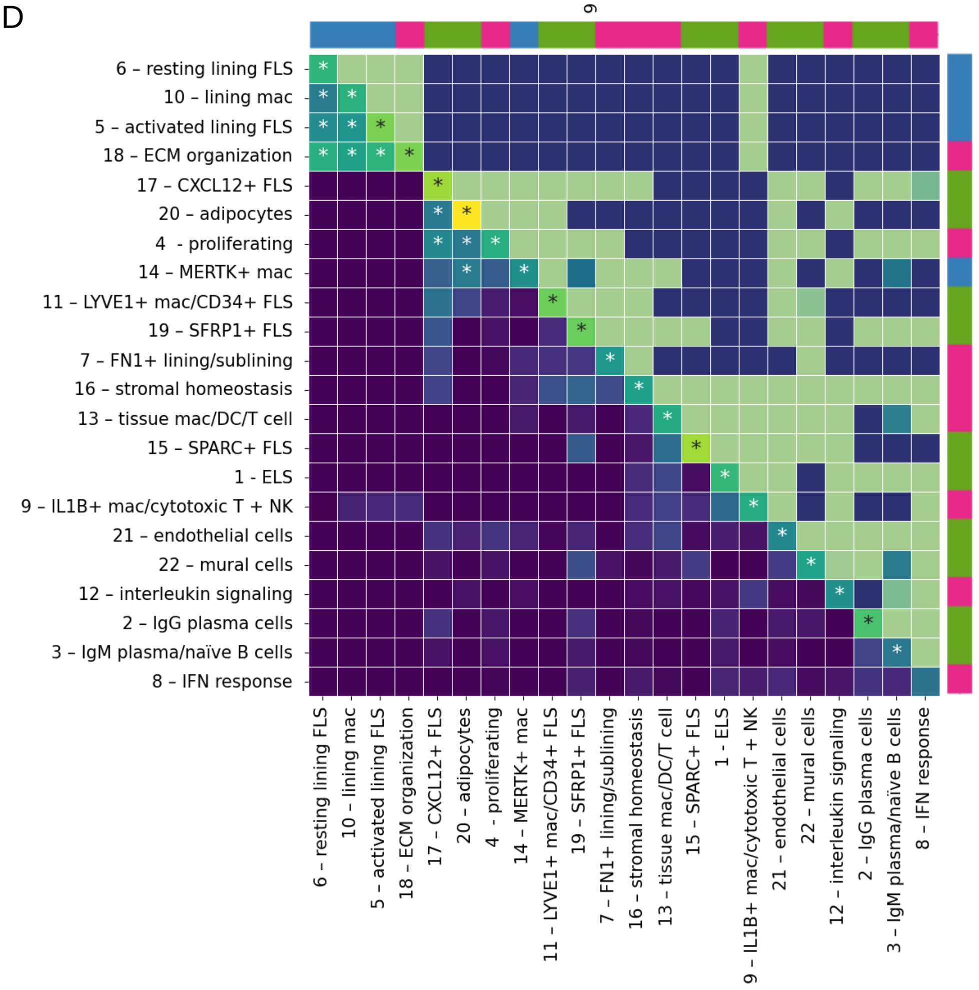

6 – resting lining FLS
10 – lining mac
5 – activated lining FLS
18 – ECM organization
17 – CXCL12+ FLS
20 – adipocytes
4 - proliferating
14 – MERTK+ mac
11 – LYVE1+ mac/CD34+ FLS
19 – SFRP1+ FLS
7 – FN1+ lining/sublining
16 – stromal homeostasis
13 – tissue mac/DC/T cell
15 – SPARC+ FLS
1 - ELS
9 – IL1B+ mac/cytotoxic T + NK
21 – endothelial cells
22 – mural cells
12 – interleukin signaling
2 – IgG plasma cells
3 – IgM plasma/naïve B cells
8 – IFN response

E

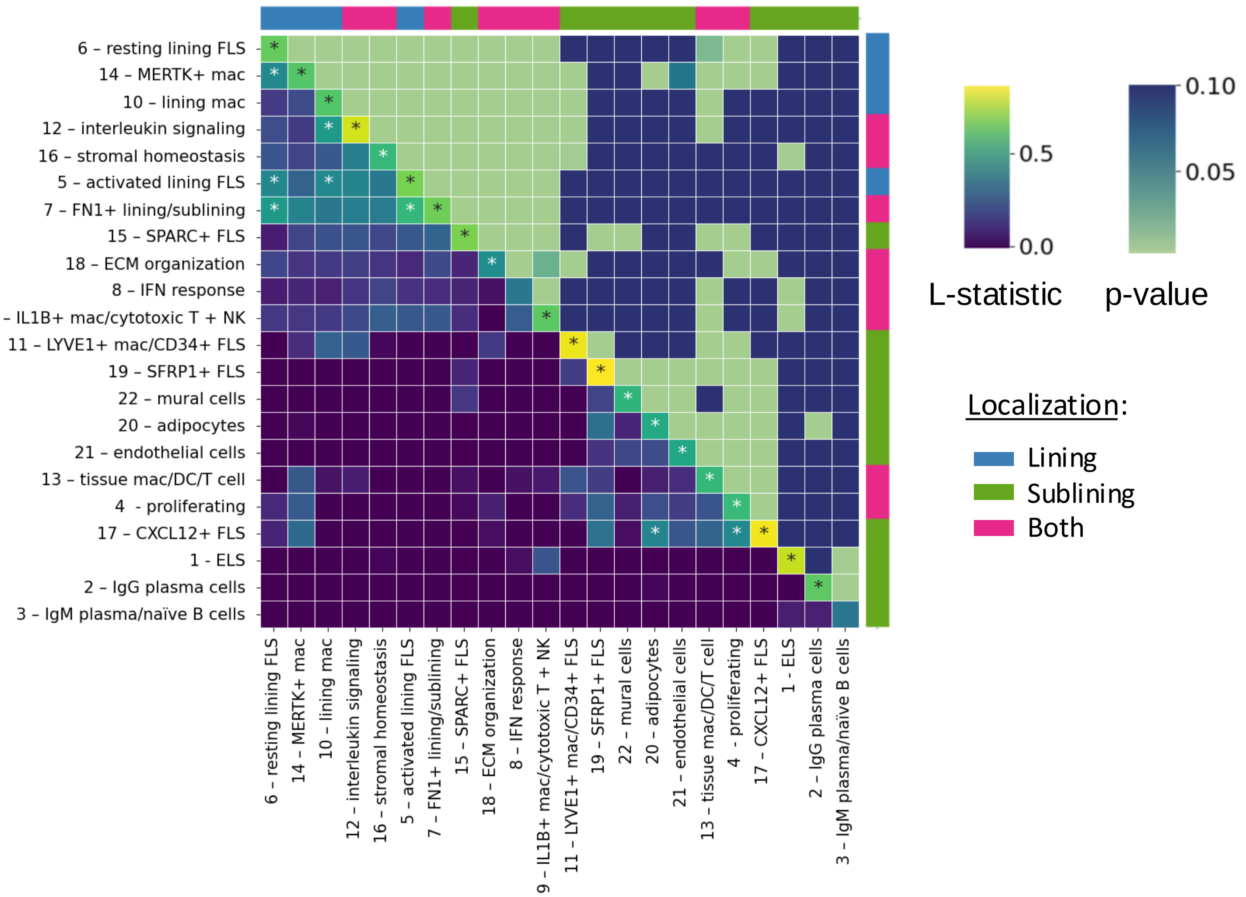
6 – resting lining FLS
14 – MERTK+ mac
10 – lining mac
12 – interleukin signaling
16 – stromal homeostasis
5 – activated lining FLS
7 – FN1+ lining/sublining
15 – SPARC+ FLS
18 – ECM organization
8 – IFN response
9 – IL1B+ mac/cytotoxic T + NK
11 – LYVE1+ mac/CD34+ FLS
19 – SFRP1+ FLS
22 – mural cells
20 – adipocytes
21 – endothelial cells
13 – tissue mac/DC/T cell
4 - proliferating
17 – CXCL12+ FLS
1 - ELS
2 – IgG plasma cells
3 – IgM plasma/naïve B cells
0.0
0.5
L-statistic
0.10
0.05
p-value
Localization:
Lining
Sublining
Both

F

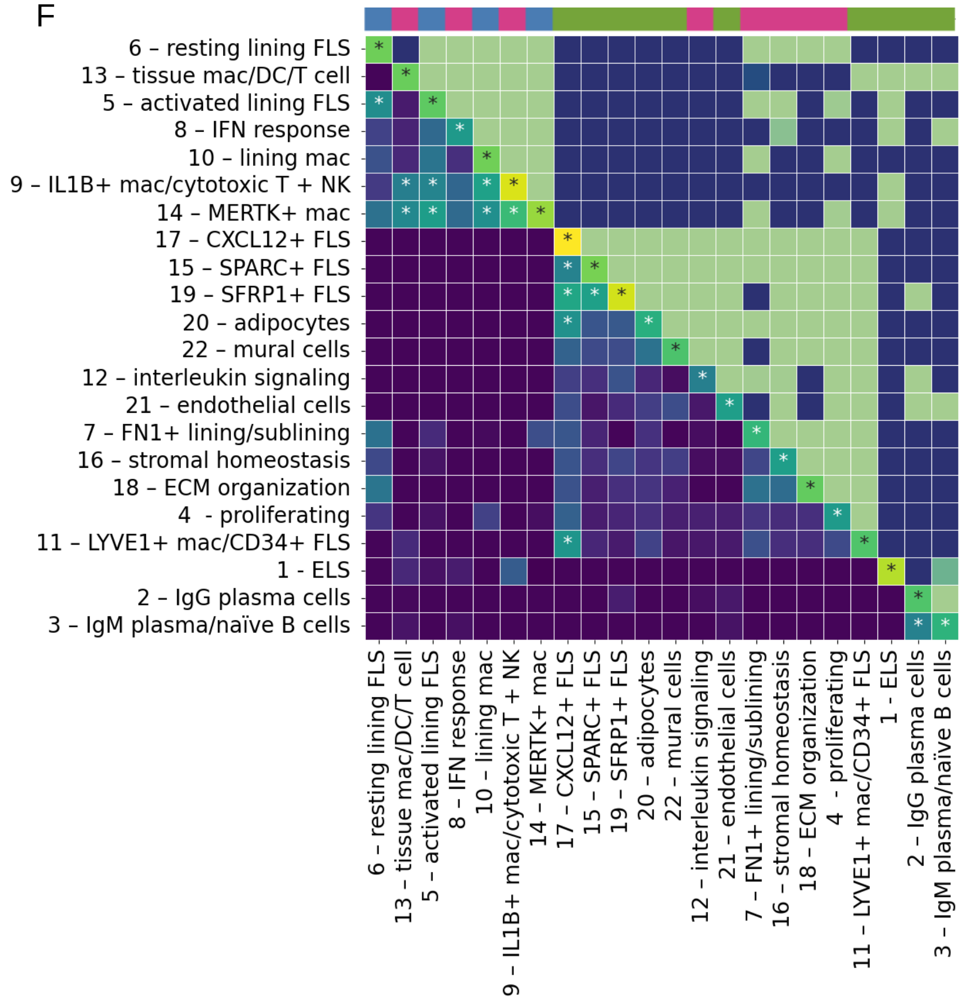
6 – resting lining FLS
13 – tissue mac/DC/T cell
5 – activated lining FLS
8 – IFN response
10 – lining mac
9 – IL1B+ mac/cytotoxic T + NK
14 – MERTK+ mac
17 – CXCL12+ FLS
15 – SPARC+ FLS
19 – SFRP1+ FLS
20 – adipocytes
22 – mural cells
12 – interleukin signaling
21 – endothelial cells
7 – FN1+ lining/sublining
16 – stromal homeostasis
18 – ECM organization
4 - proliferating
11 – LYVE1+ mac/CD34+ FLS
1 - ELS
2 – IgG plasma cells
3 – IgM plasma/naïve B cells

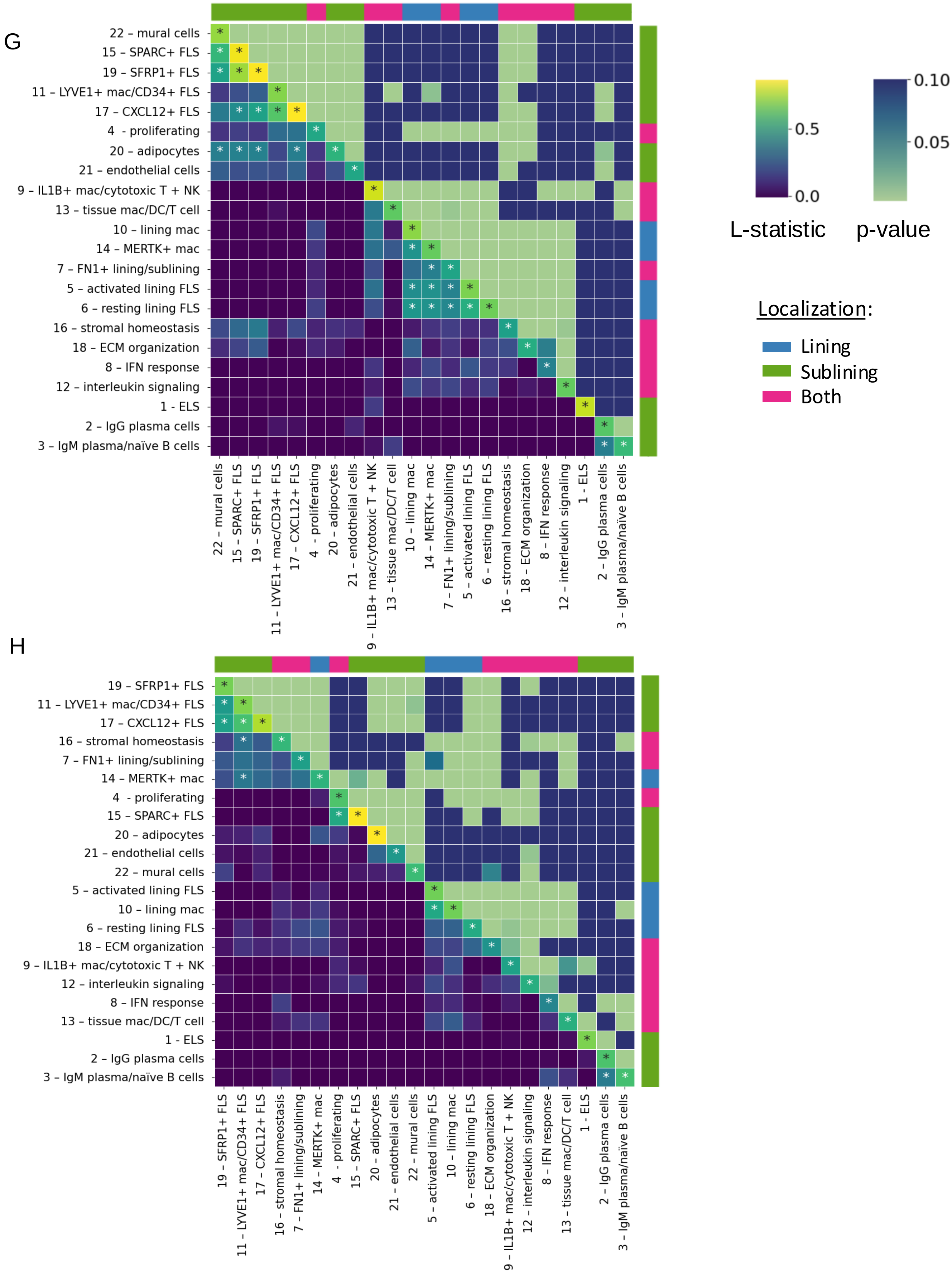


**Supplementary Figure 6:** Statistically significant (Lee's L statistic, adjusted p < 0.05) spatial topic correlations in each sample: **(A)** RA1, **(B)** RA2A, **(C)** RA2B, **(D)** RA3, **(E)** RA4, **(F)** RA5A, **(G)** RA5B, **(H)** RA6. Lower half of heatmap and diagonal: L statistic for each pair of topics. Upper half: p-value for the Lee's L test for that topic pair. Asterisks (*): Lee's L > 0.35 (strong correlation). Outer labels indicate synovial localization of the given topic (lining, sublining, both).

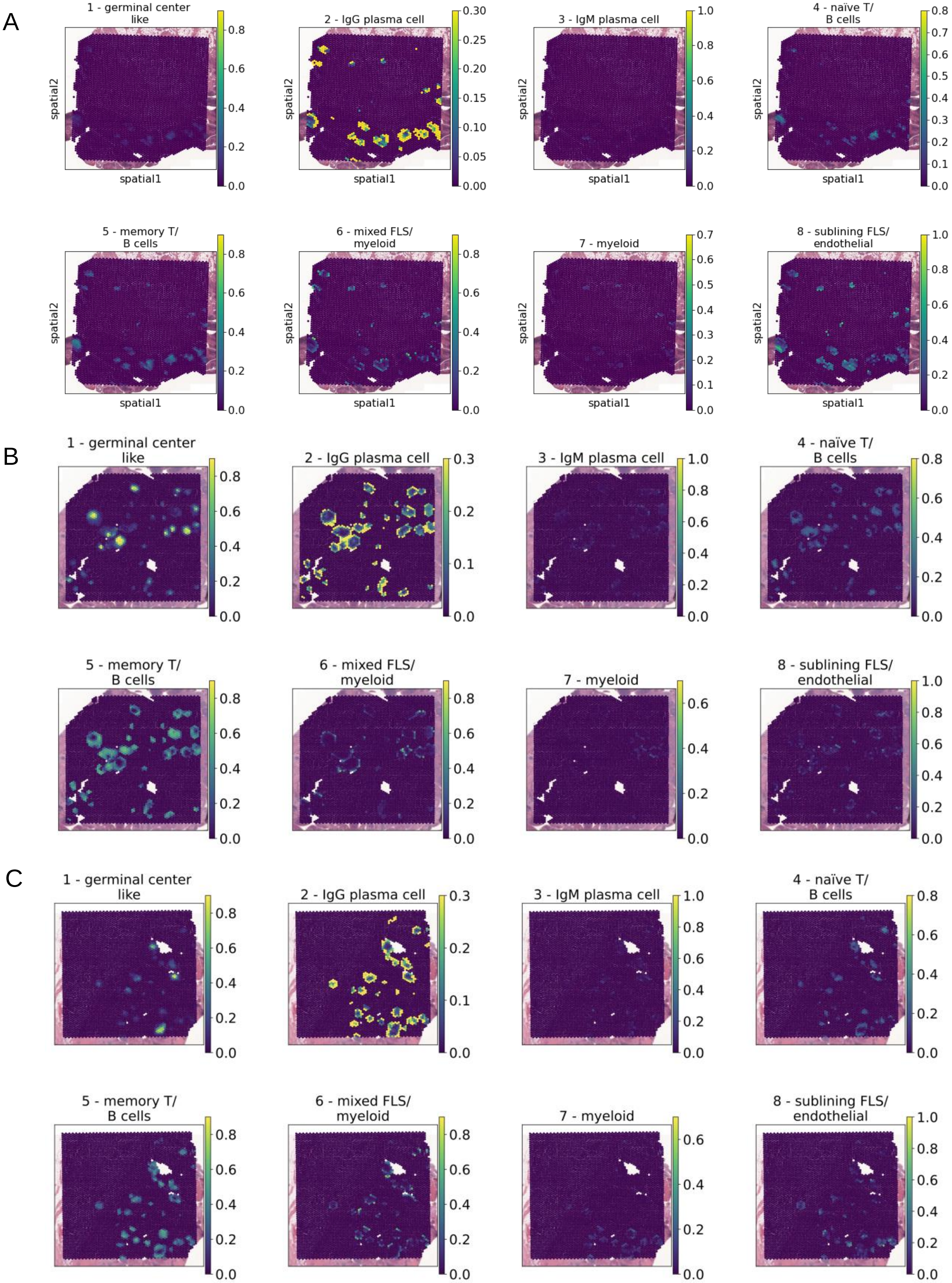
A
1 - germinal center like
2 - IgG plasma cell
3 - IgM plasma cell
4 - naïve T/ B cells
5 - memory T/ B cells
6 - mixed FLS/ myeloid
7 - myeloid
8 - sublining FLS/ endothelial
spatial1
spatial2
B
1 - germinal center like
2 - IgG plasma cell
3 - IgM plasma cell
4 - naïve T/ B cells
5 - memory T/ B cells
6 - mixed FLS/ myeloid
7 - myeloid
8 - sublining FLS/ endothelial
C
1 - germinal center like
2 - IgG plasma cell
3 - IgM plasma cell
4 - naïve T/ B cells
5 - memory T/ B cells
6 - mixed FLS/ myeloid
7 - myeloid
8 - sublining FLS/ endothelial

D

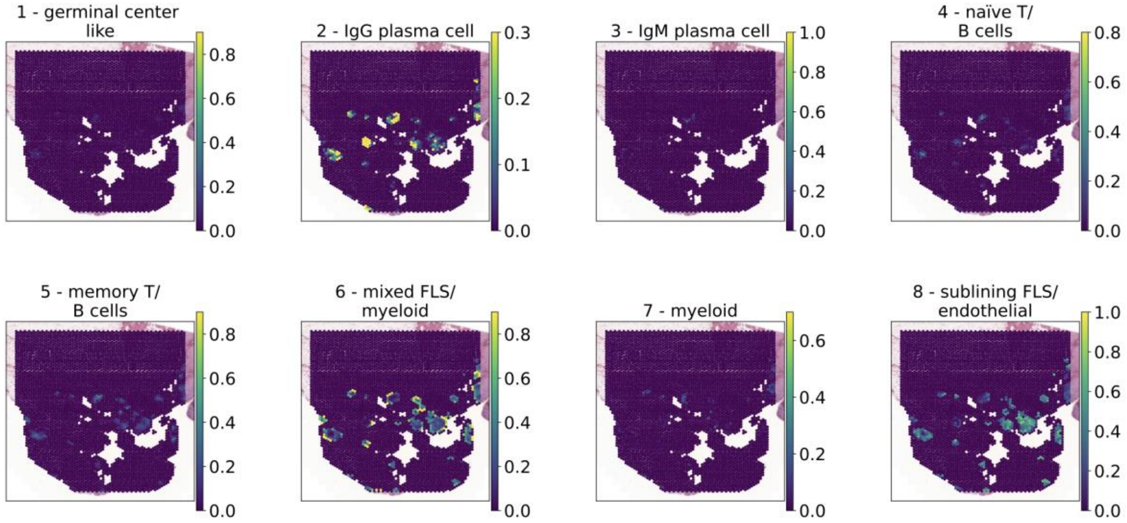
1 - germinal center like
2 - IgG plasma cell
3 - IgM plasma cell
4 - naïve T/ B cells
5 - memory T/ B cells
6 - mixed FLS/ myeloid
7 - myeloid
8 - sublining FLS/ endothelial

E

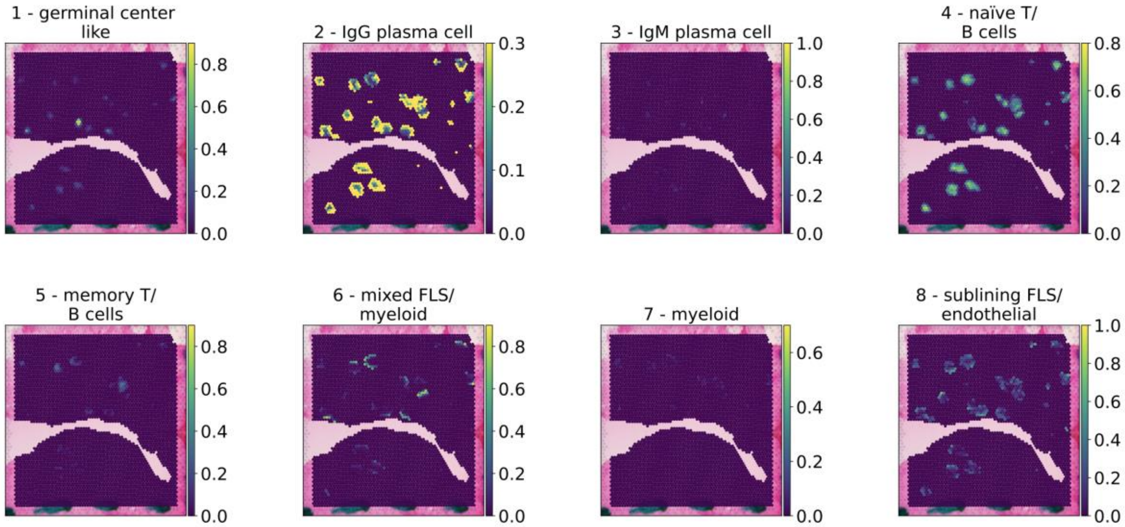
1 - germinal center like
2 - IgG plasma cell
3 - IgM plasma cell
4 - naïve T/ B cells
5 - memory T/ B cells
6 - mixed FLS/ myeloid
7 - myeloid
8 - sublining FLS/ endothelial

F

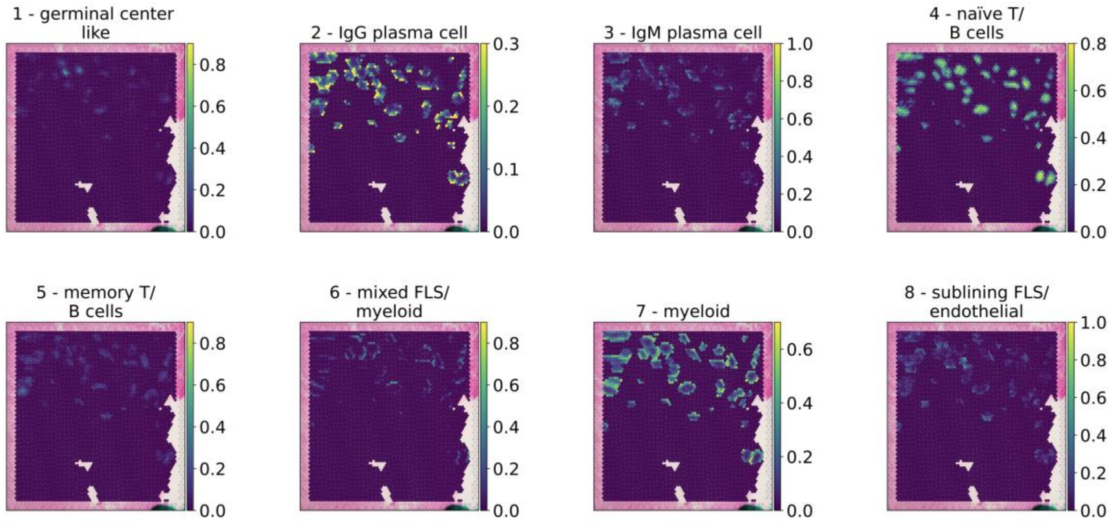
1 - germinal center like
2 - IgG plasma cell
3 - IgM plasma cell
4 - naïve T/ B cells
5 - memory T/ B cells
6 - mixed FLS/ myeloid
7 - myeloid
8 - sublining FLS/ endothelial

G

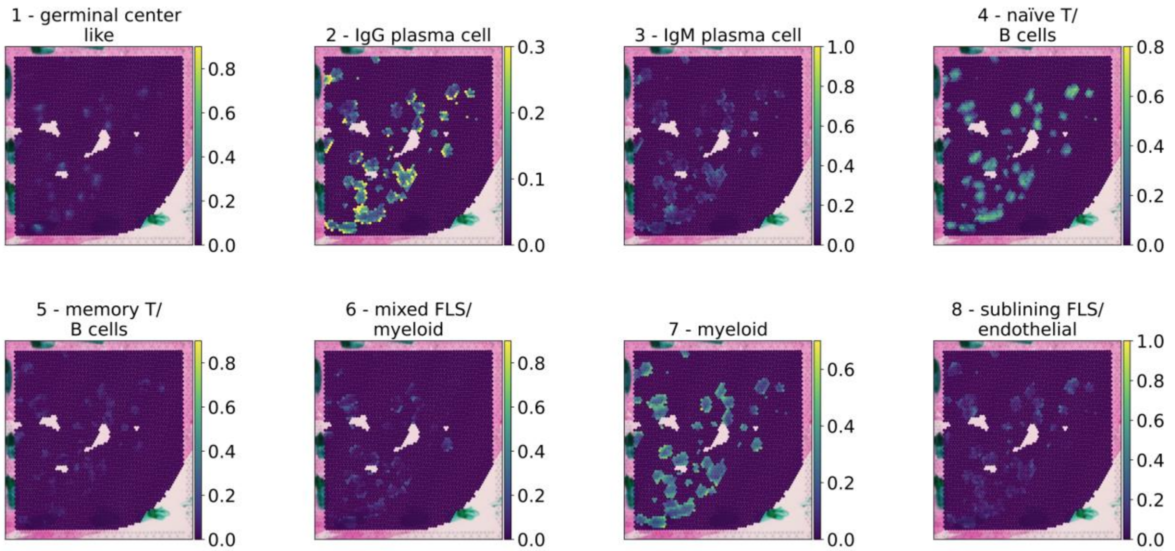


H

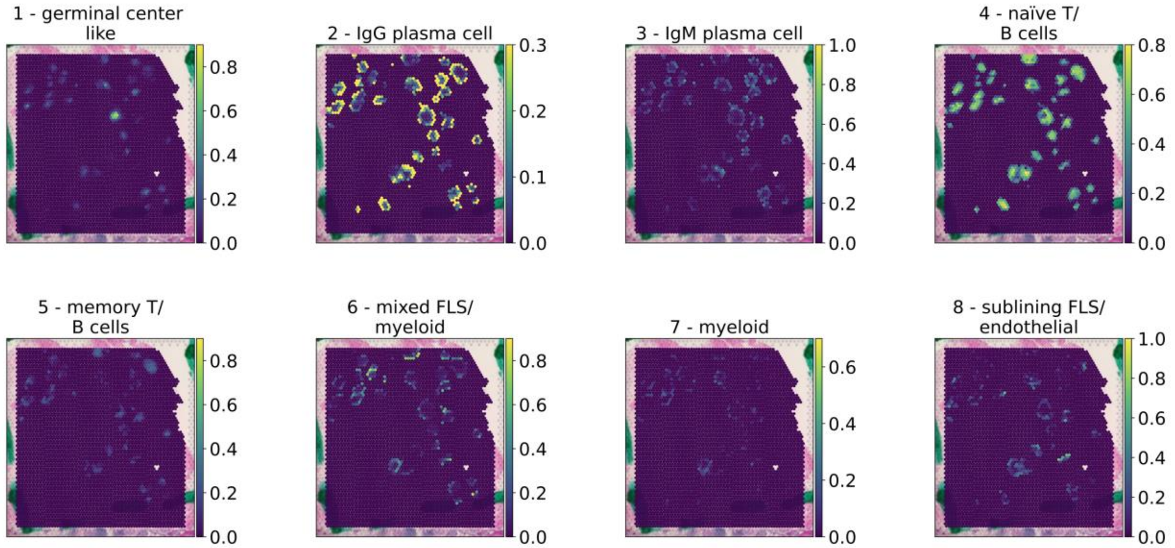


**Supplementary Figure 7:** Spatial distribution of each ELS topic in each tissue section. Posterior probabilities of each topic in each RNA capture spot overlaid onto the corresponding H&E: **(A)** RA1, **(B)** RA2A, **(C)** RA2B, **(D)** RA3, **(E)** RA4, **(F)** RA5A, **(G)** RA5B, **(H)** RA6.

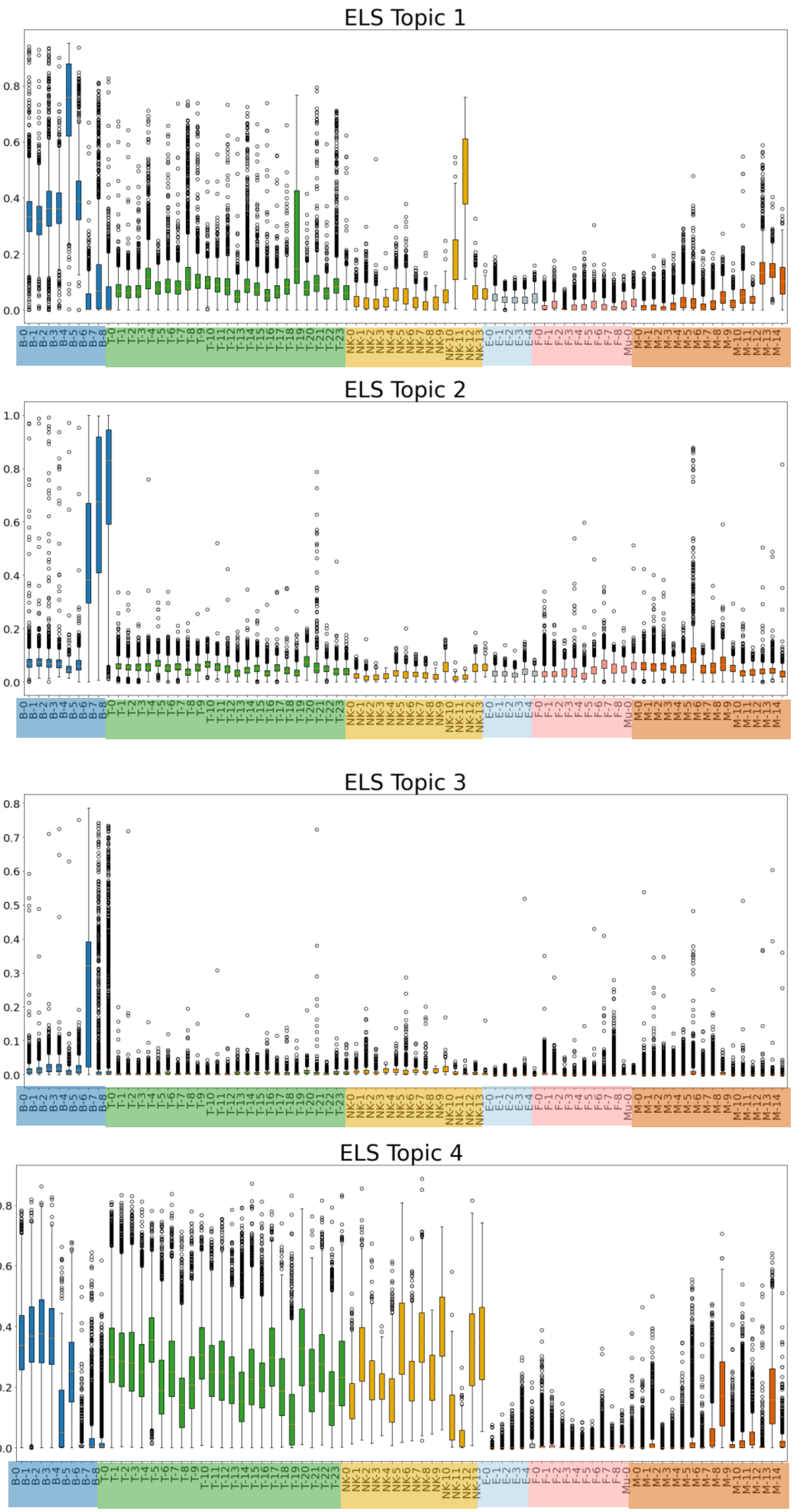


| Symbol | Cluster |
| --- | --- |
| B-0 | CD24+CD27+CD11b+ switched mem |
| B-1 | CD24++CD27+IgM+ unswitched mem |
| B-2 | IgM+IgD+TCL1A+ naive |
| B-3 | IgM+IgD+CD1c+ MZ-like |
| B-4 | AICDA+BCL6+ GC-like |
| B-5 | CD11c+LAMP1+ ABC |
| B-6 | IgM+ plasma |
| B-7 | HLA-DR+IgG+ plasmablast |
| B-8 | IgG1+IgG3+ plasma |
| T-0 | CD4+ IL7R+ memory |
| T-1 | CD4+ CD161+ memory |
| T-2 | CD4+ IL7R+CCR5+ memory |
| T-3 | CD4+ Tfh/Tph |
| T-4 | CD4+ naive |
| T-5 | CD4+ GZMK+ memory |
| T-6 | CD4+ memory |
| T-7 | CD4+ Tph |
| T-8 | CD4+ CD25-high Treg |
| T-9 | CD4+ CD25-low Treg |
| T-10 | CD4+ OX40+NR3C1+ |
| T-11 | CD4+ CD146+ memory |
| T-12 | CD4+ GNLY+ |
| T-13 | CD8+ GZMK/B+ memory |
| T-14 | CD8+ GZMK+ memory |
| T-15 | CD8+ GZMB+/TEMRA |
| T-16 | CD8+ CD45ROlow/naive |
| T-17 | CD8+ activated/NK-like |
| T-18 | Proliferating |
| T-19 | MT-high (low quality) |
| T-20 | CD38+ |
| T-21 | Innate-like |
| T-22 | Vdelta1 |
| T-23 | Vdelta2 |
| NK-0 | CD56dim CD16+ IFNG- |
| NK-1 | CD56dim CD16+ IFNG+CD160+ |
| NK-2 | CD56dim CD16+ IFNG+CD160- |
| NK-3 | CD56dim CD16+ GZMB- |
| NK-4 | CD56bright CD16- GZMA+CD160+ |
| NK-5 | CD56bright CD16- GZMA+CD69+ |
| NK-6 | CD56bright CD16- GNLY+ |
| NK-7 | CD56bright CD16- GNLY+CD69+ |
| NK-8 | CD56bright CD16- IFN response |
| NK-9 | MT-high |
| NK-10 | PCNA+ Proliferating |
| NK-11 | MKI67+ Proliferating |
| NK-12 | IL7R+ ILC |
| NK-13 | IL7R+CD161+ ILC |
| E-0 | SPARC+ capillary |
| E-1 | LIFR+ venular |
| E-2 | ICAM1+ venular |
| E-3 | NOTCH4+ arteriolar |
| E-4 | Lymphatic |
| F-0 | PRG4+ CLIC5+ lining |
| F-1 | PRG4+ lining |
| F-2 | CD34+ sublining |
| F-3 | POSTN+ sublining |
| F-4 | DKK3+ sublining |
| F-5 | CD74-hi sublining |
| F-6 | CXCL12+ SFRP1+ sublining |
| F-7 | NOTCH3+ sublining |
| F-8 | RSPO3+ intermediate |
| Mu-0 | Mural |
| M-0 | MERTK+ SELENOP+ LYVE1+ |
| M-1 | MERTK+ SELENOP+ LYVE1- |
| M-2 | MERTK+ S100A8+ |
| M-3 | MERTK+ HBEGF+ |
| M-4 | SPP1+ |
| M-5 | C1QA+ |
| M-6 | STAT1+ CXCL10+ |
| M-7 | IL1B+ FCN1+ |
| M-8 | PLCG2+ |
| M-9 | DC3 |
| M-10 | DC2 |
| M-11 | DC4 |
| M-12 | DC1 |
| M-13 | pDC |
| M-14 | LAMP3+ |

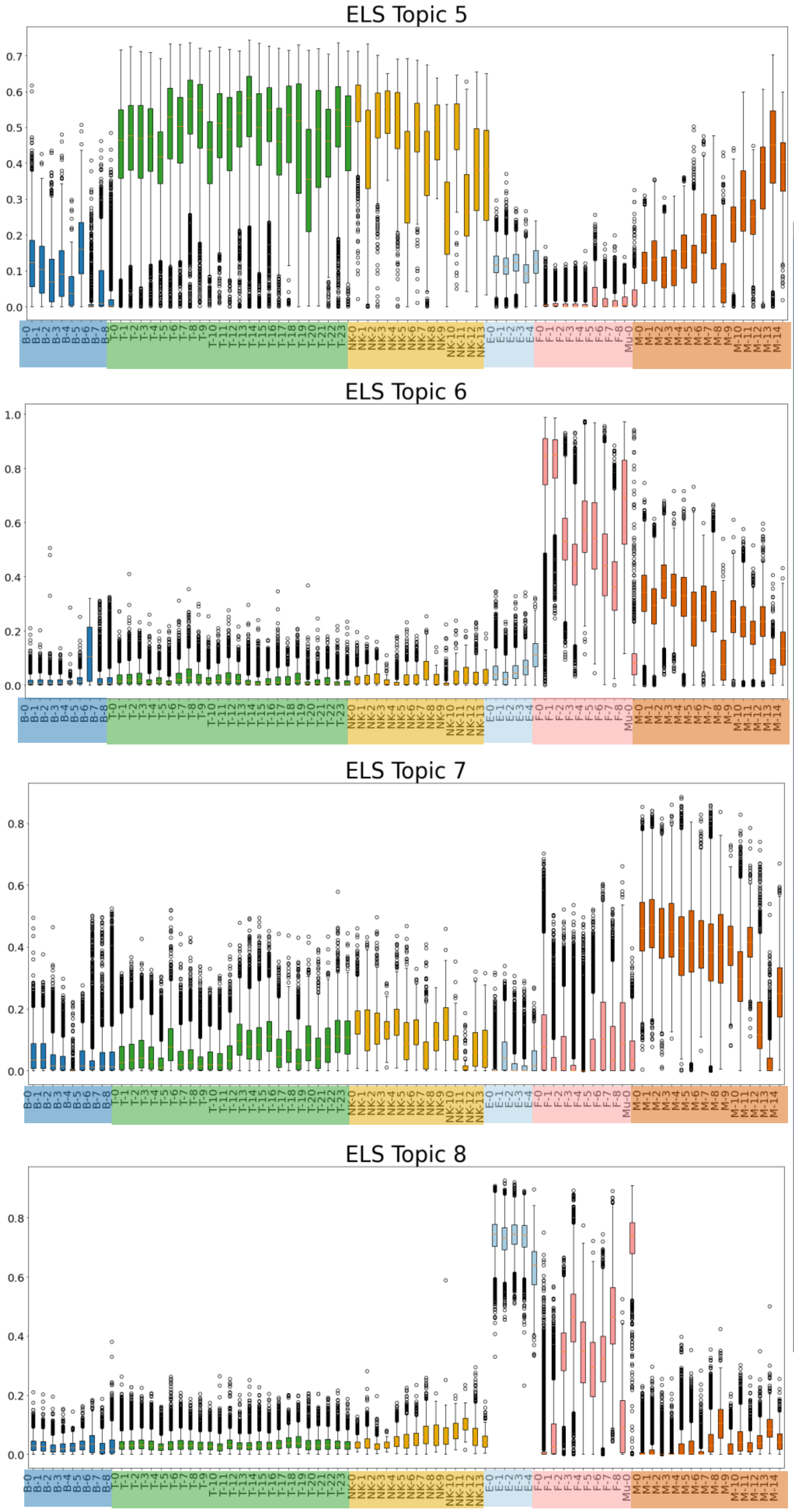


| Symbol | Cluster |
|---|---|
| B-0 | CD24+CD27+CD11b+ switched mem |
| B-1 | CD24++CD27+IgM+ unswitched mem |
| B-2 | IgM+IgD+TCL1A+ naive |
| B-3 | IgM+IgD+CD1c+ MZ-like |
| B-4 | AICDA+BCL6+ GC-like |
| B-5 | CD11c+LAMP1+ ABC |
| B-6 | IgM+ plasma |
| B-7 | HLA-DR+IgG+ plasmablast |
| B-8 | IgG1+IgG3+ plasma |
| T-0 | CD4+ IL7R+ memory |
| T-1 | CD4+ CD161+ memory |
| T-2 | CD4+ IL7R+CCR5+ memory |
| T-3 | CD4+ Tfh/Tph |
| T-4 | CD4+ naive |
| T-5 | CD4+ GZMK+ memory |
| T-6 | CD4+ memory |
| T-7 | CD4+ Tph |
| T-8 | CD4+ CD25-high Treg |
| T-9 | CD4+ CD25-low Treg |
| T-10 | CD4+ OX40+NR3C1+ |
| T-11 | CD4+ CD146+ memory |
| T-12 | CD4+ GNLY+ |
| T-13 | CD8+ GZMK/B+ memory |
| T-14 | CD8+ GZMK+ memory |
| T-15 | CD8+ GZMB+/TEMRA |
| T-16 | CD8+ CD45ROlow/naive |
| T-17 | CD8+ activated/NK-like |
| T-18 | Proliferating |
| T-19 | MT-high (low quality) |
| T-20 | CD38+ |
| T-21 | Innate-like |
| T-22 | Vdelta1 |
| T-23 | Vdelta2 |
| NK-0 | CD56dim CD16+ IFNG- |
| NK-1 | CD56dim CD16+ IFNG+CD160+ |
| NK-2 | CD56dim CD16+ IFNG+CD160- |
| NK-3 | CD56dim CD16+ GZMB- |
| NK-4 | CD56bright CD16- GZMA+CD160+ |
| NK-5 | CD56bright CD16- GZMA+CD69+ |
| NK-6 | CD56bright CD16- GNLY+ |
| NK-7 | CD56bright CD16- GNLY+CD69+ |
| NK-8 | CD56bright CD16- IFN response |
| NK-9 | MT-high |
| NK-10 | PCNA+ Proliferating |
| NK-11 | MKI67+ Proliferating |
| NK-12 | IL7R+ ILC |
| NK-13 | IL7R+CD161+ ILC |
| E-0 | SPARC+ capillary |
| E-1 | LIFR+ venular |
| E-2 | ICAM1+ venular |
| E-3 | NOTCH4+ arteriolar |
| E-4 | Lymphatic |
| F-0 | PRG4+ CLIC5+ lining |
| F-1 | PRG4+ lining |
| F-2 | CD34+ sublining |
| F-3 | POSTN+ sublining |
| F-4 | DKK3+ sublining |
| F-5 | CD74-hi sublining |
| F-6 | CXCL12+ SFRP1+ sublining |
| F-7 | NOTCH3+ sublining |
| F-8 | RSPO3+ intermediate |
| Mu-0 | Mural |
| M-0 | MERTK+ SELENOP+ LYVE1+ |
| M-1 | MERTK+ SELENOP+ LYVE1- |
| M-2 | MERTK+ S100A8+ |
| M-3 | MERTK+ HBEGF+ |
| M-4 | SPP1+ |
| M-5 | C1QA+ |
| M-6 | STAT1+ CXCL10+ |
| M-7 | IL1B+ FCN1+ |
| M-8 | PLCG2+ |
| M-9 | DC3 |
| M-10 | DC2 |
| M-11 | DC4 |
| M-12 | DC1 |
| M-13 | pDC |
| M-14 | LAMP3+ |

**Supplementary Figure 8:** Application of the DeepTopics model trained on the ELS spots to the published scRNA-seq atlas[25] for RA. Each box represents the interquartile range of posterior probabilities for the topic across cells belonging to the cluster.

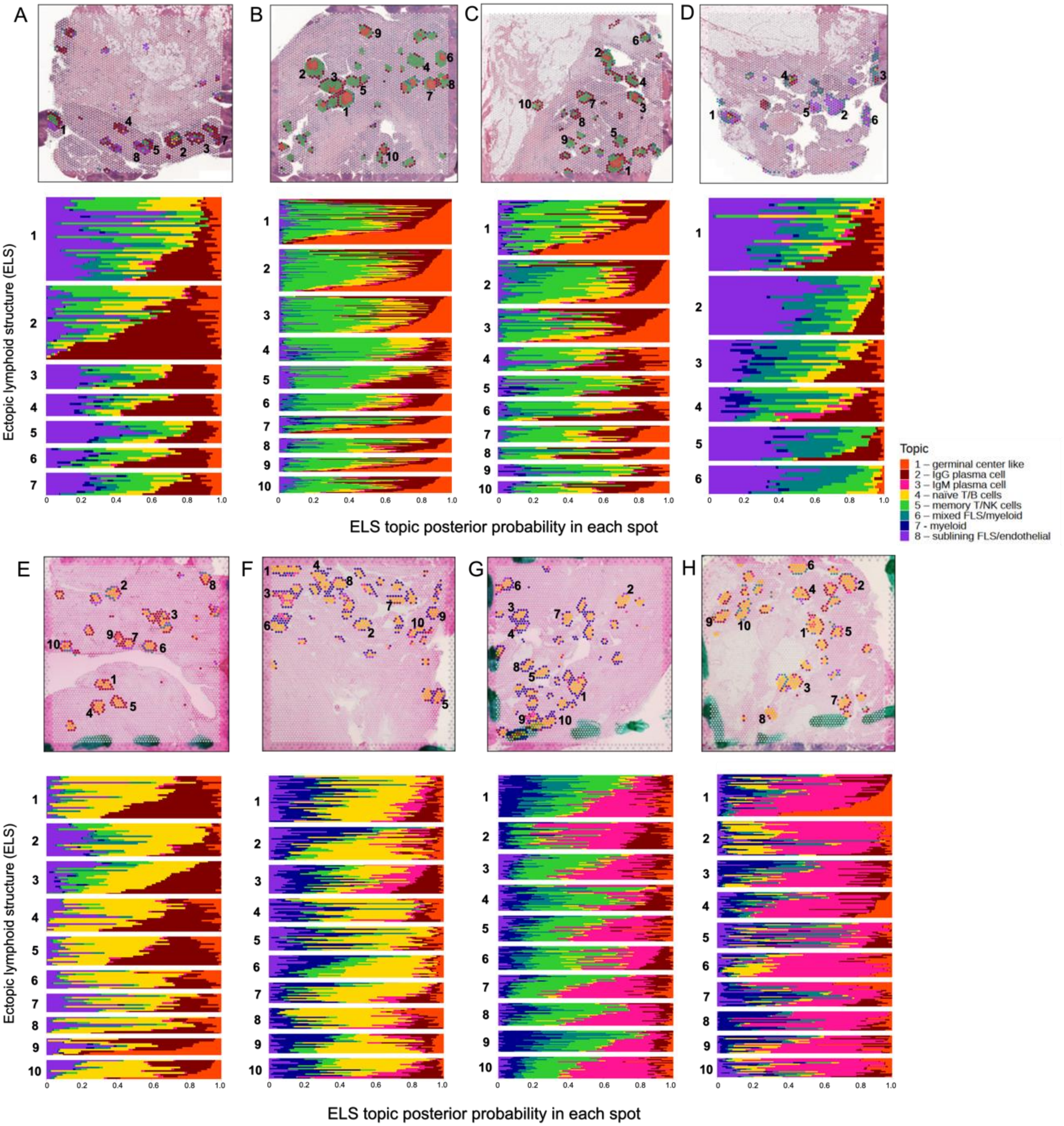


**Supplementary Figure 9:** Cell compositions of individual ELS plotted on each H&E (top), and stacked bar plots of the topic posterior probabilities in each spot within each ELS (bottom), in **(A)** RA1, **(B)** RA2A, **(C)** RA2B, **(D)** RA3, **(E)** RA4, **(F)** RA5A, **(G)** RA5B, **(H)** RA6. In the spatial plots (top), each RNA capture spot in each ELS is colored according to the topic with the maximum posterior probability in that spot. ELS (in both H&E and stacked bar plots) are numbered by size (number of RNA capture spots encompassed by an ELS). The topic colors in the legend correspond to both the spatial plots and the stacked bar plots.

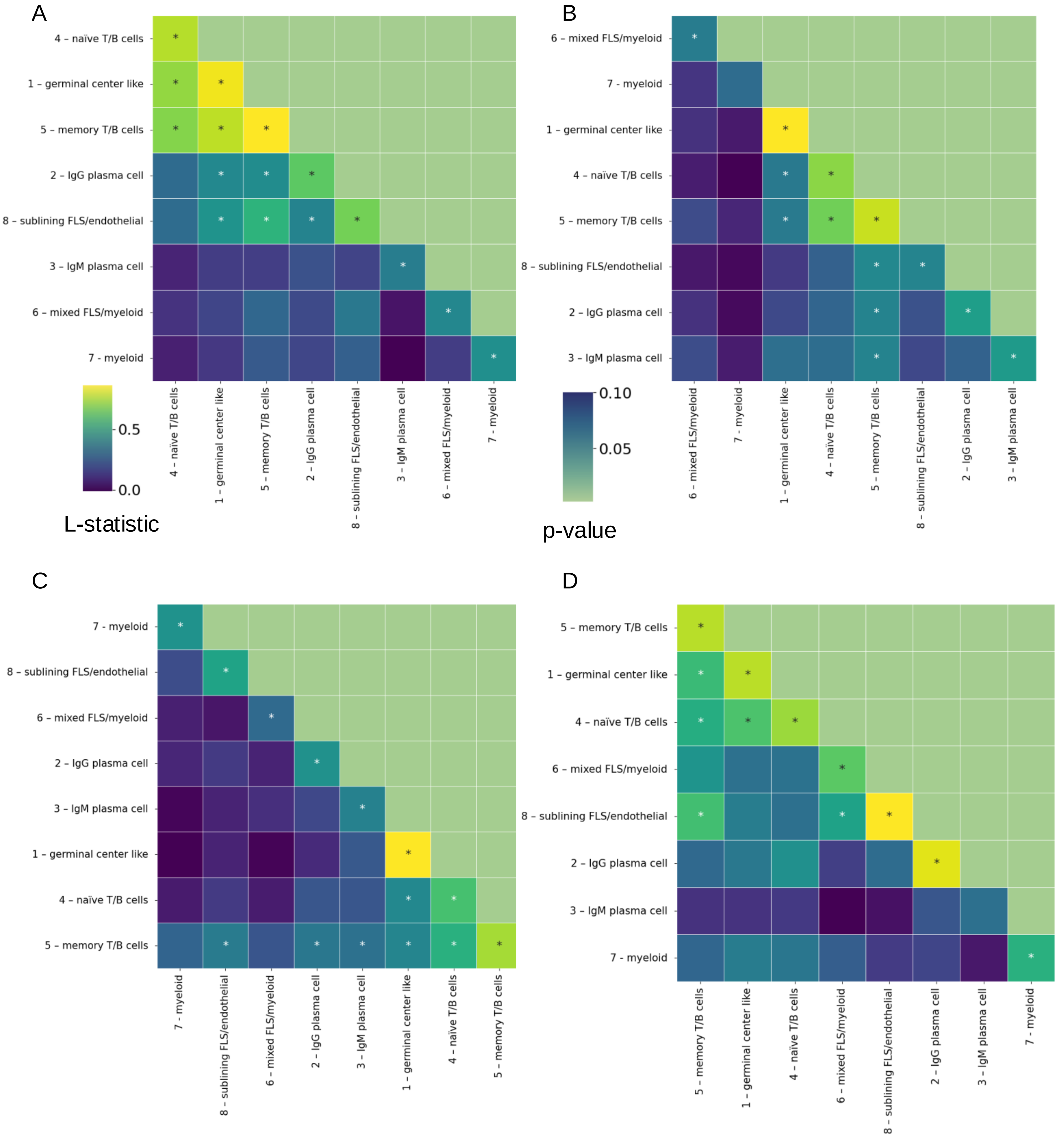

A
4 – naïve T/B cells
1 – germinal center like
5 – memory T/B cells
2 – IgG plasma cell
8 – sublining FLS/endothelial
3 – IgM plasma cell
6 – mixed FLS/myeloid
7 - myeloid
0.5
0.0
L-statistic
B
6 – mixed FLS/myeloid
7 - myeloid
1 – germinal center like
4 – naïve T/B cells
5 – memory T/B cells
8 – sublining FLS/endothelial
2 – IgG plasma cell
3 – IgM plasma cell
0.10
0.05
p-value
C
7 - myeloid
8 – sublining FLS/endothelial
6 – mixed FLS/myeloid
2 – IgG plasma cell
3 – IgM plasma cell
1 – germinal center like
4 – naïve T/B cells
5 – memory T/B cells
D
5 – memory T/B cells
1 – germinal center like
4 – naïve T/B cells
6 – mixed FLS/myeloid
8 – sublining FLS/endothelial
2 – IgG plasma cell
3 – IgM plasma cell
7 - myeloid

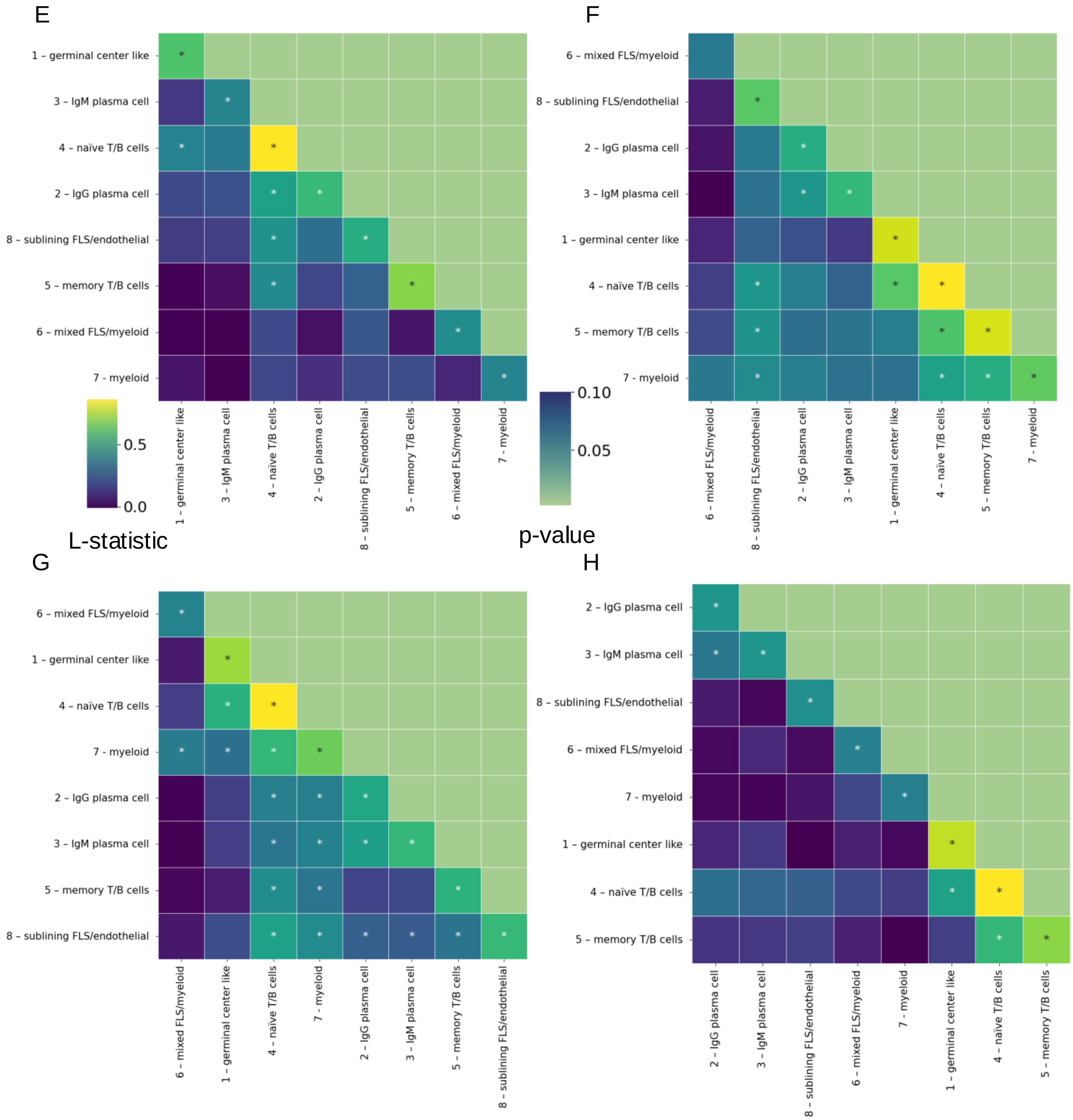


**Supplementary Figure 10:** Statistically significant (Lee's L statistic, adjusted $p < 0.05$) spatial topic correlations in each sample: **(A)** RA1, **(B)** RA2A, **(C)** RA2B, **(D)** RA3, **(E)** RA4, **(F)** RA5A, **(G)** RA5B, **(H)** RA6. Lower half of heatmap and diagonal: L statistic for each pair of topics. Upper half: p-value for the Lee's L test for that topic pair. Asterisks (*): Lee's L > 0.35 (strong correlation).

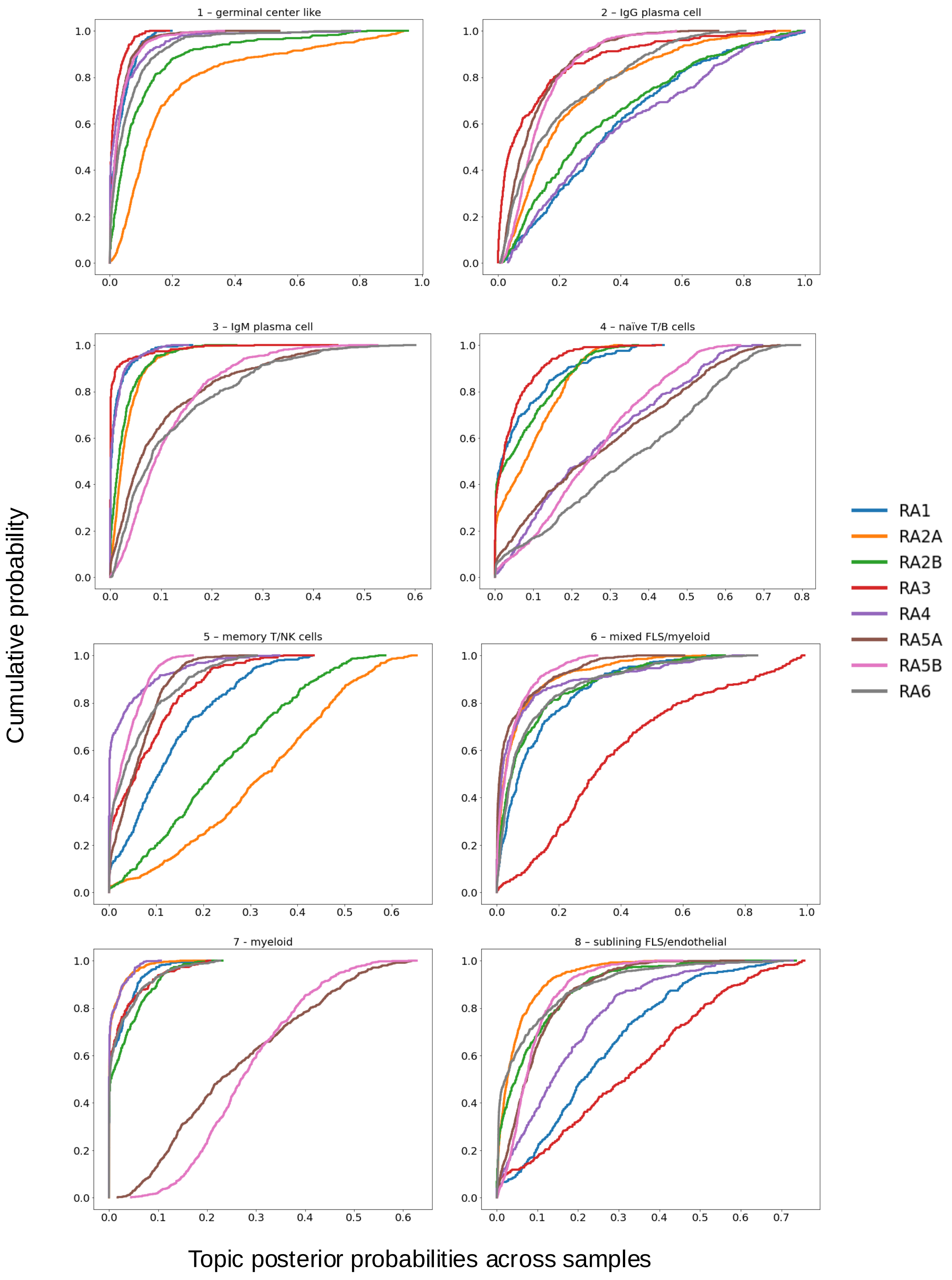


**Supplementary Figure 11:** Empirical cumulative distribution functions (CDFs) of the posterior probabilities of each ELS topic across all the ELS spots in each sample.
